# FlashDeconv reveals resolution horizons in atlas-scale spatial transcriptomics

**DOI:** 10.64898/2025.12.22.696108

**Authors:** Chen Yang, Xianyang Zhang, Jun Chen

## Abstract

Coarsening Visium HD resolution from 8 to 64 ***µ***m can flip cell-type co-localization from negative to positive (***r* = −0.12 → +0.80**), yet many widely used compositional deconvolution workflows require coarsening or sub-sampling at million-bin scale. Here we introduce FlashDeconv, which combines leverage-score importance sampling with sparse spatial regularization to achieve competitive benchmark accuracy while processing 1.6 million bins in 153 seconds on commodity hardware. Systematic multi-resolution Visium HD analysis reveals tissue-specific *resolution horizons*—a sharp 8–16 ***µ***m transition in mouse intestinal epithelium versus gradual degradation in brain cortex—both validated by independent single-cell ground truth. At sub-16 ***µ***m resolution, FlashDeconv enables sequencing-based quantification of Tuft cell chemosensory niches (16.2-fold stem cell enrichment). In a 1.6-million-bin human colorectal cancer cohort, FlashDeconv identifies neutrophil inflammatory microdomains co-localized with LAMP3^**+**^ mRegDC at the tumor–stroma interface, supported by independent single-cell spatial and cross-modality validation—mixed spatial niches that discrete-label summaries underrepresent but continuous proportions preserve.

## Introduction

Spatial transcriptomics (ST) technologies [1], such as Stereo-seq [2], Xenium [3], and Visium HD, are transforming our understanding of tissue architecture by mapping gene expression with increasingly fine spatial coordinates across large fields of view. As these technologies scale to atlas-level datasets comprising millions of spots or cells, the computational burden of analyzing such data has become a critical bottleneck. A fundamental task in ST analysis is cell type deconvolution, which infers the proportional composition of cell types within each spatial measurement unit by leveraging single-cell RNA sequencing (scRNA-seq) references.

Existing deconvolution methods—now numbering over seventy distinct approaches [4]—can be broadly distinguished by their modeling paradigms: generative probabilistic modeling versus constrained regression. Generative methods such as Cell2Location [5] and Stereoscope [6] achieve high accuracy by explicitly modeling count data distributions (e.g., Negative Binomial) via variational inference or maximum a posteriori (MAP) estimation, providing rigorous uncertainty quantification that is particularly valuable for small-scale studies requiring careful statistical inference. However, these methods require extensive iterative training, with runtimes scaling from hours to days for million-scale datasets. In contrast, regression-based approaches formulate the problem as constrained minimization, offering faster inference. This category includes the baseline NNLS, weighted variants such as MuSiC [7] and SpatialDWLS [8], likelihood-based regression (RCTD [9]), NMF decomposition (SPOTlight [10]), and deep-learning alignment (Tangram [11]). Benchmarks indicate that many methods fail to outperform NNLS consistently; only Cell2Location, RCTD, and MuSiC exceed baseline performance across all evaluation metrics [12]. However, most regression approaches treat spatial spots as independent observations, ignoring local tissue continuity. Methods that do incorporate spatial structure, such as CARD [13] with its conditional autoregressive (CAR) prior, typically rely on dense covariance matrices (*O*(*N*^2^) memory), precluding analysis of emerging high-resolution platforms with millions of spots.

Here, we present FlashDeconv, a deconvolution framework that combines accuracy, spatial awareness, and linear scalability. FlashDeconv leverages Randomized Numerical Linear Algebra (RandNLA) to reduce the original transcriptome (*G* ≈ 20, 000 genes) to a compact sketch (*d* ≈ 512) through informative gene selection followed by structure-preserving randomized projection. Unlike standard random projections that risk losing signals from rare cell types, our approach utilizes leverage-score importance sampling to preserve biological heterogeneity. Leverage scores—a classical tool from regression diagnostics and randomized numerical linear algebra [14, 15]—have been applied in single-cell biology to subsample *cells* while preserving rare populations [16, 17], with independent benchmarking confirming their effectiveness over alternative sketching strategies for spatial transcriptomics data [18]. FlashDeconv applies them to the complementary problem of compressing the *gene* dimension: rather than selecting which cells to retain, we use reference-derived leverage scores to weight which selected gene-space directions to preserve during randomized projection. Here, “structure” refers to the geometric organization of cell types in gene expression space, which leverage scores quantify by measuring each gene’s contribution to distinguishing cell type signatures. Combined with a Log-CPM data representation and a sparse graph Laplacian regularizer that scales as *O*(*N*) rather than *O*(*N*^2^), FlashDeconv achieves competitive benchmark accuracy while reducing runtime by orders of magnitude, enabling atlas-scale analysis on standard commodity hardware.

A critical challenge in deconvolution is that traditional dimension reduction methods—such as Principal Component Analysis (PCA) or Highly Variable Genes (HVG) selection—rely on *variance* as a proxy for biological information [19, 20]. Intuitively, variance measures how “loud” a gene is across the dataset—a quantity naturally dominated by abundant cell types. In contrast, leverage scores measure *distinctiveness*: whether a gene defines a unique direction in the transcriptomic space, regardless of how many cells express it. This distinction is biologically consequential: rare populations—including cancer stem cells comprising <1% of tumors [21, 22] and vascular endothelial cells that orchestrate tissue-specific stem cell niches [23]—can carry disproportionate functional significance in tissue homeostasis and pathology, yet their markers are systematically underrepresented by variance-based feature selection [24, 25]. The need to decouple biological importance from population frequency has motivated alternative approaches, including Gini-index-based gene selection [26] and deviance-based feature ranking [27]. We validate this variance-leverage decoupling through systematic experiments (Section 2), demonstrating that leverage scores identify biologically meaningful markers independently of cell type abundance. Applied to atlas-scale Visium HD data, FlashDeconv further enables systematic multi-resolution analysis, characterizing a tissue-specific “resolution horizon” for spatial information loss and revealing cellular architecture difficult to resolve at conventional spatial scales.

Taken together, FlashDeconv makes atlas-scale, multi-resolution spatial deconvolution practical on commodity hardware, supporting both million-bin Visium HD mapping and data-driven characterization of tissue-specific resolution horizons. In the following Results, we first describe the FlashDeconv framework and leverage-weighted sketching, then benchmark accuracy and linear scalability, and finally demonstrate atlas-scale and multi-resolution applications on Visium HD data.

## Results

### The FlashDeconv framework

FlashDeconv formulates spatial deconvolution as a constrained optimization problem in a compressed feature space (Fig. 1). The framework consists of three key design choices:

**Fig. 1.**
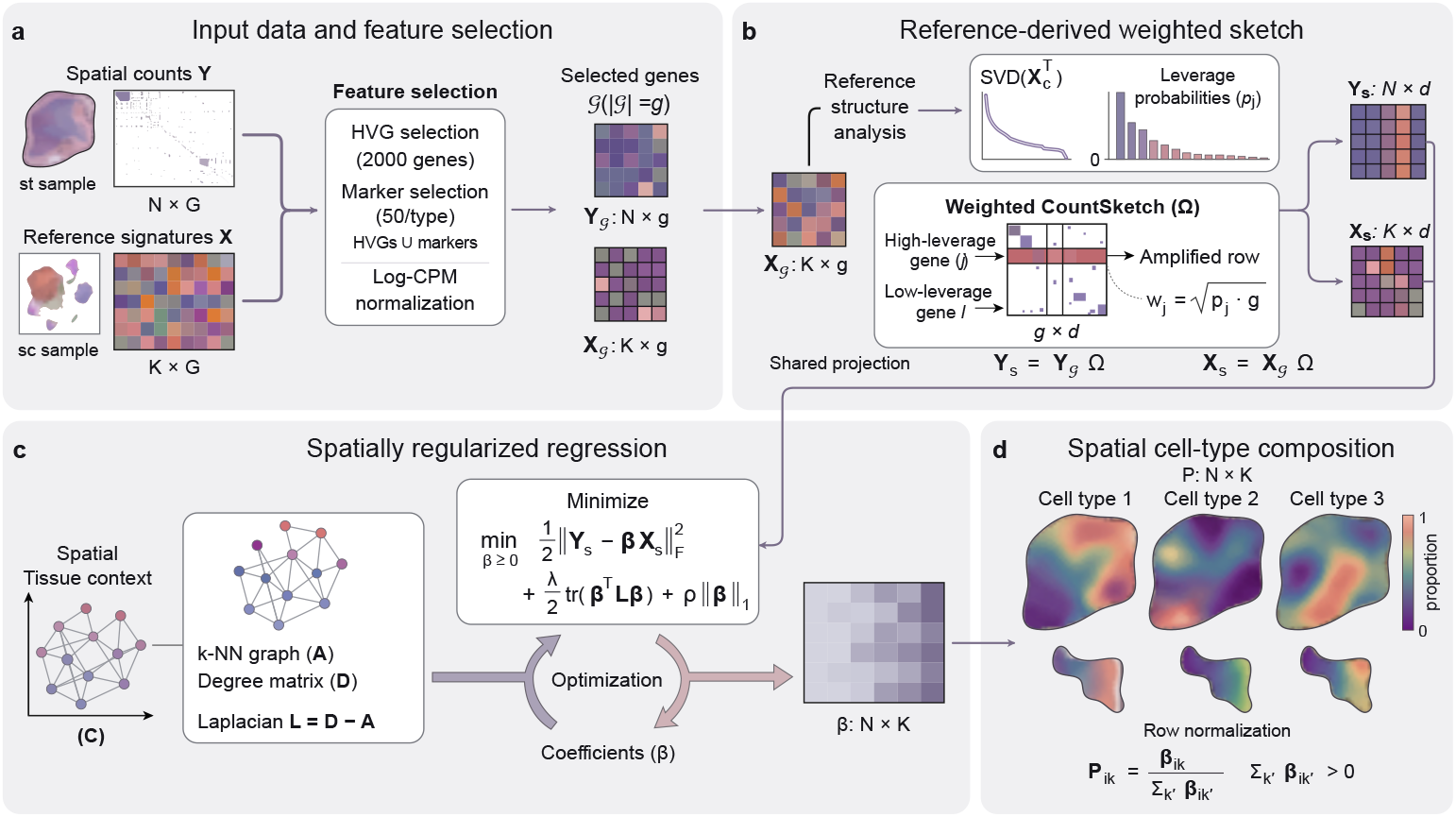
Overview of the FlashDeconv workflow. **(a)** Spatial counts *Y* ∈ ℝ^*N* ×*G*^ and reference signatures *X* ∈ ℝ^*K*×*G*^ provide the input. The selected gene set *G* is the union of spatial highly variable genes and reference cell-type markers, with *g* = |*G*|. Here, *Y*_*G*_ and *X*_*G*_ denote the selected matrices after library-size normalization and log transformation. **(b)** Reference-derived leverage scores determine gene-amplitude weights for a shared CountSketch projection Ω ∈ ℝ^*g*×*d*^. The reference SVD uses the transpose of the column-centered selected reference, denoted *X*_*c*_. Uniform random hashing and random signs are combined with leverage-based amplitude scaling, followed by column normalization. The same projection yields *Y*_*s*_ = *Y*_*G*_ Ω and *X*_*s*_ = *X*_*G*_ Ω, which feed the regression in c. The illustrated amplification is the weighting step before column normalization. **(c)** Spatial coordinates *C* define a *k*-nearest-neighbor graph with adjacency matrix *A*, degree matrix *D*, and Laplacian *L* = *D* − *A*. Nonnegative regression in the sketch space estimates coefficients *β* ∈ ℝ^*N* ×*K*^ with spatial and sparsity penalties, using block coordinate descent. **(d)** Row normalization converts the coefficients to cell-type proportions *P*_*ik*_ = *β*_*ik*_*/* ∑_*k*′_ *β*_*ik*′_ for positive row sums. The implementation assigns uniform proportions to all-zero coefficient rows. Proportions are reported at the observed spatial locations; the schematic tissue maps illustrate their spatial organization. Matrix entries and tissue maps are illustrative, rather than experimental results. The shared color bar in d denotes proportion, whereas the coefficient matrix in c uses a separate schematic sequential palette.

First, to address the extreme sparsity and mean-variance dependency of ST data, we employ a Log-CPM (Counts Per Million) transformation. While Pearson residuals are statistically principled for negative binomial count data [28], Log-CPM offers specific advantages for *L*_2_-based sketching: its bounded norm prevents high-expression genes from dominating the sketch space, while its logarithmic compression stabilizes the extreme dynamic range of spatial count data. This represents an engineering trade-off—sacrificing some statistical optimality for compatibility with randomized compression (Supplementary Note 2).

Second, we tackle the computational redundancy of gene expression via structure-preserving randomized sketching. Instead of solving the regression on the full transcriptome, we first restrict the problem to an informative gene set *G* and then project the selected gene space (*g* = |*G*|) into a lower-dimensional subspace using a sparse CountSketch matrix Ω. Unlike PCA, which maximizes explained variance and may obscure cell types that contribute little to global variance, our sketching matrix satisfies the Johnson-Lindenstrauss property [29], guaranteeing that Euclidean distances between cell type signatures are maintained in the compressed space with high probability. Crucially, the projection is weighted by statistical leverage scores [30, 31] derived from the single-cell reference. This ensures that marker genes defining transcriptomically distinct cell types—which often have high leverage despite low variance—are preserved with high probability, avoiding the signal loss that can occur with uniform subsampling or variance-based feature selection. For example, in liver scRNA-seq data, Central Vein Endothelial cells constitute only 2% of cells, yet their marker gene *Rspo3* achieves the highest leverage score (0.0105) among all genes due to its unique expression pattern that sharply distinguishes this population from all others. In contrast, Hepatocyte markers like *Egfr* exhibit low leverage scores (0.0007) despite high specificity, because Hepatocytes share metabolic gene programs with Cholangiocytes, reducing their geometric distinctiveness in the reference space. This illustrates that leverage captures *functional distinctiveness* rather than abundance: Hepatocytes are abundant but transcriptionally overlap with other lineages, while Central Vein Endothelial cells are rare but transcriptomically unique. Standard variance-based methods (PCA, HVG) show no statistical difference between functionally distinct and overlapping markers (MannWhitney U test, *p* = 0.97), whereas leverage-score ranking successfully identifies the structurally informative genes (Mann-Whitney U test, *p* = 0.0062). We validate this principle systematically in Section 2. Leverage-based weighting also addresses a second, algorithmic challenge: during the hashing step of CountSketch, high-abundance genes can dominate compressed dimensions and overwhelm rare cell markers assigned to the same hash bucket. By scaling selected genes in proportion to their leverage scores before compression (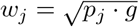, where *p*_*j*_ is the normalized leverage probability for selected gene *j*), FlashDeconv ensures that rare cell markers receive preferential amplification relative to their baseline magnitude, maintaining adequate signal strength despite random hash collisions [32] (see Methods for formal derivation and empirical validation across six tissues).

Third, we incorporate spatial information using a graph Laplacian regularizer. Unlike CARD, which constructs and operates on a dense *N* × *N* spatial kernel matrix with *O*(*N*^2^) memory and *O*(*N*^2^ · *K*) per-iteration complexity (where *K* is the number of cell types), our approach imposes spatial smoothness via a sparse penalty term Tr(*β*^*T*^ *Lβ*)—a formulation originally developed for structured genomic data [33] and shared with recent spatial-aware methods such as SDePER [34] and FAST [35]—where *L* is the graph Laplacian of the spatial neighbor network. This regularization induces local smoothness that is mathematically equivalent to a Gaussian Markov Random Field (GMRF) [36] but requires only sparse matrix operations. Because each spot connects to only *k* neighbors (default *k* = 6), the spatial term scales as *O*(*N* · *k*) rather than *O*(*N*^2^), yielding linear time complexity. This formulation allows FlashDeconv to model spatial autocorrelation across millions of spots using a fast Block Coordinate Descent (BCD) algorithm, a scale where dense matrix inversions are computationally prohibitive. Biologically, this regularization encodes the intuition that tissue composition varies smoothly: for a spot with sparse sequencing depth, the algorithm borrows strength from its spatial neighbors, allowing coherent tissue structures to emerge from noisy measurements.

### Leverage decouples biological identity from population abundance

A fundamental premise of FlashDeconv is that statistical variance conflates biological signal with population frequency, systematically disadvantaging rare cell types. To systematically test this claim, we designed a series of experiments that trace the evidence from mathematical principle through molecular function to spatial phenotype.

#### Abundance invariance test

We performed an *in silico* stress test using the mouse brain scRNA-seq reference (40,532 cells, 59 cell types, 31,053 genes). Starting from the native cell type distribution, we artificially downsampled the dominant oligodendrocyte population from 26.7% to 0.4%—a 67-fold reduction—while keeping all other populations constant. At each downsampling level, we identified the top 20 marker genes (by expression level in oligodendrocytes), computed both variance-based (normalized dispersion [37]) and leverage-based rankings across all genes, and tracked how the average marker rank changed (Fig. 2a). The results reveal a striking divergence: variance-based ranking degraded linearly with abundance, with marker rank dropping from 115 to 240 as the population shrank—a two-fold deterioration. In contrast, leverage-score ranking remained stable at rank ~150 throughout the entire range, demonstrating that leverage successfully decouples a gene’s discriminative power from its parent cell type’s numerical prevalence. Quantitatively, the stability can be attributed to leverage’s reduced coupling to mean expression: while gene variance correlates almost perfectly with mean expression (Spearman *ρ* = 0.998), leverage scores show substantially greater spread around this relationship (*ρ* = 0.965), enabling identification of low-abundance but high-specificity markers (Supplementary Fig. S5).

**Fig. 2.**
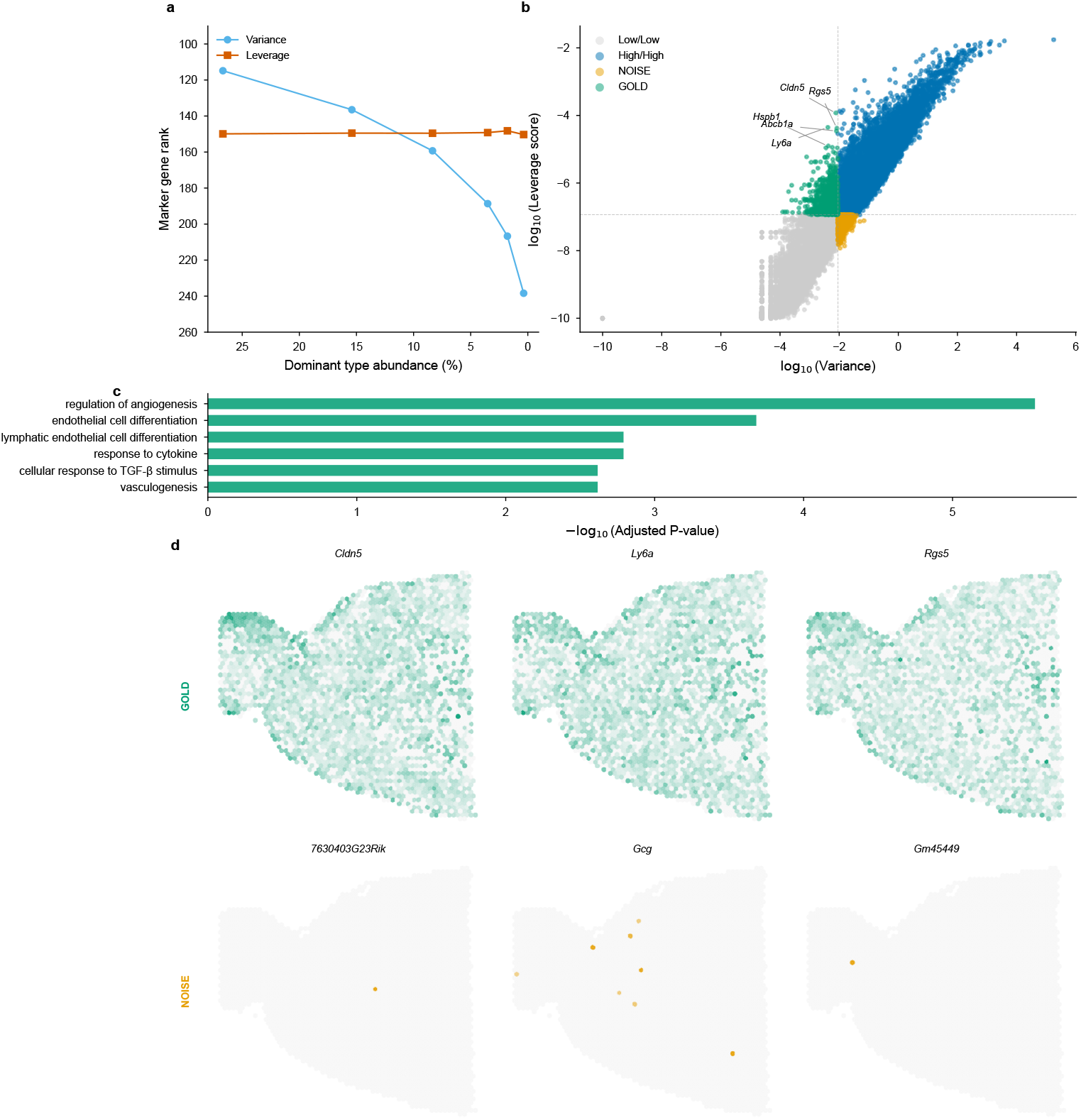
Leverage scores decouple biological identity from population abundance. **(a)** Abundance invariance test. Ranking stability of oligodendrocyte markers (top 20 genes by expression in oligodendrocytes) in mouse brain scRNA-seq (31,053 genes) as cell population is downsampled from 26.7% to 0.4%. Variance-based ranking (blue) degrades from rank 115 to 240—a two-fold deterioration. Leverage-score ranking (red) remains stable at rank ~150 regardless of population size, demonstrating true decoupling of biological identity from numerical prevalence. **(b)** The variance-leverage plane. Classification of 31,053 genes by variance (x-axis) and leverage score (y-axis). Structurally informative “GOLD” genes (green, low variance/high leverage) include vascular markers (*Cldn5, Rgs5, Ly6a*) that define rare anatomical structures; variance-dominated “NOISE” genes (orange, high variance/low leverage) contain 35% unannotated *Gm*-series transcripts versus 6% in GOLD, indicating that high variance alone does not ensure discriminative power. Cell type specificity analysis and spatial verification on Visium brain sections confirm that GOLD genes mark rarer cell types and reconstruct anatomical structure (Supplementary Fig. S31).

#### Gene quadrant analysis reveals systematic bias

To understand the broader implications of this decoupling, we mapped all 31,053 genes into a variance-leverage coordinate system (Fig. 2b). Four distinct quadrants emerged, defined by median log-transformed values to ensure equal partitioning without arbitrary threshold selection. The “GOLD” quadrant (low variance, high leverage) contains genes that standard HVG selection would discard but that carry high discriminative power—including classic vascular markers such as *Cldn5* (claudin-5, tight junction protein), *Rgs5* (pericyte marker), *Ly6a* (stem cell antigen), *Abcb1a* (blood-brain barrier transporter), and *Hspb1* (heat shock protein). These genes define rare but anatomically critical cell populations: brain endothelial cells and pericytes constitute <3% of brain tissue yet form the blood-brain barrier essential for neural function. Conversely, the “NOISE” quadrant (high variance, low leverage) contains genes with high variance but low cell-type specificity: 35% are unannotated *Gm*-series transcripts (predicted genes with unknown function) compared to only 6% in the GOLD set— a 6-fold enrichment indicating that variance-based selection systematically prioritizes genes lacking discriminative power for cell type identification.

#### Genome-wide specificity analysis

To verify that the GOLD/NOISE distinction reflects intrinsic biological structure rather than post-hoc cherry-picking, we performed a genome-wide cell type specificity analysis across all 59 cell types in the reference (Supplementary Fig. S31a). For every gene in the genome, we identified its primary cell type target based on expression specificity, without any manual pre-selection. Strikingly, GOLD genes preferentially mark significantly rarer cell populations (median target abundance 0.27%) compared to NOISE genes (median 0.51%; Mann-Whitney *U* test *p* = 3.25 × 10^−25^). The top cell type targeted by the GOLD gene set was Endothelial cells (0.05% abundance, 193 genes)—confirming that the variance-leverage plane objectively separates markers of rare anatomical structures from genes lacking cell-type discriminative power, without requiring prior knowledge of cell type identity. Extended analyses including abundance distributions and NOISE gene targets are provided in Supplementary Fig. S11.

#### Functional enrichment provides molecular validation

To confirm that GOLD genes share coherent biological functions, we performed Gene Ontology enrichment analysis using Enrichr [38, 39] with Benjamini-Hochberg FDR correction on both gene sets (Supplementary Fig. S13c). GOLD genes showed highly significant enrichment for vascular biology: *regulation of angiogenesis* (FDR-adjusted *p* = 2.8×10^−6^), *endothelial cell differentiation* (FDR-adjusted *p* = 2.1 × 10^−4^), *vasculogenesis* (FDR-adjusted *p* = 2.4 × 10^−3^), and *blood vessel morphogenesis*—consistent with the Endothelial cell enrichment identified in the specificity analysis. In stark contrast, NOISE genes yielded *zero* significant GO terms at FDR-adjusted *p* < 0.05, consistent with their enrichment for unannotated transcripts and confirming that high variance without high leverage indicates expression variability without coordinated cell-type specificity.

#### Spatial visualization confirms anatomical relevance

The ultimate test of a gene’s biological importance is whether it corresponds to recognizable tissue structure. We visualized count-matched GOLD and NOISE gene pairs on mouse brain Visium sections (Supplementary Fig. S31b), selecting pairs with comparable total UMI counts to control for expression-level confounding. GOLD genes (*Cldn5, Esam, Hspb1*) reconstructed clear vascular anatomical patterns—bright spots tracing blood vessel distributions consistent with known brain vasculature—while NOISE genes with similar expression levels (*Slc12a6, Tada2b, Pde12*) showed diffuse, spatially unstructured distributions. To quantify this across the full gene sets, we computed a spatial structure score measuring the ratio of global to local variance (higher values indicate organized spatial patterns). GOLD genes achieved a structure score of 1.33 ± 0.23, significantly exceeding NOISE genes at 0.87 ± 0.54 (Mann-Whitney *p* = 5.6 × 10^−5^). This spatial validation closes the evidence loop: genes identified as important by leverage scores correspond to real anatomical structures, while genes prioritized by variance alone do not.

#### Downstream impact on deconvolution accuracy

Finally, we tested whether these theoretical insights translate to practical performance gains through systematic ablation experiments (Supplementary Fig. S7). We compared four dimensionality reduction approaches within the FlashDeconv framework—Leverage-Score Sketching (LSS), Uniform CountSketch, PCA, and HVG selection—using identical downstream optimization across 54 Silver Standard datasets (6 tissues × 9 abundance scenarios, excluding 2 replicate conditions). LSS achieved rare cell type detection of *r* = 0.35, compared to Uniform (*r* = 0.14, +147%), PCA (*r* = 0.16, +124%), and HVG (*r* = 0.12, +197%). A stress test revealed the critical distinction: when cell type abundance was artificially reduced to 0.17%, LSS maintained detection (*r* = 0.62) while PCA (*r* = 0.02) and Uniform (*r* = 0.08) showed near-zero correlation. At 0.03% abundance, LSS remained positive (*r* = 0.33) while alternatives showed negative correlation, consistent with signal loss under extreme rarity. These results demonstrate that leverage-based sketching preferentially preserves rare cell type signals across diverse tissue contexts. Importantly, this improvement is not at the expense of common cell types in this benchmark: per-cell-type analysis across all 76 cell types shows that LSS improves Pearson correlation for every abundance category—rare (+0.198), moderate (+0.112), and abundant (+0.059) relative to uniform sketching (Supplementary Fig. S7).

#### Cross-tissue generalization

To confirm that the GOLD/NOISE framework is not specific to brain tissue, we repeated the quadrant analysis on mouse kidney scRNA-seq data from the Spotless benchmark (7,501 cells, 16 cell types). Podocytes—glomerular epithelial cells critical for kidney filtration—constitute only 0.17% of the reference population, representing an extreme test case for rare cell detection. All five canonical podocyte markers (*Nphs1, Nphs2, Podxl, Synpo, Wt1*) appeared in the GOLD quadrant, while markers of abundant tubular cell types (60% of cells) fell in high-variance regions (Supplementary Fig. S13). This cross-tissue consistency confirms that the variance-leverage separation reflects a general property of gene expression structure rather than a tissue-specific artifact.

Together, these experiments establish that leverage scores provide a principled, abundance-independent measure of biological informativeness. Critically, the ablation comparison isolates the effect of leverage weighting during compression: leverage-weighted and uniform sketching operate on identical gene sets (HVG ∪ markers), yet leverage weighting improves rare cell detection by 147%—demonstrating that the benefit arises from preserving marker signals during the *G*-to-512 projection, not from gene selection per se (see Methods for two-stage design rationale). By incorporating leverage-weighted importance sampling into the sketching process, FlashDeconv corrects a systematic bias inherent to variance-based feature selection, enabling accurate detection of rare but biologically critical cell populations.

### FlashDeconv achieves competitive accuracy on synthetic benchmarks

We systematically evaluated FlashDeconv using the *Spotless* benchmark suite [12], a reproducible pipeline for benchmarking cell type deconvolution methods. Spotless provides two types of ground-truth data: (1) Silver Standards, synthetic “pseudospots” generated by computationally mixing single-cell transcriptomes with known cell type proportions, and (2) Gold Standards, real spatial transcriptomics data (seq-FISH+, STARMap) where ground-truth composition is derived from co-registered single-molecule FISH imaging.

We benchmarked FlashDeconv on all 54 Silver Standard datasets from Spotless (6 tissues × 9 abundance patterns). Across these datasets representing various tissue types and abundance scenarios (e.g., dominant cell types, rare cell types, uniform distributions), FlashDeconv achieved a mean Pearson correlation of 0.944, mean RMSE of 0.053, and mean JSD of 0.047—ranking #1 among 13 methods on all three aggregate metrics (Fig. 3a,b; Supplementary Table S2).

**Fig. 3.**
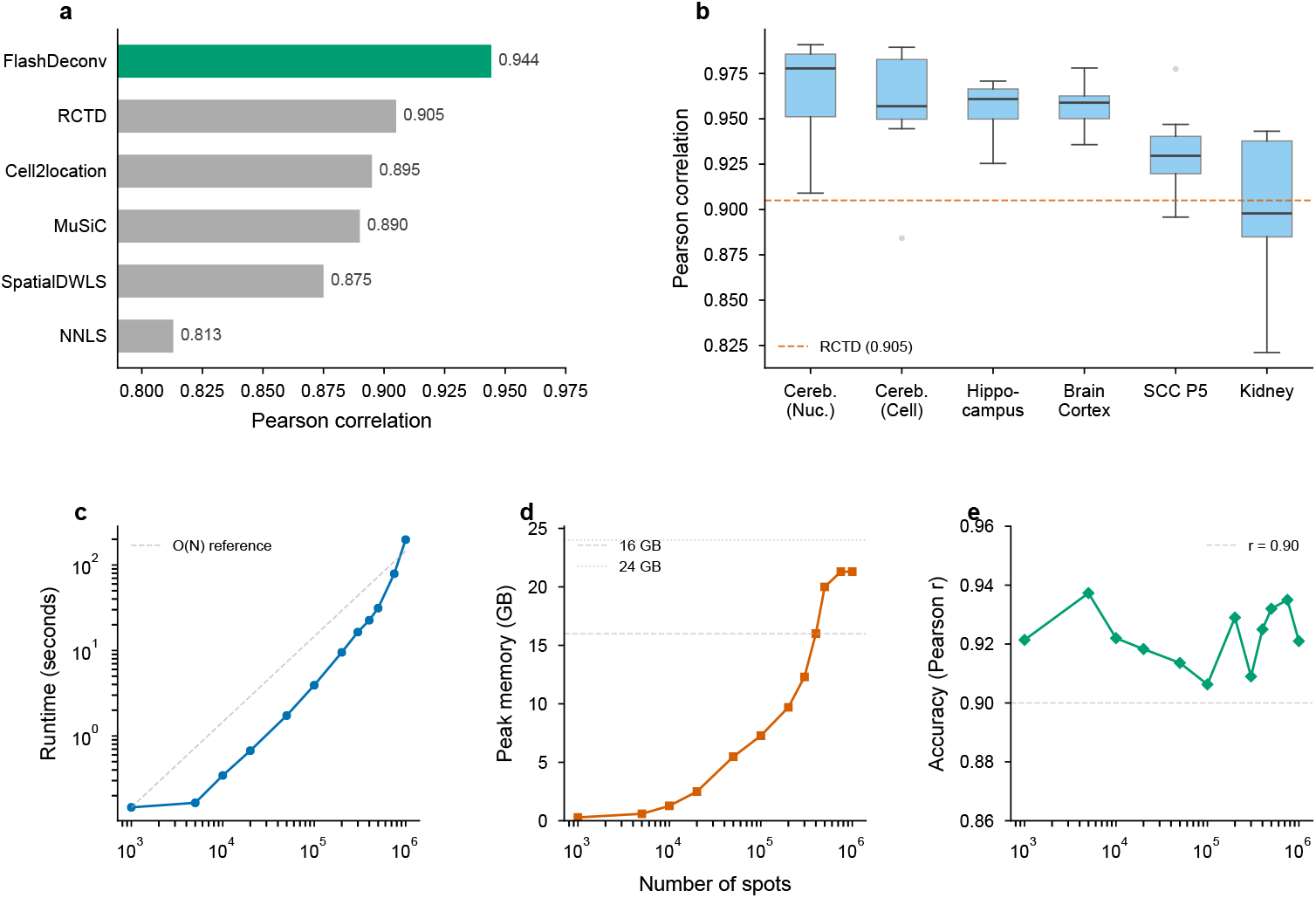
FlashDeconv achieves competitive accuracy with linear scalability. **(a)** Comparison with published methods on 54 Silver Standard datasets. FlashDeconv achieves mean Pearson = 0.944 and JSD = 0.047, compared to RCTD (Pearson = 0.934, JSD = 0.050) and Cell2Location (0.918, 0.061). Synthetic performance does not always translate to real-data ranking (see Section 2). **(b)** Performance stratified by tissue type. FlashDeconv maintains high accuracy across brain tissues (Pearson *>* 0.95 for cortex, cerebellum, hippocampus), with lower performance on kidney (Pearson = 0.90) due to highly correlated cell type signatures. **(c)** Runtime comparison with RCTD and Cell2Location (Spotless benchmark data at 20,000 genes). FlashDeconv (8 CPU cores, Numba JIT) processes 100K spots in under 4 seconds and 1M spots in approximately 3 minutes, approaching the *O*(*N*) reference line. Published RCTD and Cell2Location runtime data extend to 10K spots; extrapolation from those reported operating points makes larger-scale analyses impractical under the benchmark settings. Peak memory usage and accuracy preservation across scales are shown in Supplementary Fig. S32.

Notably, FlashDeconv exceeded the baseline NNLS method (Pearson = 0.850, JSD = 0.118) and matched or exceeded the performance of computationally intensive methods such as RCTD (Pearson = 0.934, JSD = 0.050) and Cell2Location (Pearson = 0.918, JSD = 0.061). FlashDeconv achieved mean AUPR = 0.957 (*n* = 54 datasets) on rare cell type detection, comparable to RCTD (0.951) and Cell2Location (0.921), supporting the conclusion that leverage-score-based sketching preserves signals from low-abundance populations. Per-cell-type evaluation across 684 cell-type instances (76 types × 9 patterns × 6 tissues) further quantifies performance by abundance: rare types (<5%) achieve mean Pearson = 0.856, AUPRC = 0.871, and F1 = 0.614; moderate types (5–15%) achieve Pearson = 0.933, AUPRC = 0.961, F1 = 0.803; abundant types (>15%) achieve Pearson = 0.961, AUPRC = 0.995, F1 = 0.904 (Supplementary Table S2; Supplementary Data). An ablation study further confirms that the graph Laplacian regularization provides a net benefit for rare cell type detection (AUPRC +0.008 for rare types across 76 cell types in 6 tissues), although with a mild precision–recall trade-off consistent with spatial smoothing (Supplementary Fig. S10).

To validate performance on platform effects, we utilized the “Gold Standard” seq-FISH+ and STARMap datasets. On the STARMap dataset (108 spots, 12 cell types), FlashDeconv achieved Pearson = 0.665, ranking 3rd out of 13 methods on Pearson (behind SpatialDWLS at 0.712 and RCTD at 0.679), while achieving the lowest JSD (= 0.252) and highest AUPR (= 0.661) among all methods. On the seqFISH+ olfactory bulb dataset (7 FOVs), FlashDeconv achieved mean Pearson = 0.608, ranking 7th—consistent with reduced benefit of sketching and spatial regularization in microscale regimes (<10 spots per FOV). On the seqFISH+ cortex dataset, all methods achieve low accuracy (best method Pearson < 0.30), reflecting the extreme sparsity of these micro-scale measurements (Supplementary Fig. S8; Supplementary Table S2).

### Real-world validation on Visium case studies

To assess performance on real Visium data with biologically grounded ground truth, we evaluated FlashDeconv on two case studies from the Spotless benchmark: mouse liver tissue sections and a melanoma tumor dataset. These case studies provide direct comparison with 12 competing methods on actual spatial transcriptomics data rather than simulations.

The liver dataset consists of four Visium slides (5,762 total spots) with zonation annotations and matched snRNA-seq reference (133,779 cells, 9 cell types). Following the Spotless protocol, we evaluated two metrics: Jensen-Shannon Divergence (JSD) against snRNA-seq-derived proportions, and Area Under the Precision-Recall curve (AUPR) for detecting portal and central vein endothelial cells based on spatial zonation patterns (per-spot Pearson and RMSE are not computable for this benchmark because ground truth consists of tissue-level proportions; see Supplementary Note). FlashDeconv achieved a JSD of 0.0561, ranking 3rd out of 13 methods, behind only RCTD (0.0334) and Cell2Location (0.0352), and halving the NNLS baseline (0.1056). On the AUPR metric, FlashDeconv achieved 0.66, ranking 7th. Per-cell-type decomposition reveals that this JSD–AUPR discrepancy originates from a specific identifiability constraint: the liver reference contains three endothelial populations—portal vein EC, central vein EC, and LSECs—whose signatures are nearly collinear (pairwise cosine similarity >0.975). Portal vein EC tissue-level proportion is estimated almost exactly (2.0% predicted vs. 2.0% ground truth, contributing <0.01% of total JSD), but the collinear signatures prevent reliable spot-level attribution, yielding AUPR = 0.412. Central vein EC exhibits the complementary failure: its signal is absorbed by LSECs (0.1% predicted vs. 2.0% ground truth), but the residual signal is well-localized to the central zone (AUPR = 0.909). This asymmetry—accurate tissue-level composition but imprecise spot-level localization for collinear subtypes—is a fundamental property of linear deconvolution when cos *θ* > 0.97, not specific to FlashDeconv (Supplementary Note).

Beyond accuracy metrics, we evaluated FlashDeconv’s robustness to reference protocol variations using the Spotless liver reference sensitivity test. This test measures prediction consistency when using three different reference protocols: exVivo scRNA-seq, inVivo scRNA-seq, and snRNA-seq. FlashDeconv achieved a stability JSD of 0.0138, ranking 1st out of 13 methods—25% better than RCTD (0.0185), the second-best performer. This robustness indicates that FlashDeconv’s predictions are less sensitive to the specific reference protocol choice, a practically valuable property given the heterogeneity of available scRNA-seq references.

The melanoma dataset includes three Visium slides (7,557 spots) with Molecular Cartography-derived ground truth proportions for 7 cell types. Only JSD is evaluable for this benchmark, as the ground truth provides tissue-level proportions without spatial zonation patterns required for AUPR (Supplementary Note). We evaluated FlashDeconv using the default sketch dimension (*d* = 512) and the full reference (15 cell types), identifying a fundamental trade-off between distribution-based similarity and absolute abundance estimation. When optimized for JSD—which penalizes small relative errors in rare cell types—FlashDeconv using Pearson residual preprocessing achieved a JSD of 0.015, ranking 3rd among 13 methods, behind only Cell2Location and SPOTlight. Under this configuration, FlashDeconv accurately captured rare T-cell populations (3.3% predicted vs. 0.3% with log-CPM; ground truth 4.7%) but underestimated the dominant melanocytic cells (75.5% vs. 84.8% ground truth). Alternatively, using log-CPM preprocessing prioritized the dominant signal, yielding a more accurate melanocytic proportion (81.2%) but a higher JSD (0.033, rank 7) due to increased sparsity in rare cell estimates. This sensitivity analysis demonstrates that FlashDeconv’s flexible preprocessing allows users to prioritize either broad-spectrum distribution fidelity or dominant-cell accuracy, with both modes maintaining robustness against the extreme collinearity of malignant cell states (*κ* = 63.4; Supplementary Note 2).

These results demonstrate that FlashDeconv achieves competitive accuracy on real spatial data, with performance varying across tissue types. The liver dataset illustrates a characteristic pattern: most cell type signatures are well-separated, yielding strong overall composition accuracy (JSD rank 3), but the endothelial triplet’s near-degeneracy (cos *θ* > 0.975) limits spot-level localization for those specific subtypes (AUPR rank 7). Notably, rare cell type markers in liver (Cholangiocytes 1.16%, Mesothelial cells 0.53%, Portal/Central Vein Endothelial cells ~2%) exhibit significantly higher leverage scores than markers of abundant types (*p* = 0.0062), a separation absent in variance-based ranking (*p* = 0.97; Fig. 2)—extending the variance-leverage decoupling established on brain tissue (Section 2) to a second organ. On the melanoma dataset, where malignant cell states exhibit extreme transcriptomic similarity (*r* > 0.98 for all 42 pairwise correlations; Supplementary Fig. S4), FlashDeconv with Pearson preprocessing achieves accuracy comparable to top probabilistic methods (rank 3). This demonstrates that appropriate preprocessing can partially overcome the collinearity challenge inherent in linear deconvolution, though at the cost of reduced accuracy for dominant populations—a trade-off that users can navigate based on their analytical priorities (Supplementary Note 2). This variability is consistent with the Spotless study’s conclusion that no single method excels universally across all tissue contexts [12]. Complete aggregate metrics (Pearson, RMSE, JSD, AUPR) for all 13 methods across all benchmark types are provided in Supplementary Table S2, with per-cell-type breakdowns (AUPRC, precision, recall, F1) in Supplementary Data.

### Linear scalability enables atlas-scale deconvolution

The defining advantage of FlashDeconv is its scalability. We benchmarked runtime and memory usage across datasets ranging from 10^3^ to 10^6^ spots (Fig. 3c–e). For a dataset with 100,000 spots, FlashDeconv completed the analysis in under 4 seconds. In contrast, deep learning methods face severe computational constraints at much smaller scales: benchmark studies report that Stereoscope requires over 8 hours on datasets with only 10,000 spots [12], and Cell2Location exhibits runtimes nearly 100-fold longer than regression-based alternatives [40]. Even fast regression-based methods face scalability constraints: per RCTD documentation, 11,000 spots require approximately 20 minutes on 4 cores; extrapolating linearly, 1 million spots would require over 30 hours—compared to FlashDeconv’s 3 minutes for the same scale.

Crucially, FlashDeconv exhibits *O*(*N*) linear scaling for both time and memory. On commodity hardware (32GB unified memory, no GPU), we successfully deconvolved a simulated 1-million-spot dataset in approximately 3 minutes. In contrast, probabilistic methods encounter severe scalability barriers at much smaller scales: CARD developers report that Cell2Location could not be applied to Slide-seqV2 data (~20,000 spots) due to computational burden [13], and benchmark studies observe memory errors when processing datasets of similar size on standard GPU hardware [41]. This efficiency enables interactive, iterative analysis of atlas-scale data without specialized hardware. Beyond raw throughput, linear complexity also opens systematic multi-resolution analysis—deconvolving the same tissue at multiple bin sizes to probe how spatial resolution affects biological inference—an investigation we pursue in Section 2 after first demonstrating practical utility on human cancer data.

This scalability advantage is rooted in two fundamental algorithmic design choices. First, FlashDeconv operates in a compressed sketch space rather than the full gene space. Standard regression-based methods scale with the number of genes *G*: NNLS’s active-set algorithm requires *O*(*N*· *G* · *K*^2^) due to iterative QR decomposition over up to *K* variables, and RCTD’s iteratively reweighted least squares involves Hessian computation over all cell type pairs, also yielding *O*(*N* · *G* · *K*^2^). FlashDeconv first restricts the problem to an informative gene set of size *g* and then solves the regression problem in a *d*-dimensional sketch space, reducing the data-fit precomputation to *O*(*N* · *d* · *K*) (Methods). With typical parameters *g* ≈ 2,500 and *d* = 512, the sketch provides a further reduction beyond feature selection, while the full pipeline reduces the original *G* ≈ 20,000-gene transcriptome to 512 sketch dimensions. We note that methods using HVG pre-selection (typically 2,000 genes) would see a smaller speedup factor from sketching alone. However, FlashDeconv’s scalability advantage stems primarily from two additional factors: (1) precomputation of 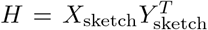 makes BCD iteration complexity *O*(*N* · *K*^2^) independent of gene dimension (Supplementary Note 1), and (2) sparse graph Laplacian regularization scales as *O*(*N* · *k*) rather than *O*(*N*^2^) for dense kernel methods. Furthermore, leverage-score weighting provides substantial accuracy advantages over HVG selection, particularly for rare cell types (Supplementary Fig. S7). These combined advantages explain the 1–2 orders-of-magnitude runtime reduction observed in our empirical benchmarks.

Second, FlashDeconv employs sparse graph-based spatial regularization rather than dense covariance modeling. Methods like CARD that model spatial dependencies through dense *N* × *N* kernel matrices incur *O*(*N*^2^) memory costs and *O*(*N*^2^ · *K*) per-iteration complexity for matrix operations, rendering them prohibitive for datasets with *N* > 10,000 spots. FlashDeconv instead constructs a sparse *k*-nearest-neighbor graph, where each spot connects to only *k* = 6 neighbors. The resulting graph Laplacian regularization scales as *O*(*N* · *k*) in both time and space, maintaining linear complexity even as *N* grows to millions. A detailed complexity comparison across methods is provided in Supplementary Table S1.

### Atlas-scale deconvolution of human colorectal cancer at 8 *µ*m resolution

To demonstrate FlashDeconv’s utility for atlas-scale human cancer analysis, we applied it to Visium HD data from a colorectal cancer (CRC) cohort [42] comprising three patients with 1,595,565 total bins at 8 *µ*m resolution and 38 cell types. FlashDeconv processed the entire cohort in 153 seconds at a throughput of approximately 10,400 bins per second, generating spatially resolved cell type proportion maps (Fig. 4a).

**Fig. 4.**
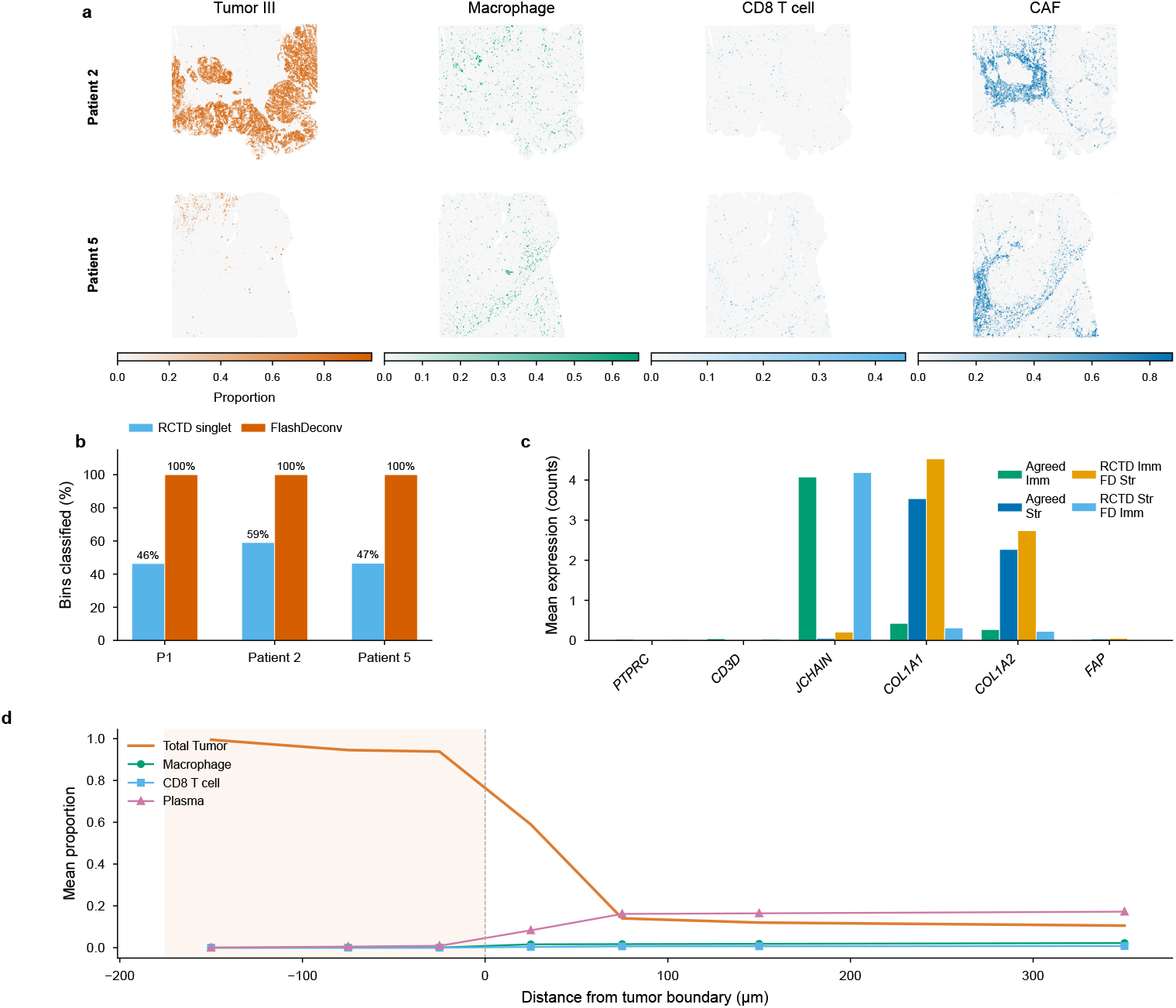
Atlas-scale deconvolution of human colorectal cancer at 8 *µ*m resolution. **(a)** Spatial distribution of four key cell types (Tumor III, Macrophage, CD8 T cell, CAF) in two representative CRC patients (P2, P5) at 8 *µ*m Visium HD resolution, showing heterogeneous tumor microenvironment architecture. **(b)** Zoom-in comparison at a 400 *µ*m tumor–immune interface (10x CRC Demo, 8 *µ*m). Each bin is colored by lineage proportion (Tumor, Immune, Stromal) using an additive Okabe–Ito palette. FlashDeconv (left) produces continuous proportions for every bin, revealing smooth transitions at cell-type boundaries. RCTD doublet mode (right) provides high-confidence discrete labels for 60% of bins, withholding classification for the remainder (light gray); FlashDeconv captures continuous mixing gradients across all bins. **(c)** Independent marker gene validation on 71,769 disputed bins where FlashDeconv and RCTD disagree on immune vs. stromal classification. Canonical lineage markers (PTPRC, CD3D, JCHAIN for immune; COL1A1, COL1A2, FAP for stromal) confirm FlashDeconv’s assignment in 19/22 gene-direction verdicts (86.4%), with unanimous agreement (11/11) when RCTD classifies as stromal but FlashDeconv assigns as immune. **(d)** Immune cell infiltration gradient as a function of signed distance from the tumor-stroma boundary, averaged across patients (shaded region = tumor interior). Total tumor proportion drops from ~94% at −25 *µ*m to ~14% at +75 *µ*m. Among immune populations, Plasma cells exhibit the steepest gradient (~16% in peri-tumoral stroma), followed by macrophages (~2%) and CD8 T cells (~0.8%). Cancer-associated fibroblasts define a distinct stromal layer peaking at ~14% near the boundary. Data: Visium HD CRC cohort from Oliveira et al. [42], 3 patients, 1,595,565 total bins at 8 *µ*m resolution, 38 cell types. Processing time: 153 seconds for the entire cohort (~10,400 bins/s).

A key distinction between continuous regression and discrete classification emerges at 8 *µ*m resolution. Oliveira et al. [42] ran RCTD in doublet mode—the recommended setting for high-resolution platforms—which restricts each bin to at most two cell types; although RCTD also provides an unrestricted full mode, simultaneously fitting all *K* = 38 proportions from the ~10–50 UMIs typical of 8 *µ*m bins is severely underdetermined without sparsity regularization, necessitating this restriction. RCTD provides confident classifications for 46–59% of bins (singlets) across the three patients, withholds 5–7% at its quality threshold, and assigns the remainder doublet labels. This conservative gating is a principled design choice: at 8 *µ*m resolution, individual bins are smaller than most cells, and mRNA diffusion causes nearly all bins to contain mixed signals from multiple cell types—a regime where discrete labeling is inherently approximate. A direct comparison at a tumor–immune interface illustrates the output-type difference: FlashDeconv produces continuous lineage-proportion gradients for every bin, while RCTD provides high-confidence discrete labels for 60% of bins, withholding classification for the remainder (Fig. 4b). FlashDeconv’s regression framework outputs continuous proportion estimates across all 38 cell types for every bin, achieving complete coverage of the tissue.

To independently validate accuracy on the 71,769 bins where FlashDeconv and RCTD disagree on immune versus stromal classification, we examined expression of canonical lineage markers (PTPRC, CD3D, JCHAIN, COL1A1, COL1A2, FAP) that serve as ground truth independent of either method’s output (Fig. 4c). FlashDeconv’s assignment was confirmed as correct in 19 of 22 gene-direction verdicts (86.4%). Notably, in the 11 cases where RCTD classified bins as stromal but FlashDeconv assigned them as immune, marker gene expression unanimously supported FlashDeconv—these bins expressed immune markers (PTPRC, CD3D, JCHAIN) at levels comparable to agreed-immune bins.

Analysis of immune cell infiltration as a function of signed distance from the tumor-stroma boundary revealed a sharp transition (Fig. 4d). Immune cell types were essentially absent inside the tumor and rose within the first 50 *µ*m of the stromal side. Plasma cells exhibited the steepest gradient, accumulating to ~16% in the peritumoral stroma, consistent with their known enrichment at the tumor-stroma interface in microsatellite-stable CRC [42]. Cancer-associated fibroblasts (CAFs) peaked at 50– 100 *µ*m outside the boundary (~14%), defining a fibroblast-rich peri-tumoral zone. CD8 T cells showed the weakest infiltration (~0.8% in distant stroma), consistent with the limited T-cell infiltration characteristic of microsatellite-stable colorectal tumors [42]. These continuous spatial gradients, detectable only at ≤8 *µ*m resolution, recapitulate the immune infiltration patterns described in the Immunoscore literature [43], demonstrating that FlashDeconv can resolve the spatial organization of immune populations at scales relevant to established tissue classification frameworks.

Beyond global infiltration patterns, FlashDeconv’s complete tissue coverage revealed discrete neutrophil inflammatory microdomains at the tumor–stroma interface in all three patients (Fig. 5a). These aggregates—bins with ≥10% Neutrophil proportion forming spatially coherent clusters—exhibited the strongest spatial self-enrichment of any cell type in the 38-type dataset (16–56×). Analysis of the surrounding microenvironment uncovered two distinct spatial contexts (Fig. 5b): stromal-resident aggregates were surrounded by a vascularized innate immune niche enriched for macrophages, mRegDC (mature regulatory dendritic cells that limit anti-tumor immunity through a conserved immunoregulatory program [44]), and mast cells, while tumor-proximal aggregates showed broad immune depletion—a dichotomy consistent across all three patients. Six neutrophil-specific marker genes independently confirmed the identity of these microdomains (11–63-fold enrichment; Fig. 5c), with the signal amplifying rather than diminishing when restricting to high-UMI bins (≥200), ruling out low-quality deconvolution artifacts (Fig. 5d). The niche structure was most pronounced at 8 *µ*m resolution, decaying at coarser scales (Fig. 5e; Supplementary Note 9). These microdomains are largely missed by discrete-label summaries: of the 16,827 hotspot bins, RCTD’s doublet mode classified 2.3% as Neutrophil singlets and withheld classification for 61%, reflecting the inherent difficulty of assigning discrete labels to bins where neutrophils are one component among continuously mixed cell types.

**Fig. 5.**
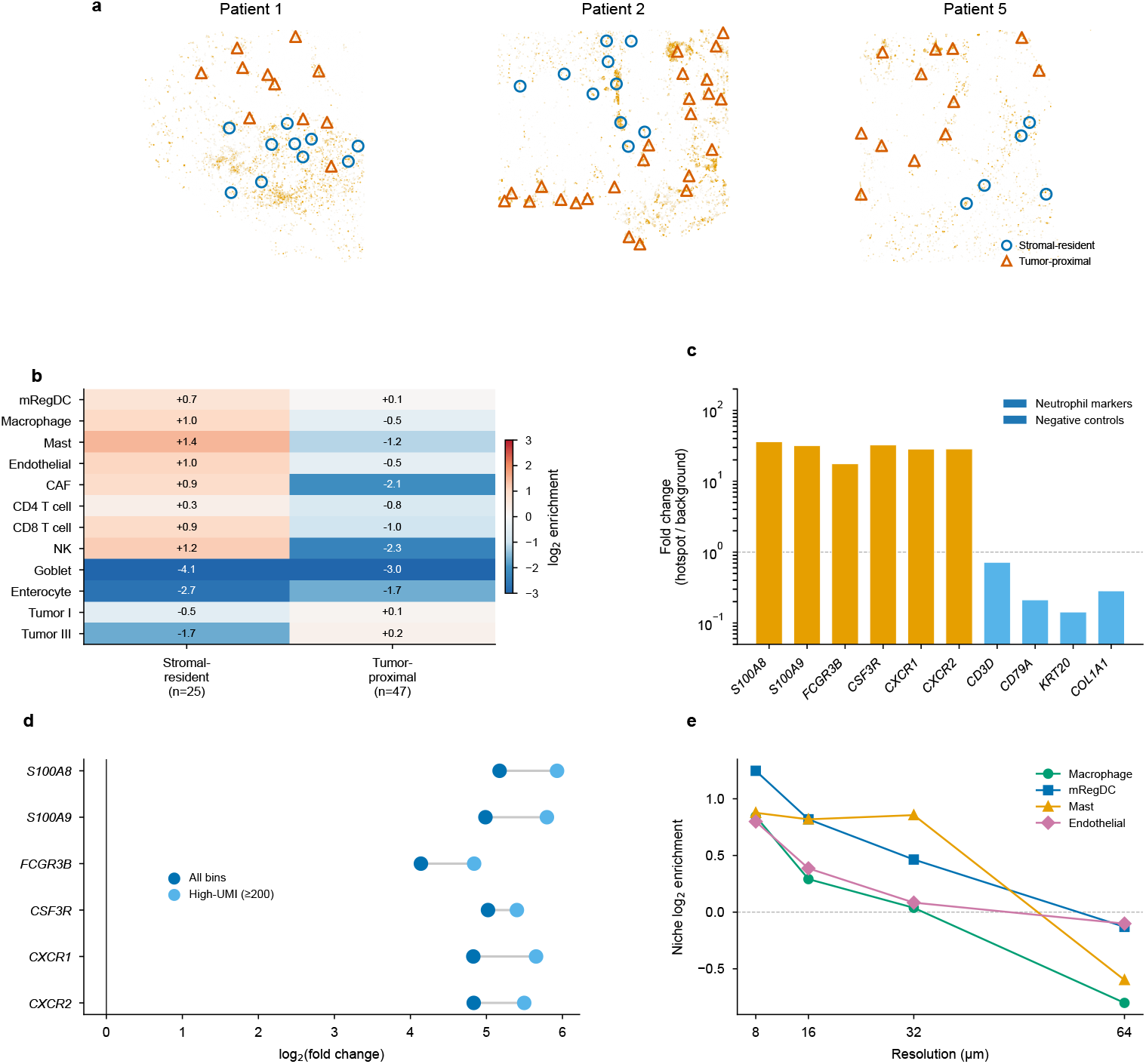
Neutrophil inflammatory microdomains at the CRC tumor–stroma interface reveal two distinct spatial contexts. **(a)** Spatial distribution of neutrophil hotspot bins (≥10% proportion) across three patients. Aggregate centroids classified as stromal-resident (blue circles) or tumor-proximal (orange triangles) based on niche tumor content. **(b)** Niche composition of the two aggregate types (median log_2_ enrichment pooled across patients). Stromal-resident aggregates are surrounded by a vascularized innate immune niche (Macrophage, mRegDC, Mast, Endothelial enriched), while tumor-proximal aggregates show broad immune depletion. **(c)** Marker gene validation. Neutrophil-specific genes show 11–63-fold enrichment in hotspot versus background bins; negative control genes show no enrichment or depletion. Y-axis: log-scale fold change. **(d)** UMI-stratified robustness. Log_2_ fold change of neutrophil marker genes in all hotspot bins versus high-UMI (≥200) bins, averaged across patients. The enrichment signal amplifies in high-UMI bins, ruling out low-quality deconvolution artifact. **(e)** Multi-resolution niche stability. Enrichment of key co-localized cell types decays from 8 to 64 *µ*m, demonstrating that the microenvironment structure requires high spatial resolution (≤8 *µ*m) to resolve. Data: Visium HD CRC cohort [42], 1,595,565 total bins, 38 cell types.

To orthogonally validate these deconvolution estimates, we leveraged Xenium in situ sequencing data from an adjacent serial section of the same tissue (Patient 1 [42]), which provides single-cell-resolution ground truth independent of any deconvolution model. We annotated 289,352 Xenium cells by correlation with the scRNA-seq reference, then computationally aggregated them into spatial bins of 8–128 *µ*m to create pseudo-bulk mixtures with known composition. FlashDeconv recovered lineage-level proportions with Pearson *r* > 0.88 across all resolutions, even at 8 *µ*m where bins contained a median of one cell (Supplementary Note 7, Supplementary Fig. S23). A systematic cross-method comparison at finer resolution (4–32 *µ*m pseudo-Visium HD bins, Xenium panel as self-reference) confirmed that FlashDeconv achieved the highest accuracy among four methods at all resolutions, with the advantage largest at the finest scales (4 *µ*m AUPRC: 0.87 vs. 0.62 for NNLS and 0.60 for marker scoring; Supplementary Table S3). Comparison of global cell type proportions across platforms revealed that at the coarser lineage level (6 categories), both FlashDeconv and RCTD agreed well with the Xenium reference (*r* = 0.96 and *r* = 0.94, respectively), but at the individual cell-type level (38 types), FlashDeconv maintained strong correspondence (*r* = 0.78) while RCTD singlet-derived proportions showed no systematic agreement (*r* = −0.02)—consistent with the information loss inherent in collapsing continuous mixtures to discrete labels.

FlashDeconv also provides per-spot uncertainty estimates at negligible additional cost. Because the BCD solver’s per-coordinate update denominator equals the diagonal of the Hessian, a Laplace approximation yields closed-form variance estimates for each cell type proportion in each bin. On the Xenium CRC data at 8 *µ*m (241,351 bins, 38 cell types), 95% analytical confidence intervals achieved 97.7% empirical coverage against ground truth, with CI width monotonically predicting estimation error (Supplementary Note, Supplementary Fig. S35). The analytical uncertainty computation added 1.1 seconds to the 7.6-second deconvolution—less than 15% overhead for 241K bins.

FlashDeconv’s accuracy extends beyond CRC: application to human ovarian cancer recovers the cell type composition differences between response groups reported by Denisenko et al. [45] (Supplementary Fig. S22).

### Scale-space analysis of Visium HD reveals resolution-dependent information loss

While the preceding CRC analysis demonstrates FlashDeconv’s practical utility for atlas-scale tumor microenvironment mapping at 8 *µ*m resolution, the Visium HD platform raises a fundamental question beyond any single biological application: at what resolution does cellular information begin to collapse? We leveraged FlashDeconv’s computational efficiency to perform a systematic scale-space analysis that would be infeasible with existing methods: at 350,000 spots (8 *µ*m resolution), probabilistic methods face prohibitive barriers—Cell2Location cannot handle datasets beyond ~20,000 spots [13], and Stereoscope’s >8-hour runtime on 10,000 spots [12] would extrapolate to over 11 days.

Using Visium HD data from mouse small intestine (10x Genomics), we characterized data quality from the native 2 *µ*m bin size through 4, 8, and 16 *µ*m to establish the feasible range for sequencing-based deconvolution (Supplementary Note; Supplementary Table S4). At 2 *µ*m, each bin captures a median of only 17 UMI (16 genes detected), with 99.9% of expression matrix entries equal to zero and marker genes for even abundant cell types detected in fewer than 8% of bins. At 4 *µ*m (2 × 2 aggregation, median 78 UMI), 40% of bins reach the 100-UMI quality threshold. At 8 *µ*m (median 325 UMI), 84% of bins carry sufficient signal for stable compositional inference (Fig. 6a,b). At current sequencing depths and for a 10-cell-type compositional model, 8 *µ*m is thus the finest bin size at which most bins carry sufficient signal for stable independent deconvolution in this tissue; 4 *µ*m is marginally viable with spatial regularization (Supplementary Note), while finer bins require cell segmentation approaches [46–48] that integrate spatial morphology rather than compositional regression.

**Fig. 6.**
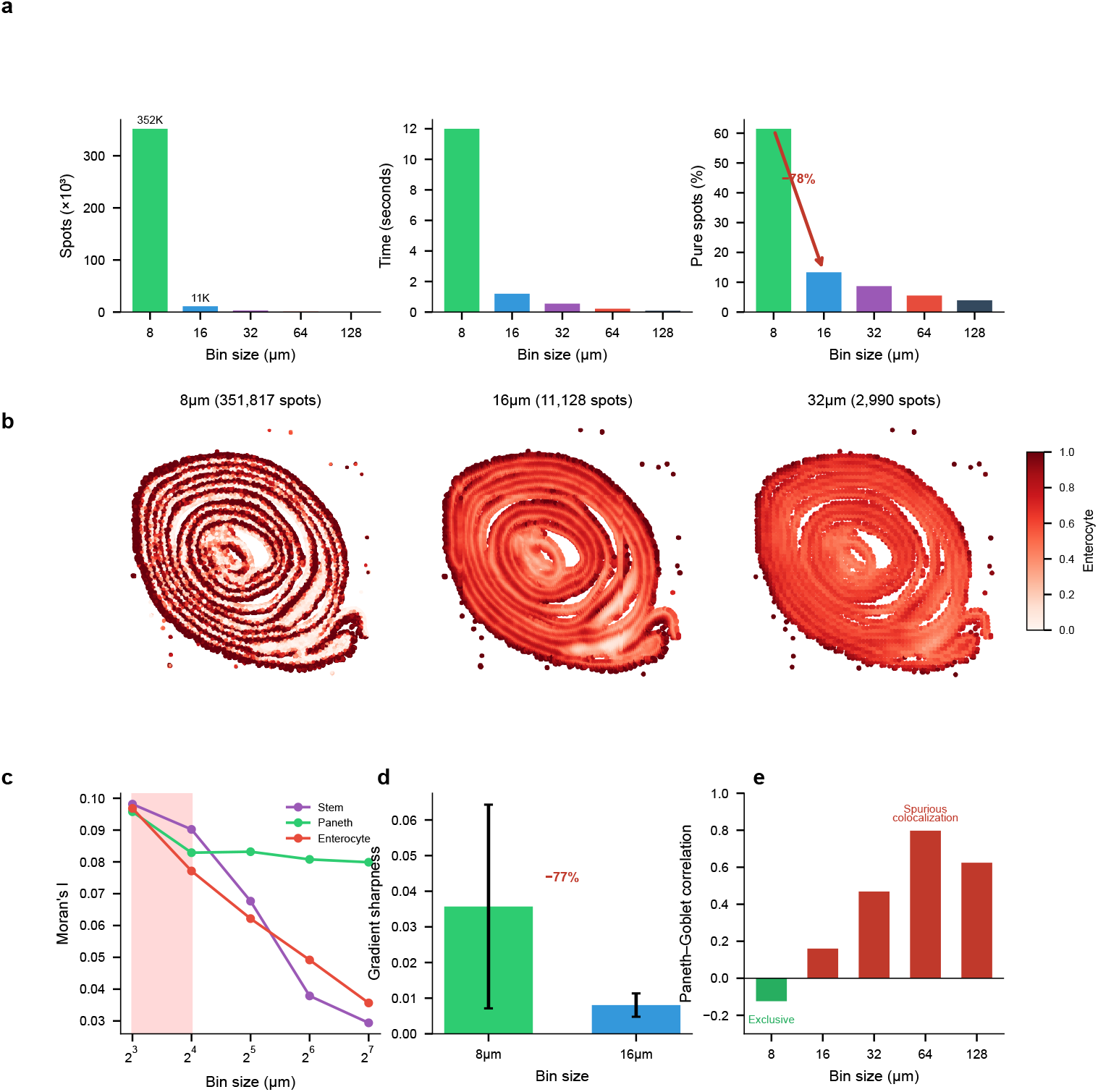
The resolution horizon: sequencing depth and spatial averaging jointly constrain Visium HD deconvolution. **(a)** Median UMI per bin across resolutions (error bars: IQR). Dashed line: 100-UMI quality threshold. At the native 2 *µ*m bin size, median UMI is 17—far below the threshold for stable 10-cell-type compositional inference. **(b)** Marker gene detection rate. Lyz1 (Paneth) and Muc2 (Goblet) are detected in *<*8% of 2 *µ*m bins; rare markers Lgr5 (stem) and Dclk1 (Tuft) are essentially absent (*<*0.05%). **(c)** Signal purity (blue circles, left axis) and normalized Shannon entropy (orange squares, right axis) across the full 2–128 *µ*m range. Purity peaks at 4–8 *µ*m and declines at both finer (data sparsity) and coarser (spatial averaging) resolutions; entropy mirrors this pattern. Gray band highlights the 8 *µ*m operating point. **(d)** Spatial maps of enterocyte proportions at 8, 16, and 32 *µ*m. Fine anatomical detail visible at 8 *µ*m becomes progressively blurred at coarser resolutions. **(e)** Resolution sensitivity varies by cell type. Stem cells show the steepest decline in spatial coherence (Moran’s I), while Paneth cells retain structure. Shaded region: 8–16 *µ*m transition. **(f)** Paneth–Goblet correlation across the full 2–128 *µ*m range. Weak mutual exclusion at 2–8 *µ*m (green bars) flips to spurious positive correlation by 64 *µ*m (red bars)—a qualitative reversal in the apparent biological relationship. Data: Visium HD Mouse Small Intestine (10x Genomics), scRNA-seq reference from Haber et al. 2017. Extended QC characterization in Supplementary Note and Supplementary Table S4.

Starting from this 8 *µ*m operating point, we deconvolved cell type proportions across five resolutions (8, 16, 32, 64, and 128 *µ*m) using a matched scRNA-seq reference [49]. FlashDeconv processed 351,817 bins at 8 *µ*m resolution in 12.0 seconds— a throughput of approximately 29,000 bins per second on commodity hardware. The complete five-resolution analysis (366,975 total bins) required only 14 seconds, enabling systematic exploration of the resolution-information trade-off.

#### The resolution horizon

Our analysis revealed a pronounced transition between 8 *µ*m and 16 *µ*m resolution (Fig. 6c). At 8 *µ*m, 61.5% of bins were dominated by a single cell type (proportion >80%), consistent with near-single-cell purity. This fraction collapsed to 13.3% at 16 *µ*m—a 78% reduction—and continued declining to 3.9% at 128 *µ*m. The normalized Shannon entropy of cell type mixing increased correspondingly, from 0.23 at 8 *µ*m to 0.48 at 16 *µ*m. This transition represents the boundary beyond which cellular identity becomes fundamentally obscured by spatial averaging. The 8–16 *µ*m transition observed in mouse small intestine was independently reproduced in mouse colon using Xenium ground truth without deconvolution (Supplementary Note 4), suggesting this scale is a robust feature of intestinal epithelium. Notably, this transition coincides with the characteristic diameter of intestinal epithelial cells (~10–15 *µ*m), consistent with the geometric intuition that the resolution horizon occurs when measurement bins transition from capturing predominantly single cells to averaging multiple cells. For tissues with substantially different cell sizes or spatial organization patterns, the resolution horizon would be expected to shift accordingly—highlighting that this is a tissue-dependent biophysical parameter rather than a universal constant.

#### Cross-tissue validation: mouse brain cortex

To test this prediction, we extended the analysis to mouse brain cortex (Visium HD, 453,820 bins; Supplementary Note). At the broad lineage level (excitatory and inhibitory neurons, astrocytes, oligodendrocytes, OPCs, microglia, vascular cells, neuroblasts), brain tissue exhibited a qualitatively different profile: purity declined gradually from 0.54 to 0.36 across 8–128 *µ*m with no inflection comparable to the intestinal cliff, and 12 of 28 lineage pairs showed correlation sign flips, including astrocyte–microglia (*r* = −0.12 → +0.51; Supplementary Fig. S33). Independent MERFISH ground truth from the Allen Brain Cell Atlas (115,772 cells, no deconvolution) confirmed the gradual degradation and reproduced sign flips (Supplementary Fig. S34), establishing that the resolution horizon is shaped by tissue architecture.

#### Cell types exhibit distinct resolution sensitivities

Different cell types showed markedly different vulnerabilities to spatial aggregation (Fig. 6e). Intestinal stem cells—localized to crypt bases with a characteristic niche size of approximately 10–15 *µ*m—exhibited the steepest information loss: spatial autocorrelation (Moran’s I) decreased by 70% from 8 *µ*m to 128 *µ*m. In contrast, Paneth cells, which form tight clusters at crypt bases, showed only 17% loss of spatial coherence across the same resolution range. These differences reflect the characteristic spatial scales of each cell type’s tissue organization.

#### Validation through crypt-villus boundary analysis

To confirm that the observed resolution effects reflect genuine biological structure, we analyzed the sharpness of crypt-villus boundaries—the transition zone between Paneth cell-rich crypts and enterocyte-covered villi (Supplementary Note). We traced 100 linear paths from crypt cores (Paneth >50%) to villus regions (enterocyte >70%) and computed the maximum gradient of enterocyte proportion along each path. At 8 *µ*m, the mean gradient sharpness was 0.036 ± 0.029. At 16 *µ*m, this decreased to 0.008 ± 0.003—a 77% reduction in boundary definition. At 32 *µ*m and coarser, crypt cores meeting our criteria could no longer be identified. This quantitative boundary analysis confirms that the information loss corresponds to anatomical blurring rather than algorithmic artifacts.

#### Coarse binning induces spurious colocalization

Beyond signal loss, spatial aggregation can fundamentally distort cell-cell relationship inference (Fig. 6f). Paneth cells and Goblet cells are both secretory epithelial types, yet occupy distinct spatial niches: Paneth cells reside exclusively at crypt bases, while Goblet cells distribute along the crypt-villus axis. At 8 *µ*m resolution, this spatial segregation manifests as a weak negative correlation (*r* = −0.12, *p* < 10^−100^, *N* = 351,817), reflecting their mutual exclusion at the cellular scale. However, as resolution coarsens, both cell types become mixed within the same measurement bins, inducing a spurious positive correlation that peaks at 64 *µ*m (*r* = +0.80). This correlation sign flip—from negative to positive—represents a qualitative reversal in the apparent cell-cell relationship, a tissue-scale instance of the modifiable areal unit problem (MAUP) from spatial statistics [50], recently highlighted in the context of spatial transcriptomics [51]. The mechanism is geometric: at fine resolution, adjacent but distinct cell populations are resolved separately, but as bin size increases beyond cells’ physical dimensions, they become averaged into the same measurement unit and appear artificially co-localized. Researchers analyzing data at conventional Visium resolution (55 *µ*m) would observe strong colocalization and might erroneously conclude that Paneth and Goblet cells share a common microenvironment or respond to similar spatial cues. To confirm that this phenomenon reflects spatial geometry rather than deconvolution methodology, we performed ground truth validation using Xenium in situ sequencing data from mouse colon, where exact cell positions are known without deconvolution. The ground truth itself exhibits the same pattern: spatially adjacent but distinct cell clusters show negative correlation at 8 *µ*m (*r* = −0.06) but strong positive correlation at 128 *µ*m (*r* = +0.68), confirming that the observed sign reversal reflects tissue geometry rather than data sparsity or deconvolution methodology (Supplementary Note 4; Supplementary Fig. S16). FlashDeconv’s ability to process high-resolution data efficiently enables detection of such resolution-dependent artifacts, revealing the true spatial organization that coarse binning obscures.

#### Leverage scores explain preservation of rare cell signals

To verify the mechanistic connection between FlashDeconv’s mathematical design and the biological findings above, we confirmed that marker genes for the key cell types exhibit high leverage scores in the intestine reference (Supplementary Fig. S15). The canonical stem cell marker *Lgr5* ranks in the top 1% by leverage (rank #283 of 27,998 genes), despite ranking only in the top 10% by variance. Overall, stem cell markers average the 95th percentile by leverage, with 5 of 6 markers in the top 5%. This confirms that the 8 *µ*m stem cell niche detection demonstrated above is a direct consequence of leverage-weighted sketching: rare cell type signals are preserved during dimensionality reduction precisely because their markers define unique directions in gene expression space, regardless of their overall expression magnitude.

#### FlashDeconv resolves the Tuft-Stem chemosensory niche

Beyond validating marker preservation, FlashDeconv enabled niche-resolution quantification of cell-type co-localization directly from capture-based spatial transcriptomics—recovering a spatial architecture consistent with known Tuft cell biology [52, 53] through deconvolution of sequencing-based platforms without requiring imaging-based cell segmentation. Tuft cells (labeled “brush cell” in the Haber et al. reference annotation [49])—rare chemosensory epithelial cells comprising only ~0.4– 2% of intestinal epithelium [54]—exhibited the highest “HVG blindness” among all cell types in our analysis (Fig. 7a): their marker genes (including *Pik3r5, Ptgs1*) rank 21 percentile points lower under variance-based selection than under leverage-based ranking, placing them at highest risk of being discarded by standard feature selection pipelines. At 8 *µ*m resolution, FlashDeconv identified 2,244 focal Tuft cell niches with proportions reaching 61%—near-pure Tuft cell spots (Fig. 7b). These niches exhibited a striking spatial pattern: a 16.2-fold enrichment for intestinal stem cells, and 2.8-fold enrichment for enteroendocrine cells (Fig. 7c,e). Conversely, differentiated cell types were strongly depleted: enterocytes (0.20×), goblet cells (0.14×), and Paneth cells (0.15×)—all significantly non-random (permutation test, *p* < 10^−3^, *n* = 1,000). This Tuft-Stem co-localization is anatomically consistent with Tuft cells’ known localization near the intestinal stem cell zone at the crypt base [52, 53] and their recently discovered capacity to function as reserve stem cells following epithelial injury [55]. Notably, exploratory ligand-receptor analysis revealed that *Il17ra*—encoding a subunit of the IL-25 receptor through which Tuft cells signal to neighboring cells [56]—is expressed 7-fold higher in Tuft-Stem niches than in background tissue (Supplementary Fig. S21), suggesting that this spatial proximity may facilitate paracrine communication. Null model validation confirms this signal is biological rather than artifactual: independent marker gene sets show consistent spatial patterns (*r* = 0.30, *p* < 10^−200^), co-localization is specific to stem cells (not random cell types), Tuft cell distribution exhibits significant spatial autocorrelation (Moran’s I = 0.44 vs. random baseline ~0), and the Tuft-Stem proximity is recapitulated directly in raw gene expression without deconvolution (*Pou2f3* –*Lgr5* neighbor enrichment, *p* < 0.001; Supplementary Note 6, Supplementary Fig. S20). The Tuft cell signal is exquisitely resolution-dependent: maximum Tuft proportion decreased from 61% at 8 *µ*m to 4% at 128 *µ*m (Fig. 7d), rendering these niches undetectable at conventional Visium resolution. This finding illustrates that resolving such niche architecture from capture-based spatial transcriptomics requires overcoming two independent barriers: rare cell markers lost to variance-based feature selection, and spatial signal diluted at resolutions exceeding the niche’s characteristic scale (~10–15 *µ*m).

**Fig. 7.**
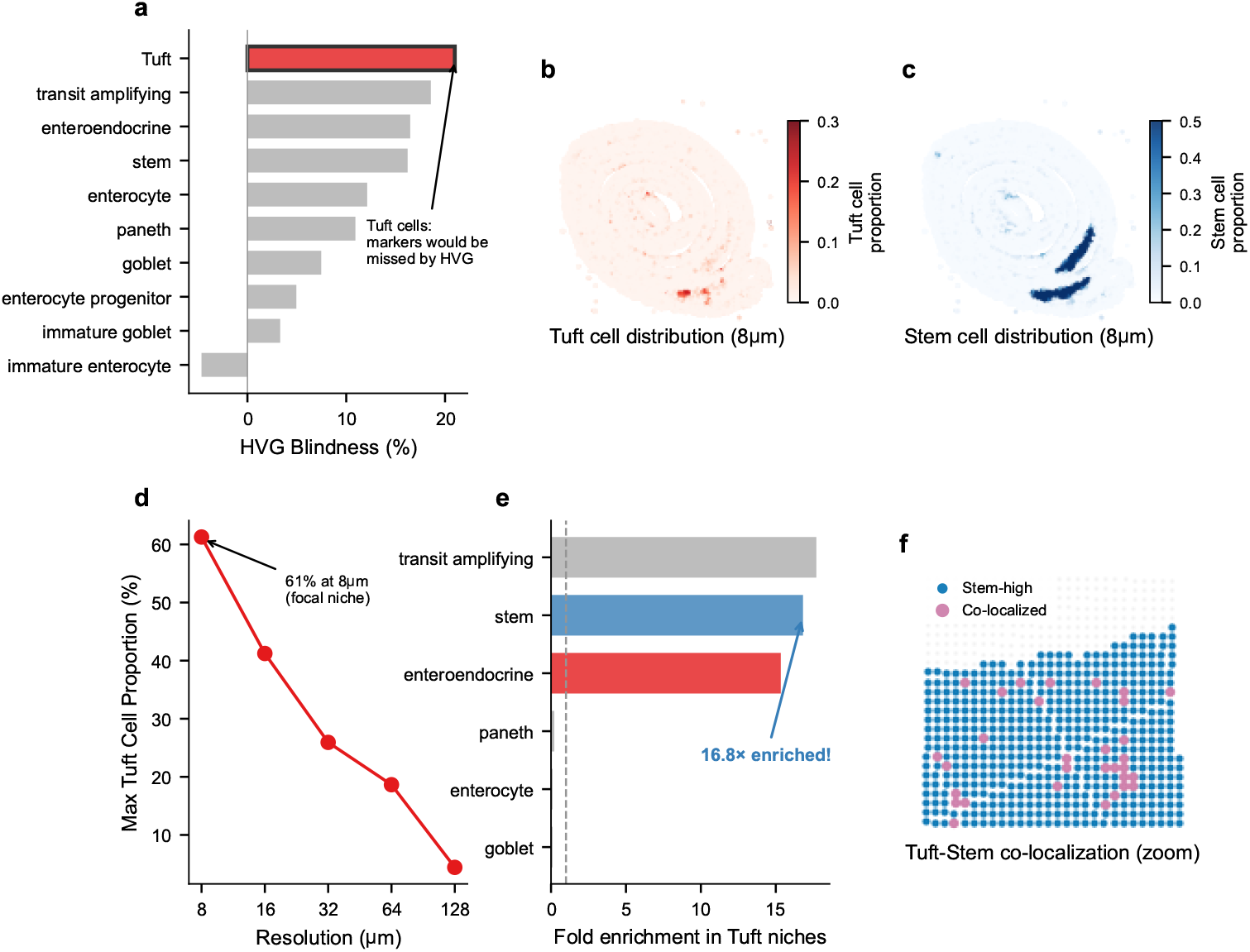
FlashDeconv resolves the Tuft-Stem chemosensory niche. **(a)** HVG blindness ranking across intestinal cell types. HVG blindness is defined as the difference in mean percentile rank of a cell type’s marker genes under variance-based versus leverage-based selection; positive values indicate systematic underweighting by HVG. Tuft cells exhibit the highest HVG blindness (21 percentile points). **(b)** Spatial distribution of Tuft cells at 8 *µ*m resolution reveals focal niches (red spots) with proportions up to 61%. **(c)** Stem cell distribution at 8 *µ*m shows concentration at crypt bases. **(d)** Resolution sensitivity of Tuft cell detection. Maximum proportion decreases from 61% (8 *µ*m) to 4% (128 *µ*m), rendering focal niches undetectable at conventional resolution. **(e)** Co-localization analysis reveals Tuft cell hotspots are enriched 16.2-fold for stem cells and 2.8-fold for enteroendocrine cells (*p <* 10^−3^, permutation test), but depleted for differentiated cell types (enterocytes 0.20×, goblet cells 0.14×). **(f)** Spatial zoom showing Tuft-Stem co-localization at crypt bases (blue: Stem-high, pink: co-localized). Tuft cells rarely appear without adjacent stem cells, consistent with their intimate niche association. Data: Visium HD Mouse Small Intestine (10x Genomics).

#### Cross-method comparison confirms niche architecture

To assess whether this spatial architecture reflects genuine tissue biology or is specific to FlashDeconv’s design, we applied RCTD—a likelihood-based method that treats each spot independently—to the same tissue at 16 *µ*m resolution, the finest scale at which RCTD’s per-spot inference remains practical on this tissue (10,952 spots; 8 *µ*m would require >350,000 independent regressions). Both methods detected stem cell enrichment in Tuft cell hotspots, confirming that the niche reflects tissue architecture rather than a regularization artifact. However, FlashDeconv produced markedly more spatially coherent assignments, consolidating Tuft cell signal into 9 clusters versus 168 scattered fragments for RCTD, with 7.7-fold stem cell enrichment at hotspots compared to 3.0-fold (Supplementary Fig. S26). This difference partly reflects FlashDeconv’s graph Laplacian regularization, which borrows information from spatial neighbors to enforce local continuity. To verify that this coherence corresponds to genuine biology rather than over-smoothing, we examined *Il17ra*—encoding a subunit of the IL-25 receptor critical for Tuft-Stem paracrine signaling [56]—as a functional readout independent of both deconvolution methods. FlashDeconv-defined niche zones showed 5.2-fold *Il17ra* enrichment over background versus 1.5-fold for RCTD-defined zones, indicating that the spatially coherent assignments more accurately delineate functional signaling microenvironments (Supplementary Note 8).

## Discussion

In this study, we present FlashDeconv, a deconvolution framework designed for atlas-scale spatial transcriptomics. By combining leverage-score importance sampling with sparse graph regularization, FlashDeconv achieves accuracy comparable to probabilistic methods while reducing computational cost by orders of magnitude. As introduced above, leverage scores have been used across computational biology for both cell subsampling [16, 17] and gene-space feature identification via CUR decomposition [57] and variable screening [58]. FlashDeconv extends this principle by integrating leverage-score importance weights directly into a CountSketch randomized projection for spatial deconvolution—a gene-space compression operation distinct from the cell-subsampling approach used in Seurat v5’s sketch analysis [17]. Despite the inherent randomness in sketching, FlashDeconv produces highly reproducible results: across 10 runs with different random seeds, pairwise Pearson correlations exceeded 0.98, confirming that users can obtain consistent results without fixing random seeds.

The core insight underlying our approach is the distinction between geometric structure and statistical variance. Variance-based methods—including PCA and highly variable gene selection—conflate biological signal with population frequency, systematically underweighting rare populations regardless of their biological importance. Leverage scores instead capture each gene’s contribution to the discriminative structure among cell types, independent of expression magnitude or population size. Our experiments confirm this decoupling: leverage ranking remains stable under 67-fold changes in cell type abundance, and the structurally informative (GOLD) gene set shows coherent functional enrichment and clear spatial organization, while variance-dominated (NOISE) genes lack both—indicating that high variance alone does not ensure biological relevance.

Our choice of Log-CPM normalization reflects an engineering trade-off rather than a claim of statistical superiority. While Pearson residuals are statistically principled for negative binomial count data, Log-CPM provides specific advantages for *L*_2_-based sketching: its bounded norm prevents high-expression genes from dominating the compressed space. This pragmatic choice, combined with leverage-weighted sketching, allows FlashDeconv to preserve essential biological signals during dimensionality reduction.

Our case studies reveal three complementary failure modes in spatial deconvolution that FlashDeconv addresses through distinct mechanisms. The melanoma dataset represents a feature-space challenge: highly correlated malignant states (*r* > 0.98) along a continuous phenotypic spectrum create an ill-conditioned regression problem. Here, sparse NNLS naturally selects the dominant signals present in the tissue without requiring manual reference curation—a property particularly valuable for atlas-scale mapping where exhaustive reference curation is impractical. We note that this favorable behavior depends on sparsity: when correlated subtypes are both present at appreciable abundance and must be individually quantified, signature collinearity poses a general identifiability challenge for linear regression frameworks—as we demonstrate quantitatively for the endothelial triplet in liver (cosine similarity >0.975; Supplementary Note) and for macrophage subtypes in breast cancer (Supplementary Note). The Visium HD mouse intestine dataset represents a signal-space challenge: at 8 *µ*m resolution with sparse counts, rare cell markers risk being discarded during dimensionality reduction. Leverage-weighted sketching preserves these geometrically distinctive signals, enabling detection of the Tuft-Stem chemosensory niche—spatial architecture that requires both rare-marker-aware feature selection and sub-16 *µ*m resolution to resolve. Cross-method comparison with RCTD confirms that the niche is detectable by either approach, but highlights the role of spatial regularization in resolving niche architecture: spot-independent methods that treat each location in isolation disperse the signal across the tissue, producing niche zones with weaker enrichment for the paracrine signaling gene *Il17ra* than spatially regularized estimates (Supplementary Note 8). This intimate spatial proximity between Tuft cells and intestinal stem cells aligns with recent findings that Tuft cells can dedifferentiate into functional stem cells during regeneration [55], suggesting that the niche architecture we observe may facilitate both paracrine signaling and direct cellular interconversion. Quantitatively, our measurements—16.2-fold stem cell enrichment within Tuft cell foci at ~10–15 *µ*m resolution—are consistent with lineage tracing and imaging studies that localize Tuft cells to the Lgr5-positive crypt base compartment [52, 53] and with the spatial range required for IL-25 paracrine signaling [56]. FlashDeconv’s contribution here is not the discovery of this proximity per se, but its quantitative characterization from capture-based spatial transcriptomics, showing that this architecture can be recovered from sequencing-based spatial data and that sub-16 *µ*m resolution is required to resolve it. The CRC Visium HD analysis reveals a third challenge specific to high-resolution platforms: continuous cell type mixing within measurement bins creates a fundamental mismatch with discrete classification. RCTD’s doublet mode—which allows up to two cell types per bin—provides confident singlet labels for 46–59% of bins and withholds 5–7% at its quality threshold, a conservative design appropriate for single-cell classification but one that limits spatial coverage in the continuous-mixing regime. FlashDeconv’s regression framework produces continuous proportions for every bin, and independent marker gene validation confirms accuracy in >86% of disputed assignments. The analytical consequence of this output-type difference is illustrated by the neutrophil microdomain analysis (Fig. 5): RCTD’s doublet mode classifies 2.3% of the 16,827 hotspot bins as Neutrophil singlets, leaving the organized niche structure—neutrophils co-localized with immunoregulatory mRegDC [44], macrophages, mast cells, and endothelial cells at vascularized stromal sites—difficult to summarize through discrete classification alone. This tension between continuous mixing and sparse per-bin counts intensifies as spatial transcriptomics moves toward finer resolution: methods without sparsity regularization must fall back to discrete classification to avoid underdetermined estimation. FlashDeconv’s *ℓ*_1_ penalty and graph Laplacian smoothing keep continuous regression well-posed across all *K* cell types without restricting the number of types per bin. No single mechanism addresses all three challenges; FlashDeconv’s combination of structure-preserving compression, signal-driven inference, and regularized continuous regression provides robustness across these qualitatively different failure modes.

The neutrophil-mRegDC co-localization is notable in light of recent characterizations of mRegDC spatial niches in tumor immunity. You et al. [59] identified a peri-lymphatic Treg-mRegDC niche in which mRegDC facilitate regulatory T cell activation, constraining antigen trafficking to draining lymph nodes. The niche we report occupies a distinct anatomical compartment—the vascularized tumor–stroma interface—and re-analysis of the Marteau et al. Xenium CRC atlas [60] (15 patients, 3.7 million cells) directly confirms the spatial relationship at single-cell resolution without deconvolution: LAMP3^+^ mRegDC are enriched within 50 *µ*m of neutrophil clusters in 10 of 15 patients (median 1.49-fold, permutation *P* < 0.05), with macrophages enriched in 14 of 15 patients and tumor cells depleted—recapitulating the niche architecture inferred by FlashDeconv (Supplementary Note 9). Cross-modality concordance supports broader generality: granulocytes and mRegDC co-occur in 53 of 60 patients in the Pelka scRNA-seq atlas [61] (*ρ* = 0.44, *P* = 4 × 10^−4^), and a neutrophil–mRegDC transcriptomic signature correlation persists across 592 TCGACOAD patients (*ρ* = 0.52, *P* = 2.5 × 10^−42^; Supplementary Note 9). Given that mRegDC abundance predicts immunotherapy response in independent cohorts [62], these observations generate a testable hypothesis: that neutrophil extravasation at the tumor–stroma interface may promote the mRegDC regulatory program, serving as an upstream source for the immunoregulatory dendritic cells that populate perilymphatic suppressive niches [59]. This spatial cascade would link low-UMI bins at sites of immune cell extravasation—where discrete labeling is least complete—to a clinically relevant immunoregulatory pathway (Supplementary Note 9).

Beyond accuracy, FlashDeconv’s computational efficiency enables systematic scale-space exploration. The phenomenon of resolution-dependent information loss upon spatial aggregation—including correlation sign reversal—is well established in spatial statistics as the modifiable areal unit problem (MAUP) [50], and has recently been identified as relevant to spatial transcriptomics [51]. Our multi-resolution analysis provides systematic, tissue-specific quantification of this effect in high-resolution spatial transcriptomics, identifying what we call the *resolution horizon*: the spatial scale at which biological segregation collapses into geometric mixing. Signal purity drops precipitously between 8 and 16 *µ*m, while cell-cell correlations undergo sign inversion (from *r* = −0.12 to *r* = +0.80 by 64 *µ*m), as validated in Xenium ground truth. Systematic characterization from 2 to 16 *µ*m further reveals that 8 *µ*m is bounded from below as well: at 2 *µ*m, bins contain a median of 17 UMI—too sparse for stable compositional estimation—establishing 8 *µ*m as the finest resolution at which, for this tissue and sequencing depth, most bins carry sufficient signal for stable compositional inference without spatial smoothing (Supplementary Note). Critically, the resolution horizon is both tissue-specific and cell-type-dependent. Within mouse intestine, stem cells require 8 *µ*m resolution while Paneth cells tolerate 32 *µ*m. Across tissues, the horizon manifests as a sharp transition in organized epithelium but as gradual degradation in brain cortex, where cell types are highly intermingled—a contrast independently confirmed by Xenium and MERFISH ground truth (Supplementary Note). This cross-scale, cross-tissue granularity is not captured by prior theoretical treatments of MAUP. FlashDeconv’s linear scalability transforms resolution selection from an arbitrary experimental choice into a data-driven determination.

Our sparse graph Laplacian regularization provides coverage-dependent benefits particularly relevant for emerging high-resolution platforms (Supplementary Note). Systematic ablation across 76 cell types confirms that spatial regularization improves rare cell type detection (AUPRC) despite a mild precision–recall trade-off: the dominant source of false positives is probability mass competition in the NNLS framework, not Laplacian propagation (Supplementary Fig. S10). At the sub-8 *µ*m depths characterized above, individual bins carry insufficient signal for independent compositional inference, and spatial regularization becomes essential for borrowing strength from neighboring measurements. As spatial transcriptomics scales to finer resolution with increasingly sparse measurements, this adaptive regularization becomes increasingly important.

A complementary question is whether compositional deconvolution provides value over simpler alternatives. Recent work has shown that differential expression marker gene scoring can achieve competitive rare cell type detection in certain settings [63], and hybrid approaches such as SMART [64] integrate marker gene priors into topic models to bridge scoring and regression paradigms. We systematically compared FlashDeconv against marker gene scoring (top-50 DE markers per type, mean-expression scoring with background subtraction) across all 76 cell types in the six Spotless tissues. On multi-cell spots, FlashDeconv achieves higher accuracy than marker scoring across all abundance categories, with the largest advantage for rare types (Pearson *r* = 0.902 vs. 0.724, Δ = +0.178; AUPRC 0.919 vs. 0.792, Δ = +0.127; Supplementary Note). Xenium ground-truth validation corroborates this on CRC tissue: at 4 *µ*m, FlashDeconv achieves AUPRC 0.87 versus 0.62 for NNLS, 0.60 for marker scoring, and 0.30 for RCTD (full mode; its internal UMI filter removes 73% of 4 *µ*m bins). The gap is largest for cell types central to the niche analyses above (mRegDC: 0.62 vs. 0.21 for marker scoring; Neutrophil: 0.53 vs. 0.16; Supplementary Table S3). This gap reflects a structural difference: marker scoring evaluates each type independently and cannot model compositional constraints (proportions summing to 1), while deconvolution jointly estimates all types within each spot, leveraging the shared variance structure to resolve overlapping signatures. For downstream analyses requiring proportions—co-localization statistics, niche composition, cell–cell interaction quantification—deconvolution provides the compositional output that scoring does not.

We acknowledge scenarios where alternative approaches may be preferable. For small datasets (<50 spots), sketching and spatial smoothing provide limited benefit. For resolving fine-grained cell states along continuous phenotypic spectra, probabilistic models with full posterior inference remain valuable; however, the Laplace approximation described above provides per-spot confidence intervals that capture the dominant sources of estimation uncertainty at a fraction of the computational cost (Supplementary Note). More broadly, as recent reviews emphasize [4], spatial transcriptomics analysis encompasses distinct computational tasks—deconvolution of multi-cell measurement units, classification of individual cells, and spatial domain identification— each requiring different methodological assumptions. When the analytical task reduces to classifying individual cells—as in single-cell-resolution platforms such as Xenium— rather than deconvolving multi-cell spots, dedicated classification methods (e.g., TopACT [65]) or doublet models are better suited for distinguishing closely related subtypes that share the majority of their expression programs, because spatial regularization can propagate signals across cell boundaries in this regime (Supplementary Note). The biological analyses in this work—resolution-dependent co-localization sign reversal, spatial proportion gradients, and niche composition quantification—all operate on continuous proportions; discrete cell-type labels, whether from classification or doublet models, cannot serve as input for these computations. When platform-specific batch effects dominate biological signal, methods with explicit noise modeling may be more robust, although leverage scores provide inherent robustness to batch effects through reference anchoring (Supplementary Note). A related limitation is reference incompleteness: when the spatial tissue contains cell types absent from the scRNA-seq reference, deconvolution estimates are necessarily biased. However, FlashDeconv’s unnormalized regression framework provides an implicit out-of-distribution detection mechanism through reconstruction residuals, which can flag regions of potential reference incompleteness (Supplementary Fig. S19). A subtler form of mismatch arises when the same cell type exhibits different marker gene rankings across modalities—for example, a gene that is a top marker in dissociated scRNA-seq may be less discriminative in spatial platforms due to differences in capture efficiency, gene panel coverage, or tissue-context-dependent expression. Because FlashDeconv’s leverage scores are derived from the reference signature matrix, such cross-modality inconsistency can reduce weighting accuracy for affected genes. Two design features partially mitigate this: the HVG selection stage draws from the spatial data itself, providing a data-driven filter that downweights genes absent or uninformative in the spatial modality, and the Log-CPM normalization reduces platform-specific dynamic range differences. Empirically, FlashDeconv ranks 1st out of 13 methods on the Spotless liver reference sensitivity test—which evaluates consistency across exVivo scRNA-seq, inVivo scRNA-seq, and snRNA-seq references—suggesting robustness to moderate reference variation, though systematic evaluation across a broader range of platform mismatches remains an important direction for future work. We view FlashDeconv not as a universal replacement, but as a purpose-built tool for million-spot atlases where linear scalability is essential.

More broadly, this work illustrates that measuring geometric structure rather than statistical variance can decouple biological importance from numerical prevalence—a principle that may prove valuable in other computational biology settings where rare populations carry disproportionate functional significance. As spatial biology scales to increasingly fine resolution, feature selection strategies that preserve critical signals regardless of abundance will be essential to ensure computational efficiency does not compromise biological discovery.

## Methods

### Algorithm Overview

Algorithm 1 presents the complete FlashDeconv pipeline, which consists of five main stages: gene selection, data preprocessing, structure-preserving sketching, spatial graph construction, and optimization via block coordinate descent.

### Two-Stage Feature Selection

FlashDeconv employs a two-stage feature selection strategy that combines variance-based noise filtering with reference-guided marker inclusion, followed by leverage-based importance weighting during sketching. This hybrid approach addresses the inherent tension between noise reduction and signal preservation: purely variance-based selection (e.g., PCA on all genes) would carry forward technical noise and house-keeping genes, while purely leverage-based selection on raw noisy data could amplify measurement artifacts.

#### Algorithm 1

FlashDeconv: Structure-Preserving Spatial Deconvolution

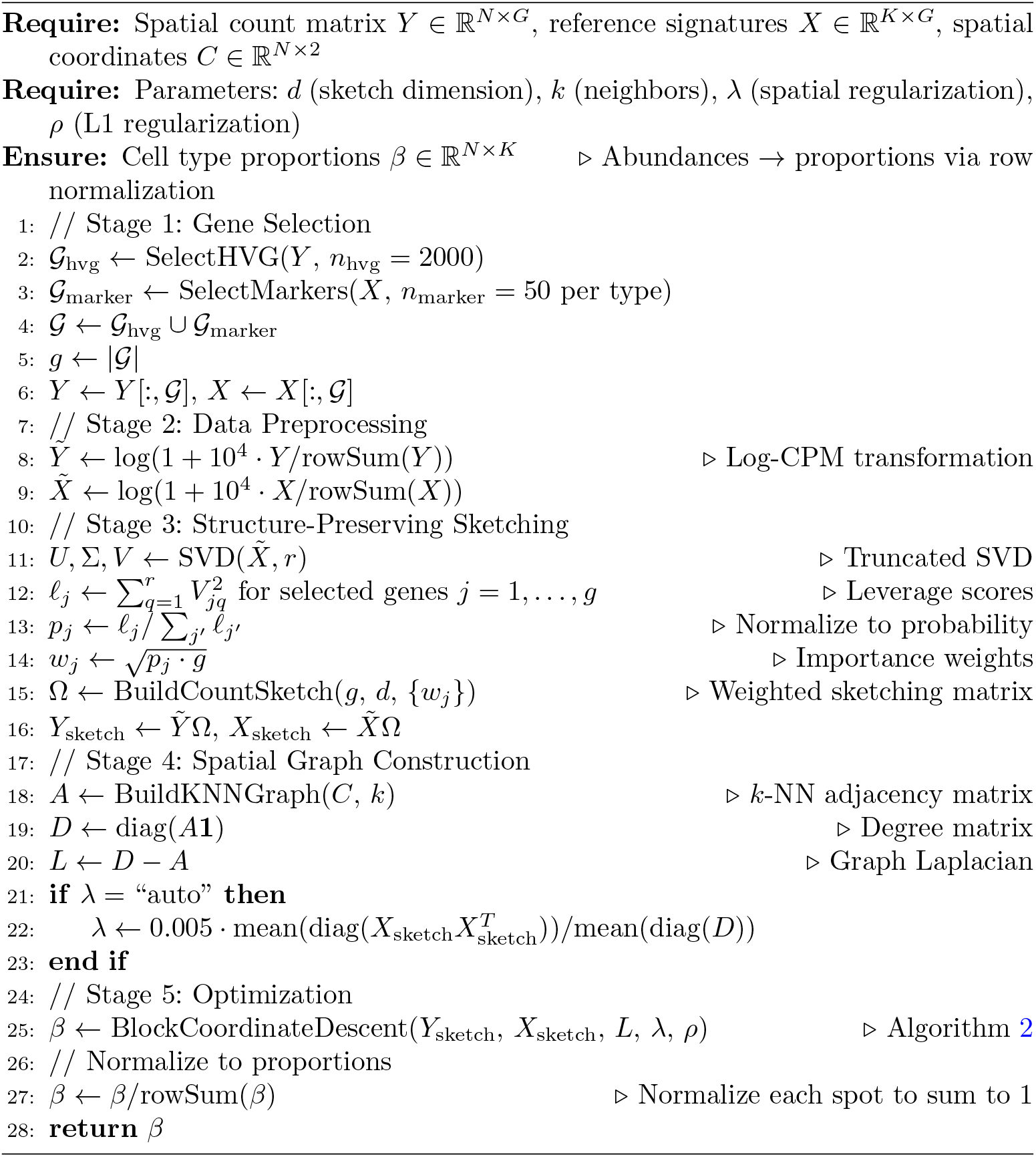

#### Stage 1: Variance-based noise filtering (HVG selection)

We first apply Highly Variable Gene (HVG) selection as a coarse filter to remove uninformative genes dominated by technical noise. Genes are binned by mean expression, and normalized dispersion is computed within each bin using the Seurat v3 method [37]. We select the top 2,000 genes with normalized dispersion exceeding 0.5 and mean expression between 0.0125 and 3.0 (log-scale). This stage removes housekeeping genes with near-constant expression and lowly-expressed genes dominated by dropout noise, reducing the input space from *G* ≈ 20,000 to *G*^*′*^ ≈ 2,000 genes. We validated this two-stage strategy by comparing it against whole-transcriptome leverage calculation (without HVG filtering): the two approaches showed 92.4% overlap in top-ranked leverage genes, with all discordant genes belonging exclusively to the high-variance NOISE quadrant (Supplementary Fig. S12). This confirms that HVG pre-filtering effectively removes technical artifacts without discarding biologically important high-leverage genes.

#### Stage 2: Reference-guided marker inclusion

Independently from HVG selection, we identify cell-type marker genes from the *whole transcriptome* of the reference matrix *X*. For each gene, we compute the expression gap 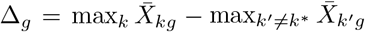 between its highest-expressing cell type and the runner-up. For each cell type, the top 50 genes maximizing Δ_*g*_ are selected as markers. We adopt this gap criterion rather than statistical differential expression (e.g., Wilcoxon rank-sum tests as implemented in Scanpy) because deconvolution solves a regression problem in the aggregated signature space *X* ∈ ℝ^*K*×*G*^, not at the single-cell level: what determines identifiability is the separation between cell type *mean profiles*, not the statistical significance of per-cell differences. The max-minus-second-max gap directly measures this separation, operates on the *K* × *G* signature matrix in *O*(*KG*) time without requiring access to individual cells, and is invariant to within-type heterogeneity that may inflate or deflate *p*-values without affecting the regression geometry. The final gene set is the union of HVGs and markers (*G* = *G*_hvg_ ∪ *G*_marker_), typically |*G*| ≈ 2,500 genes. Leverage scores are then computed on this combined gene set for structure-preserving sketching. Critically, the subsequent sketching step (Section 2) applies leverage-score-based importance sampling to this filtered gene set, ensuring that rare cell type markers are preserved in the final *d* = 512 dimensional sketch.

This two-stage design avoids the “variance-only” pitfall where rare cell signals are lost during dimension reduction, while simultaneously preventing leverage scores from being computed on noisy, uninformative genes. The combination of variance filtering, marker inclusion, and leverage-based sketching enables FlashDeconv to reduce the original *G* ≈ 20,000-gene transcriptome to *d* = 512 sketch dimensions without sacrificing rare cell type detection accuracy (Algorithm 1, lines 2–5, 11–13). Empirical validation on liver scRNA-seq data confirms that leverage-based importance sampling successfully enriches rare cell type markers by 2.5-fold compared to variance-based methods (Fig. 2).

### Data Preprocessing

We evaluated multiple variance-stabilizing transformations—including Pearson residuals commonly used for negative binomial data—and selected Log-CPM as the default. Our signal-to-noise analysis demonstrates that Log-CPM provides favorable properties for sketching-based methods by avoiding both the high-expression saturation and low-expression noise amplification inherent to Pearson residuals (Supplementary Note 2). After gene selection, let *Y* ∈ ℝ^*N* ×*g*^ be the selected spatial count matrix. We normalize by library size and log-transform:

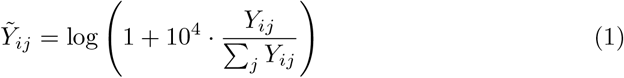

The selected reference matrix *X* ∈ ℝ^*K*×*g*^ is transformed identically to ensure the regression problem *Y* ≈ *βX* operates in a consistent feature space.

### Structure-Preserving Randomized Sketching

Randomized sketching compresses high-dimensional data via random projections— instead of computing expensive eigendecompositions (as in PCA), we randomly assign each selected gene to one of *d* lower-dimensional feature groups, achieving similar dimensionality reduction at a fraction of the computational cost. To reduce the dimensionality of the regression problem from *g* selected genes to *d* features (*d* ≪ *g*), we construct a sparse sketching matrix Ω ∈ ℝ^*g*×*d*^ using a leverage-score-weighted CountSketch transform [66].

First, we compute the statistical leverage scores *ℓ*_*j*_ from the singular value decomposition of the transformed selected reference 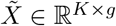:

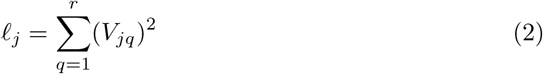

where *V* ∈ ℝ^*g*×*r*^ contains the right singular vectors of 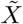 (corresponding to selected genes) and *r* is the numerical rank. Genes with high leverage scores correspond to discriminative markers that distinguish between cell types; for example, in liver scRNA-seq data, *Rspo3* achieves the highest leverage score among evaluated genes as a marker of Central Vein Endothelial cells (2% of cells), while abundant Hepatocyte markers exhibit lower leverage due to transcriptional overlap with other lineages.

We then construct Ω as follows. For each selected gene *j*, we draw an independent uniform random assignment *h*(*j*) ~ Uniform {1, …, *d*}, placing that gene in exactly one sketch dimension and setting all other entries to zero:

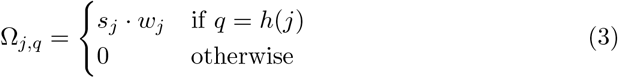

where *s*_*j*_ ∈ {−1, +1} is a random sign (Rademache r variable) and 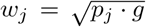 is a leverage-based importance weight, with *p*_*j*_ = *ℓ*_*j*_*/ ∑*_*j*_′ *ℓ*_*j*_′ denoting the normalized leverage probability over the selected gene set. This weighting ensures that high-leverage genes (often marker genes with low total abundance but high discriminative power) contribute proportionally more to the sketch, preserving their signal in the low-dimensional projection. This amplification is critical because CountSketch hashes each gene to exactly one of *d* dimensions: when a rare cell marker and a high-abundance housekeeping gene collide (are assigned to the same dimension), their contributions are summed, and without importance weighting the abundant gene’s magnitude can overwhelm the marker signal. Leverage-proportional scaling provides asymmetric amplification—empirically, structurally informative genes (low variance, high leverage) exhibit 3–12× higher leverage-to-magnitude ratios than variance-dominated genes across six tissues (mean 6.2×; Fig. 2b)—ensuring that rare cell type signals survive hash collisions while technical noise is relatively attenuated. This selective weighting mitigates hash collision variance [32] by maintaining adequate signal-to-interference ratio in the compressed space. Finally, columns of Ω are normalized to have unit *ℓ*_2_ norm, scaled by 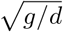 to approximately preserve Frobenius norms.

The data are then projected into the sketch space:

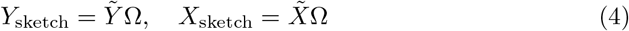

This allows us to solve the deconvolution problem in the *d*-dimensional sketch space (default *d* = 512; validated in Supplementary Fig. S1) with theoretically bounded approximation error [31, 67, 68]. Since *Y* primarily lies in the row space spanned by *X* (i.e., *Y* ≈ *βX* + *ϵ*), leverage scores computed from *X* effectively capture the geometric structure of both matrices, ensuring that Ω preserves not only *X* but also the projection of *Y* onto the biological subspace spanned by reference cell types.

#### Choice of weighting scheme

The leverage-proportional weight 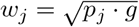 is motivated by the importance sampling theory underlying randomized matrix approximation [14, 15]: sampling or weighting proportional to leverage scores minimizes expected approximation error for the gene-space geometry of *X*. During development, we evaluated four alternatives: (i) uniform weights (*w*_*j*_ = 1, standard CountSketch), (ii) binary thresholding (retain only the top-*d* leverage genes with equal weight), (iii) variance-proportional weights 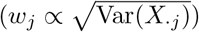, and (iv) a variance-leverage product 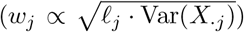. On the six-tissue Spotless benchmark, leverage-proportional weighting achieved the highest rare cell type Pearson correlation (0.649), followed by the variance-leverage product (0.618), uniform (0.451), variance-proportional (0.430), and binary thresholding (0.394). The advantage of continuous leverage weighting over thresholding reflects the gradual nature of leverage scores: hard cutoffs discard moderate-leverage genes that collectively contribute to distinguishing cell types with partially overlapping signatures. Full results are provided in Supplementary Fig. S30.

### Spatial Regularization and Optimization

#### Spatial Graph Construction

Given spatial coordinates *C* ∈ ℝ^*N* ×2^, we construct a *k*-nearest neighbor graph to encode spatial proximity. For each spot *i*, let *N*_*k*_(*i*) denote its *k* nearest neighbors under Euclidean distance. The binary adjacency matrix *A* ∈ {0, 1}^*N* ×*N*^ is defined as:

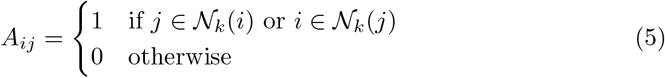

The symmetrization ensures an undirected graph. We use *k* = 6 by default, which matches the hexagonal packing geometry of standard Visium arrays; for square-grid platforms such as Visium HD, *k* = 6 captures the four cardinal plus two closest diagonal neighbors. Sensitivity analysis confirms that performance is robust across *k* ∈ {4, 6, 8, 12, 20} (Supplementary Fig. S2a, RMSE variation < 0.1%), making this default applicable across diverse spatial geometries. The graph Laplacian is then *L* = *D* − *A*, where *D* is the diagonal degree matrix with *D*_*ii*_ = ∑_*j*_ *A*_*ij*_.

#### Optimization Problem

We formulate the deconvolution task as a graph-regularized non-negative least squares problem:

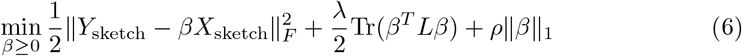

where ∥*β*∥_1_ = ∑_*i,k*_ |*β*_*ik*_| denotes the element-wise *ℓ*_1_ norm promoting sparsity.

Notably, we solve for unnormalized cell type abundances *β* ≥ 0 rather than directly optimizing proportions on the probability simplex (∑_*k*_ *β*_*ik*_ = 1). This formulation is better understood as *cellular density estimation*: *β*_*ik*_ represents the absolute abundance of cell type *k* at spot *i*, not its relative proportion. This approach offers three advantages: (1) it simplifies optimization to standard non-negative least squares; (2) the spatial Laplacian term encourages similar *total* cell densities across neighboring spots, a physically meaningful constraint; and (3) the unnormalized sum ∑_*k*_ *β*_*ik*_ implicitly captures spot-level capture efficiency variation, which would otherwise be forced into proportion estimates under simplex constraints (Supplementary Fig. S14). Cell type proportions are obtained via post-hoc row normalization, equivalent to maximum likelihood estimation under a multinomial model with unknown scale (Algorithm 1, final step).

#### Scale-Invariant Regularization

To ensure consistent regularization across datasets with varying signal magnitudes, both *λ* and *ρ* are internally scaled relative to the Gram matrix 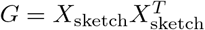. Let 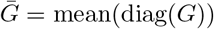 denote the mean Gram diagonal, which characterizes the data fidelity gradient scale. For spatial regularization:

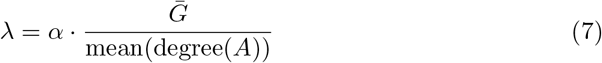

where *α* ∈ [0, 1) is a dimensionless tuning coefficient (default 0.005) that controls the relative contribution of the spatial term to the Hessian diagonal. For sparsity regularization, the partial residual *r*_*ik*_ in the BCD update scales as 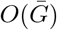, so the soft-thresholding parameter must be commensurate:

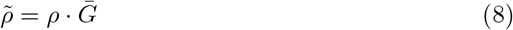

This makes the tuning coefficients *α* and *ρ* dimensionless fractions independent of sequencing depth or spot density; the resulting *λ* and 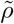 inherit the scale of the data automatically. Sensitivity analysis confirms robust performance across varying regularization strengths (Supplementary Fig. S9), including a per-cell-type ablation demonstrating that spatial regularization preserves rare cell type detection (Supplementary Fig. S10). The problem is solved using a fast Block Coordinate Descent (BCD) algorithm implemented in Python with Numba acceleration.

### Block Coordinate Descent Solver

We solve the optimization problem via coordinate descent with Jacobi-style parallel updates. At iteration *t*, updates read from *β*^(*t*)^ and write to a separate buffer *β*^(*t*+1)^, enabling row-wise parallelism across spots while keeping neighbor terms consistent within each iteration. Let 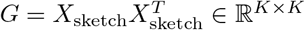 denote the precomputed Gram matrix. The partial derivative of the objective with respect to *β*_*ik*_ (ignoring the *ℓ*_1_ term and non-negativity constraint) is:

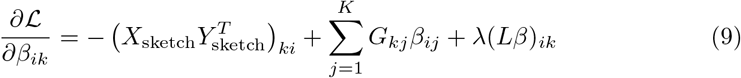

The spatial regularization term expands as (*Lβ*)_*ik*_ = *d*_*i*_*β*_*ik*_ − ∑_*n*∈*N*(*i*)_ *β*_*nk*_, where *d*_*i*_ = |*N*(*i*)| is the degree of spot *i*. Setting the gradient to zero and solving for *β*_*ik*_ yields the closed-form update:

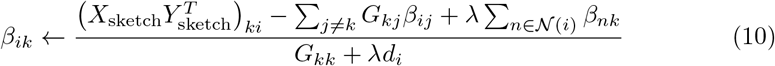

To incorporate *ℓ*_1_ regularization and non-negativity, we apply the proximal operator for the non-negative *ℓ*_1_ penalty: 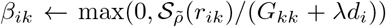, where 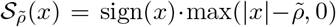 is the soft-thresholding operator with the scale-invariant threshold 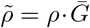 (Section 2). Algorithm 2 details the complete procedure [69]. Importantly, our objective decomposes as *f*(*β*) + *g*(*β*), where *f* (data fidelity and Laplacian terms) is smooth and convex, and *g* (*ℓ*_1_ penalty and non-negativity constraint) is separable and convex. For this class of composite convex problems, block coordinate descent with proximal updates is guaranteed to converge to the global optimum [70]; empirical validation on real biological data is provided in Supplementary Fig. S3.

The key computational advantages of this algorithm are: (1) the Gram matrix *G* is precomputed once with size *K* × *K* (typically *K* ≈ 10–20), and the cross-product 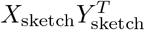 is cached to avoid redundant computation across iterations, (2) neighbor lookups are *O*(*k*) per spot where *k* ≪ *N*, and (3) the inner loops are JIT-compiled using Numba for near-C performance. The overall complexity per iteration is *O*(*N* · *K* · (*K* + *k*)), which is linear in the number of spots.

### Uncertainty Quantification

At convergence, the BCD update denominator for coordinate (*i, k*) equals 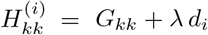, the diagonal of the Hessian of the smooth loss. A Laplace approximation gives the marginal variance of the unnormalized coefficient:

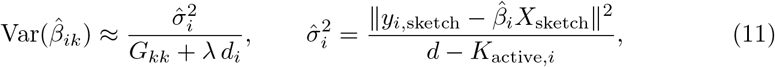

#### Algorithm 2

Block Coordinate Descent for Spatial Deconvolution

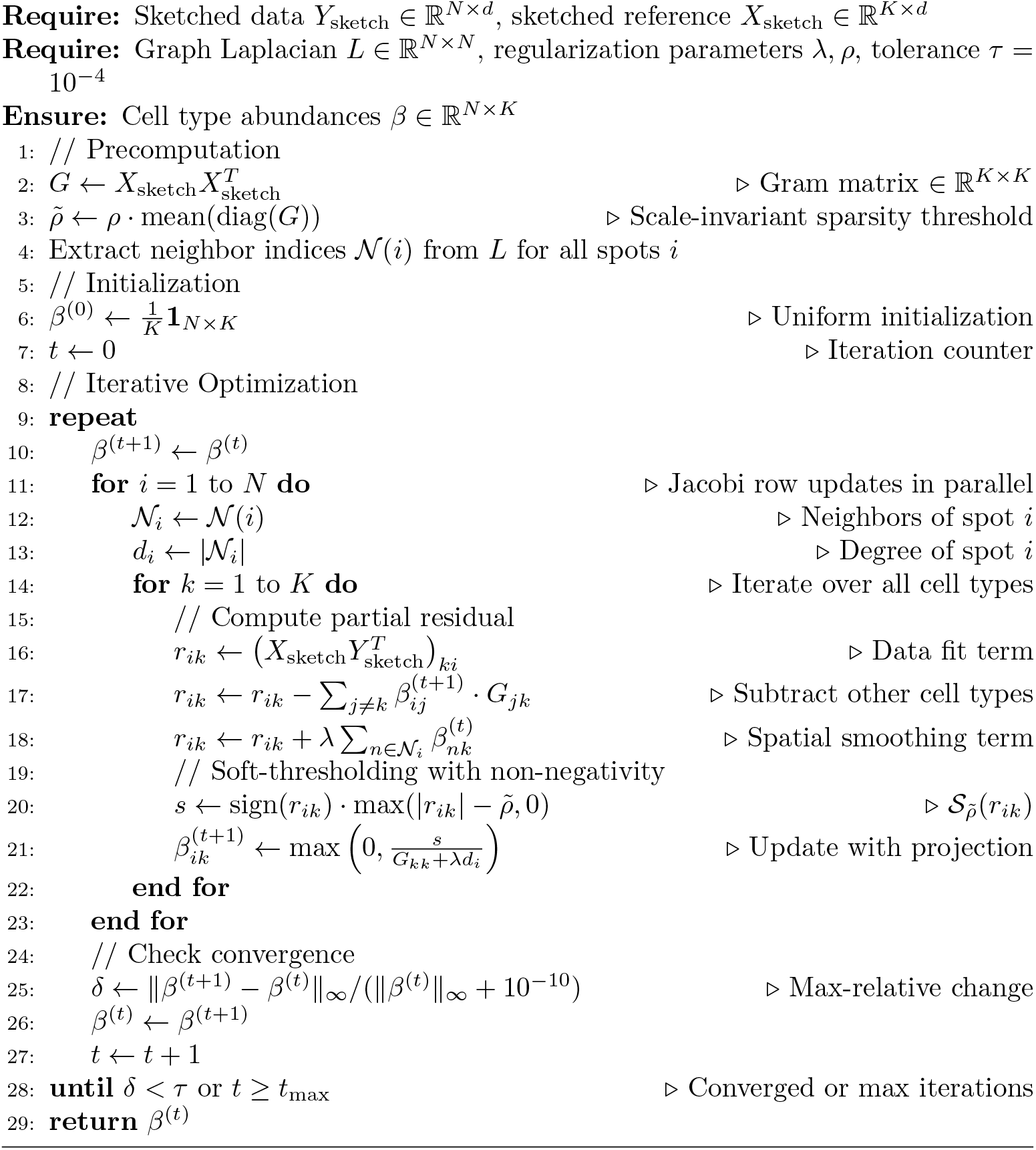

where *K*_active,*i*_ is the number of cell types with 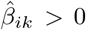 at spot *i*. Proportion-scale variance follows by the delta method: 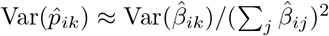. As an independent validation, we implement a Poisson parametric bootstrap: counts are resampled as 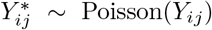, re-preprocessed, re-sketched with the same hash function, and solved via warm-start BCD (20 iterations). Percentile confidence intervals from *B* = 100 replicates provide model-free uncertainty bands (Supplementary Note).

### Benchmarking

We evaluated FlashDeconv on the Spotless benchmark suite [12], which provides standardized ground-truth data for deconvolution method comparison:

- Silver Standard: 54 synthetic datasets (6 tissues ×9 abundance patterns) generated by computationally mixing scRNA-seq profiles. Each dataset contains 1,000–5,000 pseudo-spots with known cell type proportions, enabling systematic evaluation across diverse biological contexts (brain cortex, cerebellum [cell and nucleus references], hippocampus, kidney, skin) and abundance scenarios (dominant types, rare types, uniform distribution).
- Gold Standard: Real spatial transcriptomics data with ground-truth proportions derived from co-registered imaging. STARMap (1 dataset, 108 spots, mouse visual cortex) provides single-molecule imaging-derived validation, while seqFISH+ (7 fields of view, <10 spots each, mouse cortex and olfactory bulb) tests performance on extremely small sample sizes.

To ensure strictly comparable evaluation, performance metrics for competing methods (Cell2Location, RCTD, Stereoscope, etc.) were obtained directly from the official Spotless benchmark results [12]. FlashDeconv was evaluated on identical source datasets using the same ground-truth labels and metric computation procedures.

Performance metrics include Root Mean Square Error (RMSE), Pearson correlation coefficient, and Area Under the Precision-Recall Curve (AUPR) for rare cell type detection. For Gold Standard data, we additionally report Jensen-Shannon Divergence (JSD) following the Spotless protocol. All runtime and memory benchmarks were performed on an Apple MacBook Pro with M2 Max chip (32GB unified memory) running macOS, representing consumer-grade hardware without GPU acceleration.

## Supporting information

Supplementary Data 1: Benchmark Results

## Data Availability

The Spotless benchmark datasets are available at Zenodo (https://zenodo.org/records/10277187) with code at https://github.com/saeyslab/spotless-benchmark. The Mouse Brain Visium dataset was obtained from the cell2location data portal (https://cell2location.cog.sanger.ac.uk/tutorial/); raw data are available at ArrayExpress (accession E-MTAB-11114 for Visium data, E-MTAB-11115 for snRNA-seq reference) [5]. The Visium HD Mouse Small Intestine (FFPE) dataset was obtained from 10x Genomics (https://www.10xgenomics.com/datasets/visium-hd-cytassist-gene-expression-libraries-of-mouse-intestine). The Xenium Fresh Frozen Mouse Colon dataset, used for ground truth validation of deconvolution accuracy with single-cell resolution data, was obtained from 10x Genomics (https://www.10xgenomics.com/datasets/fresh-frozen-mouse-colon-with-xenium-multimodal-cell-segmentation-1-standard). The intestinal scRNA-seq reference was obtained from Haber et al. [49]; we used the pre-processed version from Zenodo (https://zenodo.org/records/4447233), which contains 10,896 cells with cell type annotations. The original data is deposited at GEO (GSE92332). The human ovarian cancer Visium dataset is available at GEO (https://www.ncbi.nlm.nih.gov/geo/query/acc.cgi?acc=GSE211956) [45]. The Visium HD colorectal cancer cohort data and associated Chromium Flex scRNA-seq reference are from Oliveira et al. [42] and available at Zenodo (https://doi.org/10.5281/zenodo.15042463) and 10x Genomics (https://www.10xgenomics.com/datasets/visium-hd-cytassist-gene-expression-libraries-of-human-crc).

## Code Availability

FlashDeconv is implemented as an open-source Python package available at https://github.com/cafferychen777/flashdeconv. Complete analysis scripts for reproducing all figures and benchmarks presented in this paper are available at https://github.com/cafferychen777/flashdeconv-reproducibility.

## Acknowledgements

We acknowledge the publicly available datasets used in our benchmarking, including those from the Spotless benchmark suite, Cell2location mouse brain data, and 10x Genomics Visium HD datasets. Portions of this research were conducted with the advanced computing resources provided by the Texas A&M Department of Statistics Arseven Computing Cluster. We also thank the developers of the open-source tools utilized in this work.

## Author Contributions

C.Y., X.Z., and J.C. jointly conceptualized the study. C.Y. developed the methodology, implemented the computational framework, conducted formal analysis, and performed data curation, visualization, and validation. X.Z. and J.C. jointly supervised the research, secured funding, provided resources, served as corresponding authors, and administered the project, contributing equally in these senior roles. C.Y., X.Z., and J.C. all participated in manuscript writing and revision.

