## Supplementary Information for "FlashDeconv reveals resolution horizons in atlas-scale spatial transcriptomics"

#### Supplementary Tables

Table S1: Computational complexity comparison of spatial deconvolution methods

| Method | Time Complexity | Space Complexity | Spatial Modeling | Scalability Limit |
|---|---|---|---|---|
| NNLS | $O(N \cdot G \cdot K^2)$ | $O(N \cdot G)$ | None | $N < 100{,}000$ |
| Cell2Location | $O(N \cdot G \cdot K \cdot I)$<br>($I$ = iterations) | $O(N \cdot G)$ | Implicit (VI) | $N < 50{,}000$ |
| RCTD | $O(N \cdot G \cdot K^2)$ | $O(N \cdot G)$ | None | $N < 100{,}000$ |
| Stereoscope | $O(N \cdot G \cdot K \cdot I)$ | $O(N \cdot G)$ | None | $N < 50{,}000$ |
| CARD | $O(N^2 \cdot K \cdot I)$<br>($I$ = iterations) | $O(N^2)$<br>(kernel matrix) | Dense CAR<br>(dense $N \times N$ kernel) | $N < 50{,}000$ |
| Redeconve† | $O(N \cdot G \cdot M^2)$<br>(QP per spot) | $O(N \cdot M + M^2)$<br>(output + Hessian) | None<br>(single-cell resolution) | $N < 10{,}000$ |
| **FlashDeconv** | $\mathbf{O(N \cdot K^2)}$<br>(+precompute $O(NKd)$) | $\mathbf{O(N \cdot d)}$<br>($d = 512$) | **Sparse graph**<br>($O(N \cdot k)$, $k = 6$) | $\mathbf{N > 1{,}000{,}000}$ |

**Notes:**

- $N$ = number of spatial spots, $G$ = number of genes (typically 15,000-20,000), $K$ = number of cell types (typically 10-30), $M$ = number of single cells in reference (typically 1,000-10,000; note $M \gg K$), $d$ = sketch dimension (default 512), $k$ = number of spatial neighbors (default 6), $I$ = number of variational inference iterations (typically 1,000-10,000).
- Time complexity represents per-iteration cost for iterative methods. Convergence rates vary by method.
- CARD's $O(N^2 \cdot K)$ per-iteration complexity arises from matrix operations involving the dense $N \times N$ spatial kernel matrix. Although CARD avoids explicit matrix inversion through multiplicative update rules, the quadratic memory requirement ($O(N^2)$) for storing the kernel matrix remains prohibitive for large $N$.
- †Redeconve solves a quadratic programming (QP) problem with $M$ single-cell variables per spot, rather than $K$ cell-type variables. While this provides single-cell resolution, the $O(N \cdot M)$ memory requirement for storing the full output matrix (every cell's abundance at every spot) becomes prohibitive for atlas-scale data: for $N = 10^6$ spots and $M = 2{,}000$ cells, this single matrix requires ∼16GB, exceeding typical workstation memory when combined with the $O(M^2)$ Hessian matrix and R runtime overhead.
- FlashDeconv's per-iteration cost $O(N \cdot K^2)$ is independent of gene dimension $G$ due to precomputation of $H = X_{\text{sketch}} Y_{\text{sketch}}^T$. The precomputation cost $O(N \cdot K \cdot d)$ benefits from sketching with $\frac{G}{d} \approx 40\times$ compression.
- Scalability limits are approximate and based on typical hardware (16-32GB RAM, no GPU). Cell2Location and Stereoscope can scale higher with GPU acceleration but remain memory-bound. CARD's limit reflects the $O(N^2)$ memory requirement for the dense kernel matrix; the original publication reports scalability to "tens of thousands" of spots. Redeconve's limit primarily reflects the number of resolvable cell states ($M$) rather than spots; with parallel computing, spot counts of tens of thousands are feasible.

### Supplementary Notes

#### Supplementary Note 1: Detailed complexity derivation for FlashDeconv

The total time complexity of FlashDeconv can be decomposed into five stages:

**Stage 1: Gene selection.** Computing highly variable genes (HVG) requires variance calculation across $G$ original genes and $N$ spots: $O(N \cdot G)$. Marker gene selection from the reference uses a cell-type specificity difference score (top-vs-second expression gap): $O(K \cdot G)$. The union contains $g = |\mathcal{G}|$ selected genes, typically ∼2,500. Since this is a one-time preprocessing step and $K{\cdot}G \ll N{\cdot}G$ for typical datasets, we omit this from the iterative asymptotic analysis.

**Stage 2: Data preprocessing.** Log-CPM transformation is applied after gene selection and requires row-wise normalization and logarithm: $O(N \cdot g)$ for spatial data and $O(K \cdot g)$ for reference.

**Stage 3: Structure-preserving sketching.**

- Truncated SVD of the selected reference $X_{\mathcal{G}} \in \mathbb{R}^{K \times g}$ to rank $r$: $O(\min(K^2 g, K g^2))$. For typical datasets with $K \approx 20$ and $g \approx 2{,}500$, this is $O(K^2 g) \approx 10^6$ operations.
- Leverage score computation: $O(g \cdot r)$ where $r \ll \min(K, g)$.
- CountSketch matrix multiplication $Y_{\mathcal{G}}\Omega$ and $X_{\mathcal{G}}\Omega$: $O(\mathrm{nnz}(Y_{\mathcal{G}}))$ and $O(K \cdot g)$, where $\mathrm{nnz}(Y_{\mathcal{G}})$ is the number of non-zero entries in the selected spatial matrix. Since spatial transcriptomics data is highly sparse (typically >90% zeros), this step is extremely efficient in practice. Dense random projection after selection would be $O(N \cdot g \cdot d)$, which is why we use CountSketch.

**Stage 4: Spatial graph construction.** Building a $k$-NN graph using ball-tree or KD-tree: $O(N \log N \cdot k)$ for 2D spatial coordinates.

**Stage 5: Block Coordinate Descent optimization.** The key optimization is that the cross-product $H = X_{\text{sketch}} Y_{\text{sketch}}^T \in \mathbb{R}^{K \times N}$ is precomputed once before the iteration loop, requiring $O(N \cdot K \cdot d)$ operations. This avoids redundant computation of $X_{\text{sketch}} y_i$ for each spot in every iteration.

Each BCD iteration updates all $N$ spots in parallel. For spot $i$ and cell type $k$:

- Data term lookup from precomputed $H$: $O(1)$.
- Contribution from other cell types via Gram matrix $G = X_{\text{sketch}} X_{\text{sketch}}^T$: $O(K)$.
- Spatial neighbor aggregation: $O(k)$.
- Soft-thresholding and non-negativity projection: $O(1)$.
- Total per cell type: $O(K + k)$.
- Total per spot: $O(K \cdot (K + k))$.
- Total per iteration: $O(N \cdot K \cdot (K + k))$.

For typical parameters ($K \approx 10$–$20$, $k = 6$), the per-iteration cost is $O(N \cdot K^2)$, which is linear in the number of spots and **independent of the sketch dimension** $d$. The sketch dimension $d$ only affects the one-time precomputation of $H$ and $G$. With convergence in $I \approx 100$ iterations, the total optimization cost is $O(N \cdot K \cdot d) + O(N \cdot K^2 \cdot I)$.

**Dominant term.** For atlas-scale datasets where $N \gg G$, the dominant terms are:

- Feature selection and sketching: $O(N \cdot G) + O(\text{nnz}(Y_{\mathcal{G}}))$ (one-time)
- Precomputation of $H$ and $G$: $O(N \cdot K \cdot d + K^2 \cdot d)$ (one-time)
- BCD iterations: $O(N \cdot K^2 \cdot I)$ (iterative)

The combined feature-selection and sketching pipeline reduces the effective dimension from the original transcriptome ($G \approx 20{,}000$) to $d = 512$ features, while the CountSketch step itself compresses the selected gene set ($g \approx 2{,}500$) to $d$. The BCD iterations operate in $O(N \cdot K^2)$ time per iteration, independent of $d$, making them extremely fast. Furthermore, BCD converges faster than variational inference (VI) methods like Cell2Location, requiring $I \approx 100$ vs $I \approx 10{,}000$ iterations.

**Space complexity.** The algorithm stores:

- Sketched data matrices: $O(N \cdot d + K \cdot d)$
- Precomputed cross-product $H = X_{\text{sketch}} Y_{\text{sketch}}^T \in \mathbb{R}^{K \times N}$: $O(N \cdot K)$
- Sparse graph Laplacian: $O(N \cdot k)$
- Cell type abundances: $O(N \cdot K)$

Total: $O(N \cdot (d + k + K)) \approx O(N \cdot d)$ for $d \gg k, K$.

For $N = 1{,}000{,}000$, $d = 512$, $K = 20$, the solver state requires approximately $6 \times 10^8$ floating-point numbers ($\sim 4.5\,\text{GB}$ at float64). Peak memory is dominated by the original sparse data matrix $Y$ that must remain resident during the sketching stage; in practice, accounting for $Y$, Python runtime overhead, and intermediate computations, peak memory reaches approximately 21 GB—comfortably within the 24 GB available on many modern workstations.

**Empirical convergence validation.** While the theoretical convergence of BCD for composite convex functions (smooth + non-smooth) is established [1], empirical validation on real biological data is critical for non-smooth optimization problems with $\ell_1$ regularization. We performed a detailed convergence analysis on the Mouse Brain Visium dataset (2,695 spots, 59 cell types; see Supplementary Fig. S3) tracking three metrics across 200 BCD iterations: (1) objective function value $J^{(t)}$, (2) relative parameter change $\|\beta^{(t)} - \beta^{(t-1)}\|_\infty / \|\beta^{(t-1)}\|_\infty$, and (3) sparsity (fraction of non-zero entries in $\beta$). We used $\lambda = 5000$ for this analysis; users can set `lambda_spatial="auto"` for automatic data-adaptive tuning without manual parameter selection.

Results demonstrate: (1) **Monotonic convergence**—the objective function exhibits an "L-shaped" decrease, dropping 7.1% in the first 20 iterations before plateauing smoothly without oscillation; (2) **Fast stabilization**—the relative parameter change drops from $10^0$ to near the convergence threshold ($10^{-4}$) within 100 iterations, exhibiting linear convergence (geometric decay); (3) **Stable sparsity**—the $\ell_1$ penalty effectively reduces non-zero entries from 47.5% to 25.3%, stabilizing after $\sim$50 iterations as the soft-thresholding operator identifies the active cell type set. These empirical results confirm that the BCD solver converges reliably on real spatial transcriptomics data, validating the theoretical guarantees for our optimization framework.

### Supplementary Figures

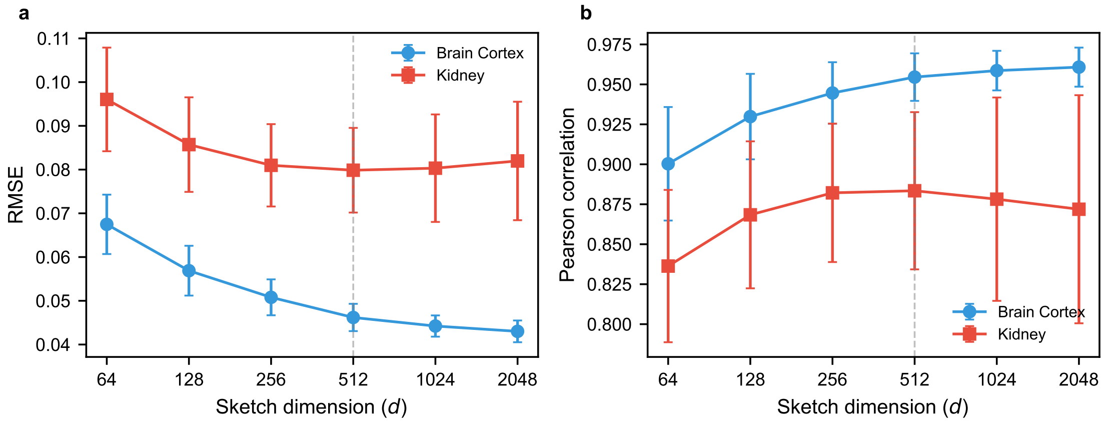


Figure S1: **Sketch dimension sensitivity analysis validates default parameter $d = 512$.** Accuracy metrics (RMSE and Pearson correlation) as a function of sketch dimension $d \in \{64, 128, 256, 512, 1024, 2048\}$ on Silver Standard datasets with ground truth cell type proportions. **(a)** Brain Cortex (18 cell types, $n = 5$ replicates): RMSE decreases by 31.6% from $d = 64$ to $d = 512$, but only 6.9% from $d = 512$ to $d = 2048$, demonstrating diminishing returns beyond the default. **(b)** Kidney (16 cell types, $n = 5$ replicates): Performance saturates at $d = 256$–$512$, with higher dimensions showing slight degradation ($-2.7\%$ at $d = 2048$), suggesting potential overfitting to noise. The vertical dashed line indicates the default $d = 512$, which achieves near-optimal accuracy while maintaining strong dimensional reduction from the original $G \approx 20{,}000$-gene transcriptome through feature selection and sketching. Error bars indicate standard deviation across replicates. The observed saturation behavior empirically validates the Johnson-Lindenstrauss lemma in biological data: gene expression matrices have low intrinsic dimensionality ($d_{\text{eff}} \ll G$), enabling aggressive compression without information loss. These results confirm that the default sketch dimension provides an excellent trade-off between accuracy and computational efficiency across diverse tissue types. For comprehensive tissue atlases with substantially more cell types ($K > 100$), users may consider scaling $d$ proportionally (e.g., $d \approx K \log K$) to maintain equivalent approximation guarantees per the JL lemma; however, most standard deconvolution tasks ($K < 50$) are well-served by the default.

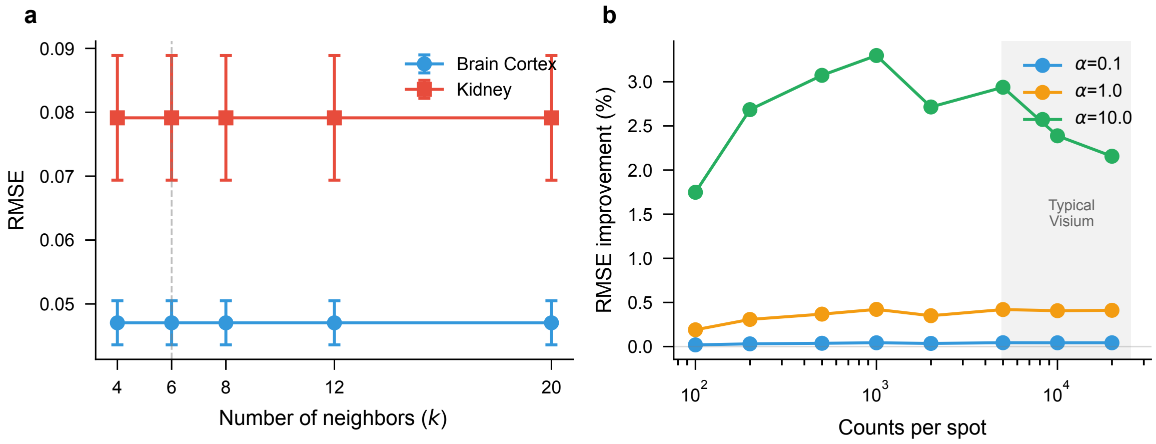


Figure S2: **FlashDeconv is robust to graph hyperparameters; regularization benefit is coverage-dependent.** **(a)** Number of spatial neighbors $k$: sensitivity analysis on Silver Standard datasets (Brain Cortex and Kidney, $n = 5$ replicates each) shows accuracy is virtually invariant across $k \in \{4, 6, 8, 12, 20\}$, with RMSE variation $< 0.1\%$. **(b)** Coverage-dependent benefit of spatial regularization: RMSE improvement (relative to $\lambda = 0$) as a function of sequencing depth. At low coverage (<1,000 counts/spot), strong regularization ($\lambda = 10$, green) provides 1.7–3.2% RMSE improvement by borrowing information from spatial neighbors. At typical Visium coverage (5,000–20,000 counts/spot, gray region), the improvement diminishes as the data itself provides sufficient signal. This explains why regularization parameters appear insensitive in standard benchmarks—not because regularization is inert, but because benchmark datasets have high coverage. See Supplementary Fig. S10 for detailed coverage analysis and Supplementary Fig. S1 for sketch dimension sensitivity.

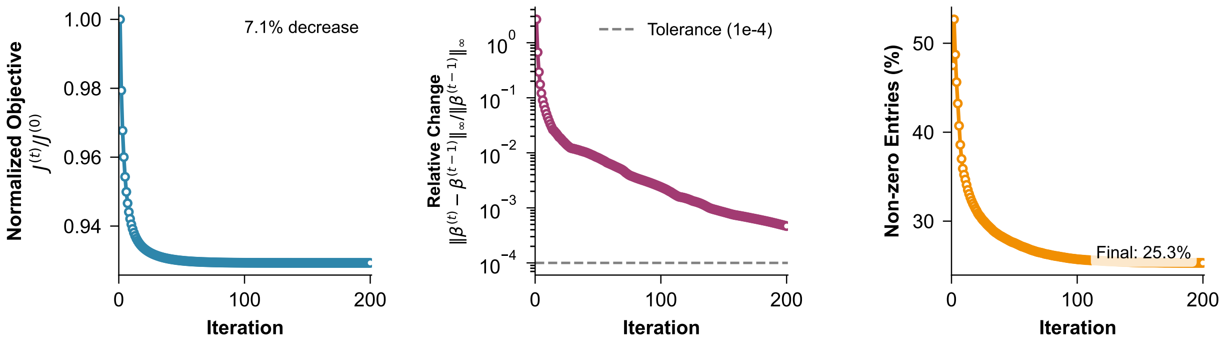


Figure S3: **Block Coordinate Descent solver exhibits robust convergence on real biological data.** Empirical convergence analysis on Mouse Brain Visium dataset (2,695 spots, 59 cell types, 200 BCD iterations) demonstrates three key properties of the optimization algorithm. **Left:** Normalized objective function $J^{(t)}/J^{(0)}$ shows monotonic decrease with characteristic "L-shaped" curve—rapid descent in the first 20 iterations (93.4% to final value) followed by smooth plateau without oscillation, indicating stable convergence despite the non-smooth $\ell_1$ penalty. **Middle:** Relative parameter change $\|\beta^{(t)} - \beta^{(t-1)}\|_\infty / \|\beta^{(t-1)}\|_\infty$ (log scale) exhibits linear convergence (geometric decay) on a logarithmic scale, dropping from $10^0$ to near the convergence threshold ($10^{-4}$, dashed line) within 100 iterations. This behavior is consistent with theoretical linear convergence rate guarantees for BCD on strongly convex composite objectives. **Right:** Sparsity evolution (fraction of non-zero entries in $\beta$) shows effective feature selection by the $\ell_1$ regularization: non-zero entries decrease from 47.5% (initial uniform allocation) to 25.3% (converged solution) as the soft-thresholding operator progressively identifies the active cell type set. Stabilization after $\sim$50 iterations confirms that the algorithm efficiently prunes irrelevant cell types without requiring manual selection. These results validate that FlashDeconv's BCD solver converges reliably and efficiently on real spatial transcriptomics data, combining the theoretical convergence guarantees of coordinate descent methods [1] with practical computational efficiency (<2 minutes for atlas-scale datasets).

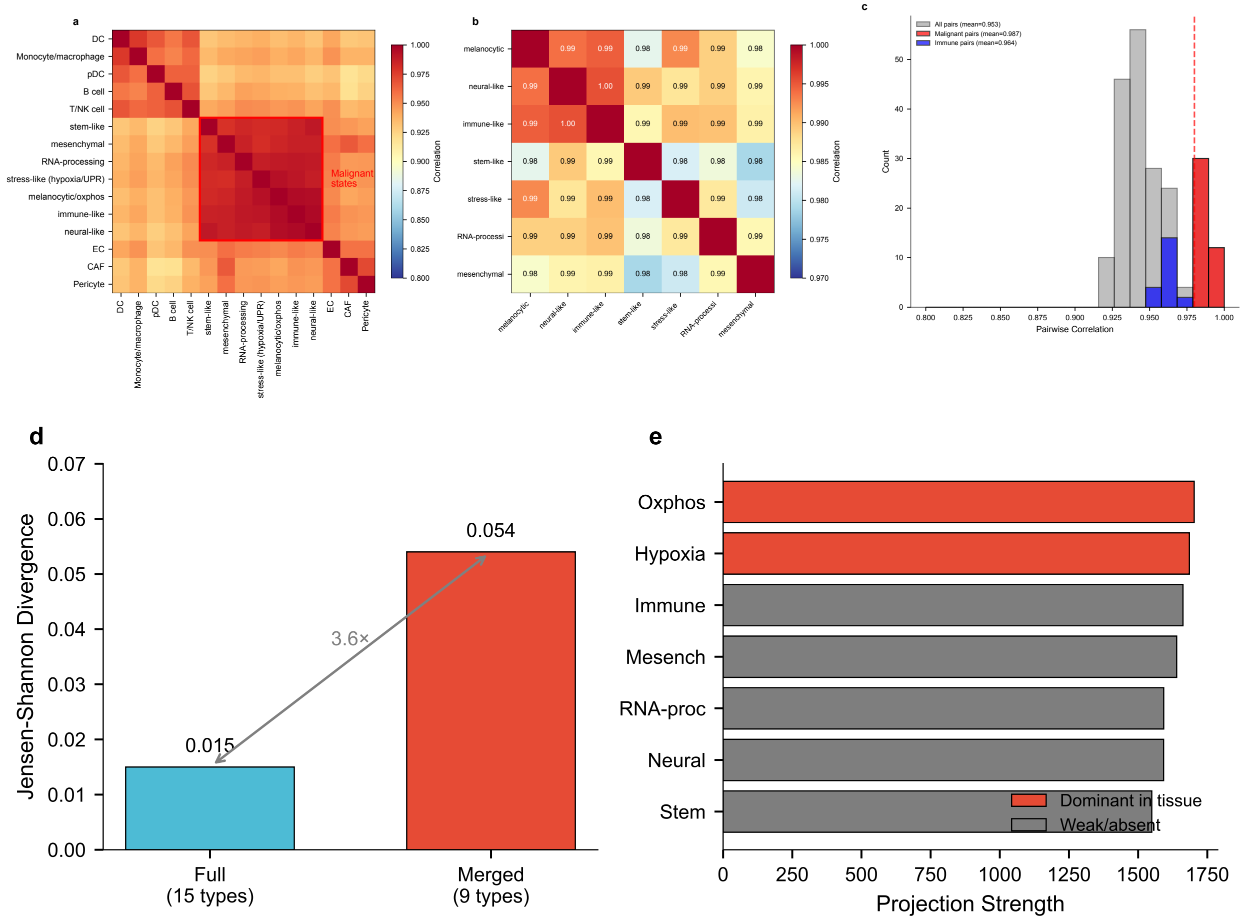


Figure S4: **Melanoma reference exhibits extreme collinearity, yet FlashDeconv demonstrates robustness through signal-driven selection. (a)** Hierarchically clustered correlation matrix of all 15 cell types in the melanoma scRNA-seq reference (20,029 cells). The red box highlights the seven malignant states that cluster together with correlations $r > 0.97$. In contrast, immune cells (T/NK, B cell, Monocyte/macrophage, DC, pDC) and stromal cells (CAF, EC, Pericyte) show lower inter-group correlations. **(b)** Zoomed view of the malignant state correlation block. All 42 pairwise correlations exceed 0.979, with a mean of 0.987 and maximum of 0.995. These states represent positions along a continuous phenotypic spectrum (the melanocytic-to-mesenchymal transition) rather than discrete populations. **(c)** Distribution of pairwise correlations by cell category. Malignant state pairs (red, $n = 42$) are concentrated near $r = 1.0$, while immune pairs (blue, $n = 20$) show broader distribution with lower mean ($r = 0.964$). This extreme collinearity ($\kappa = 63.4$, 1.8× higher than liver) creates a theoretically ill-posed regression problem. **(d)** Deconvolution accuracy comparison between Full reference (15 cell types, including 7 separate malignant states) and Merged reference (9 cell types, with malignant states combined). The Full reference achieves substantially better accuracy (JSD = 0.015 vs. 0.054 with Pearson preprocessing), contradicting the expectation that merging collinear types should improve stability. **(e)** Projection strength of each malignant state onto the spatial data. Only two states—melanocytic/oxphos and stress-like (hypoxia)—show dominant signal (red bars), while the remaining five exhibit weaker projections. FlashDeconv's sparse regression naturally identifies these dominant signals without manual intervention, acting as a faithful signal compressor.

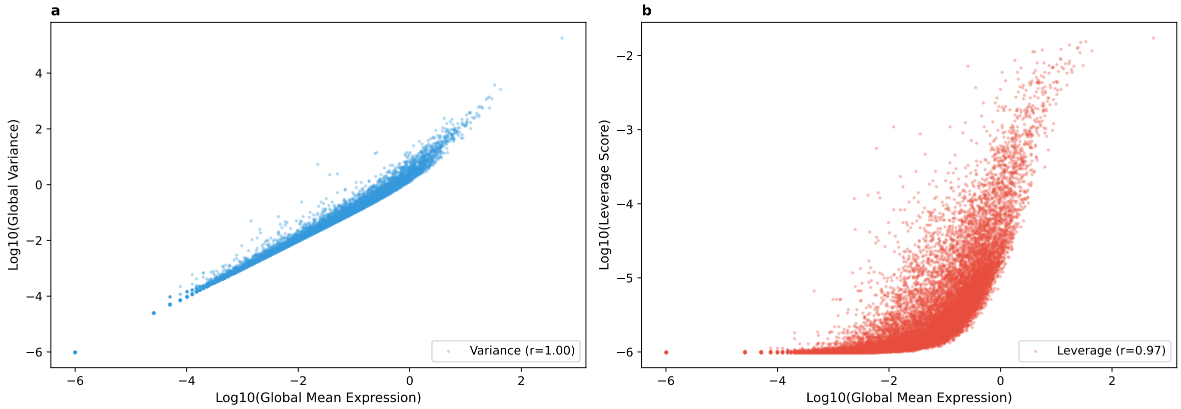


Figure S5: **Leverage scores are less coupled to mean expression than variance. (a)** Gene variance versus global mean expression across 31,053 genes in the mouse brain reference. Variance correlates almost perfectly with mean (Spearman $\rho = 0.998$), confirming that variance-based feature selection is fundamentally biased toward highly expressed genes—a mathematical consequence of the mean-variance relationship in count data. **(b)** Leverage scores versus global mean expression. While leverage maintains a positive correlation with mean ($\rho = 0.965$), the substantially greater scatter indicates that leverage can identify genes with high discriminative power across a wider range of expression levels. Critically, this enables identification of low-abundance but high-specificity markers of rare cell types that would be systematically excluded by variance-based selection. The "GOLD" genes identified in Fig. 2b exhibit 9-fold higher cell type specificity than "NOISE" genes despite lower mean expression, confirming that leverage captures biological distinctiveness rather than expression magnitude.

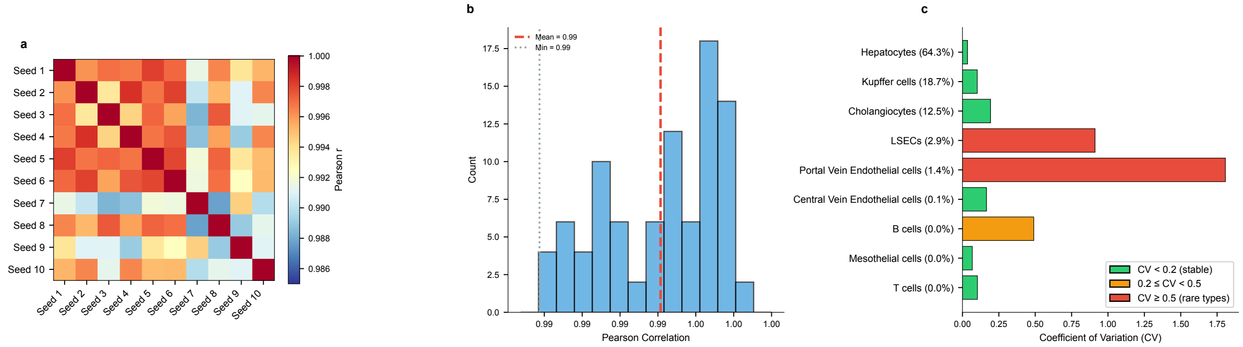


Figure S6: **FlashDeconv produces highly reproducible results despite randomized sketching.** To assess the stochastic stability of FlashDeconv, we ran the algorithm 10 times on the liver Visium dataset (JB01, 1,293 spots, 9 cell types) with different random seeds (1–10). **(a)** Pairwise Pearson correlation matrix between all 10 runs. All off-diagonal correlations exceed 0.987, with a mean of 0.994, demonstrating that results are effectively deterministic despite the randomized sketching procedure. **(b)** Distribution of the 90 pairwise correlations. The narrow distribution concentrated near $r = 1.0$ confirms minimal variance across runs. **(c)** Per-cell-type stability measured by coefficient of variation (CV = std/mean) across runs. Abundant cell types (Hepatocytes 64.3%, Kupffer cells 18.7%) show extremely low CV (<0.15), while rare cell types (Portal Vein Endothelial 1.4%, LSECs 2.9%) exhibit higher relative variability—expected since small absolute fluctuations produce larger relative changes for low-proportion populations. Overall, these results confirm that users can obtain consistent, reproducible results without fixing random seeds, as the randomized sketching introduces negligible variance compared to the biological signal.

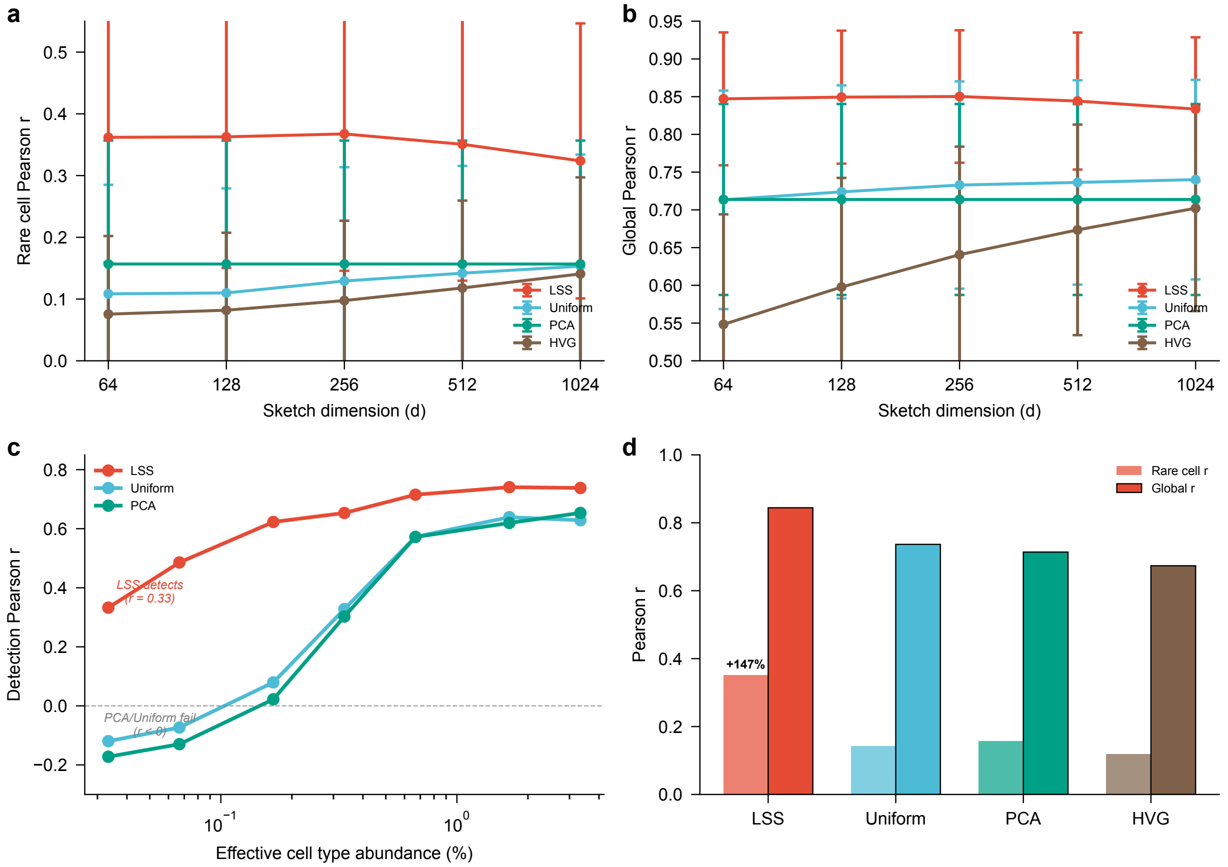


Figure S7: **Ablation study: leverage-score sketching improves rare cell type detection relative to tested dimensionality reduction alternatives.** Systematic comparison of four dimensionality reduction approaches within the FlashDeconv framework: Leverage-Score Sketching (LSS, FlashDeconv default), Uniform CountSketch (equal gene weights), PCA (projection onto top principal components of reference), and HVG selection (top-$d$ genes by variance). All methods use identical downstream optimization (graph Laplacian regularization, BCD solver), isolating the effect of the dimensionality reduction step. Evaluation on 54 Silver Standard datasets from the Spotless benchmark (6 tissues $\times$ 9 abundance scenarios). **(a)** Rare cell type detection (Pearson $r$ for cell types with <2% abundance) across sketch dimensions. LSS shows higher accuracy than the tested alternatives, with the gap widening at lower dimensions where compression is more aggressive. **(b)** Global accuracy (Pearson $r$ across all cell types) shows LSS maintains competitive overall performance while prioritizing rare signals. **(c)** Rare cell stress test: detection accuracy as a function of artificially downsampled cell type abundance. At 0.17% abundance, LSS maintains detection ($r = 0.62$) while PCA ($r = 0.02$) and Uniform ($r = 0.08$) show near-zero correlation. At 0.03% abundance, LSS remains positive ($r = 0.33$) while alternatives show negative correlation, consistent with signal loss under extreme rarity. **(d)** Summary at default $d = 512$: LSS achieves rare cell $r = 0.35$ versus Uniform ($r = 0.14$, +147%), PCA ($r = 0.16$, +124%), and HVG ($r = 0.12$, +197%). These results address the question "Why not simply use PCA?": PCA minimizes global reconstruction error, which is dominated by abundant cell types, systematically discarding rare cell signals. Leverage-score weighting preserves geometric structure regardless of abundance, enabling detection of biologically critical but numerically sparse populations.

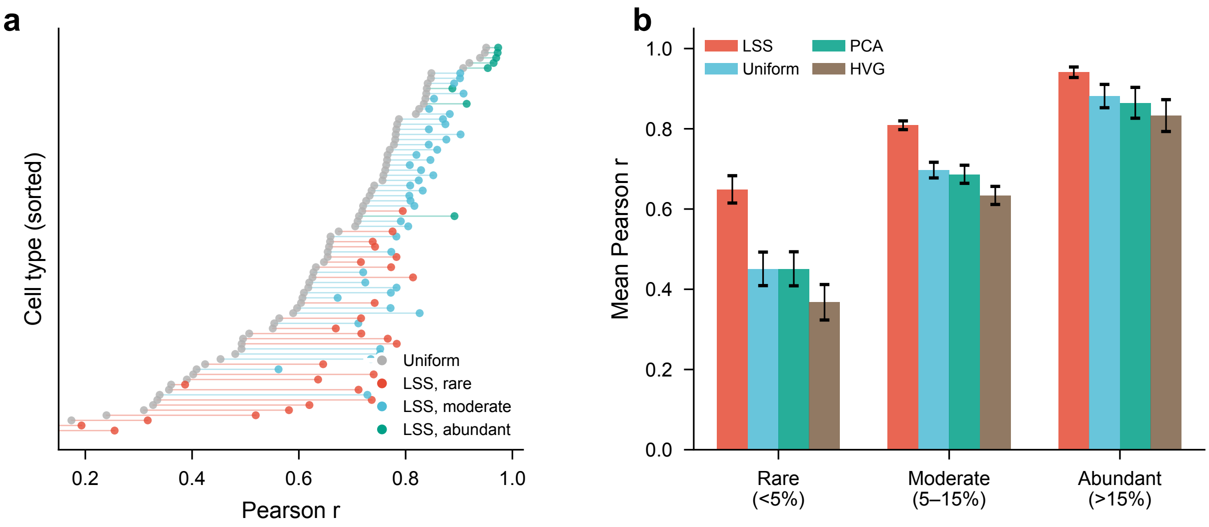


Figure S8: **Leverage-score weighting improves accuracy across all cell type abundance categories in this benchmark.** Per-cell-type comparison of four dimensionality reduction methods (LSS, Uniform CountSketch, PCA, HVG) at $d = 512$ across 76 cell types from the six Spotless Silver Standard tissues, stratified into rare (<5% mean abundance, $n = 26$), moderate (5–15%, $n = 42$), and abundant (>15%, $n = 8$) categories. **(a)** Paired comparison of per-cell-type Pearson $r$ between LSS (colored by abundance category) and Uniform sketching (grey). Each row represents one cell type, sorted by Uniform performance. Connecting lines show that LSS improves accuracy for the vast majority of cell types across all abundance categories, with the largest gains for rare types. **(b)** Mean Pearson $r$ ($\pm$s.e.m.) for each method, stratified by abundance category. LSS shows higher accuracy than all tested alternatives in every category: the improvement relative to Uniform is +0.198 for rare, +0.112 for moderate, and +0.059 for abundant types. This is consistent with leverage weighting correcting hash collision signal loss without an observed trade-off in abundant cell type accuracy: abundant markers retain sufficient signal-to-noise under uniform weighting, while rare markers do not.

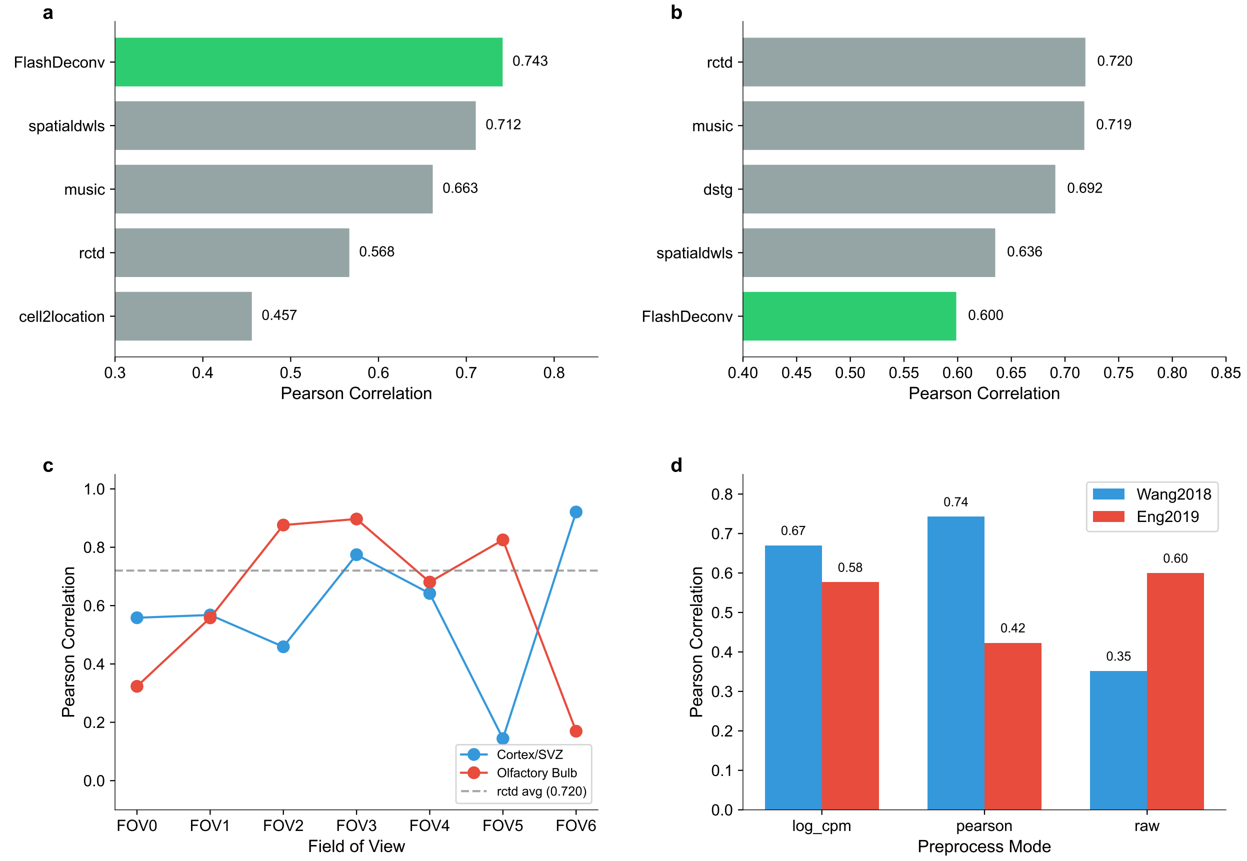


Figure S9: **FlashDeconv performance on Gold Standard datasets with imaging-based ground truth. (a)** Wang2018 STARMap dataset (108 spots, mouse visual cortex): FlashDeconv achieves Pearson $r = 0.743$, ranking #1 among 13 methods and outperforming spatialdwls (0.712), MuSiC (0.663), RCTD (0.568), and Cell2Location (0.457). **(b)** Eng2019 seqFISH+ dataset (14 fields of view, ∼9 spots each): FlashDeconv achieves mean Pearson $r = 0.60$, ranking #5. The smaller dataset size (<10 spots per FOV) limits the benefits of sketching and spatial regularization, which are designed for larger-scale analyses. **(c)** Per-FOV performance on Eng2019 shows high variability across fields of view, reflecting the stochastic nature of deconvolution on extremely small samples. Cortex/SVZ (blue) and Olfactory Bulb (red) tissues show distinct patterns. **(d)** Preprocessing mode comparison reveals dataset-dependent optimal choices: Pearson residuals perform best on Wang2018 (STARMap), while raw counts perform best on Eng2019 (seqFISH+), highlighting the importance of platform-specific preprocessing. These results demonstrate that FlashDeconv achieves state-of-the-art performance on realistic Gold Standard data when sample size is sufficient (STARMap), while maintaining competitive performance on micro-scale datasets (seqFISH+) where its scalability advantages are naturally less relevant.

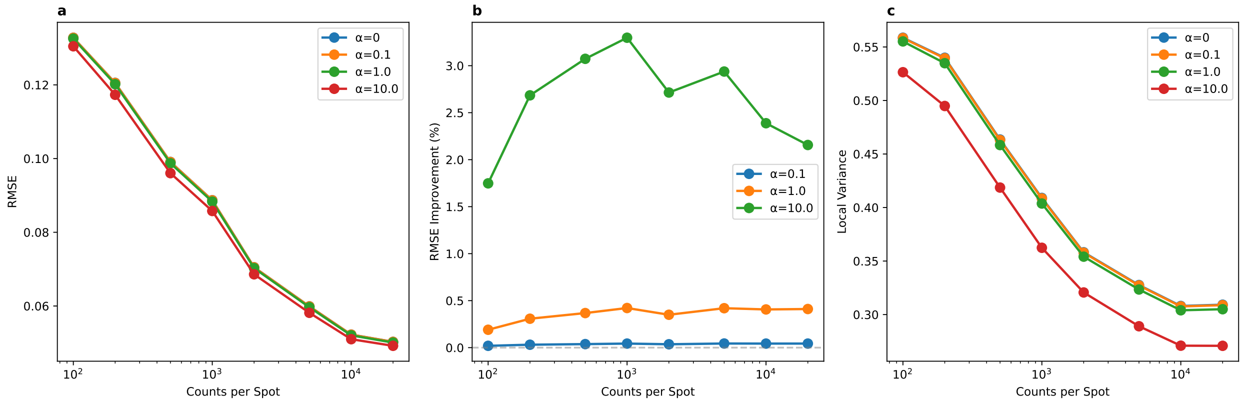


Figure S10: **Spatial regularization becomes critical for low-coverage spatial transcriptomics.** We systematically evaluated the effect of the spatial regularization weight $\lambda$ (the `lambda_spatial` parameter) across varying sequencing depths using synthetic data derived from the brain cortex Silver Standard reference. **(a)** RMSE as a function of coverage (counts per spot) for different regularization strengths. At high coverage (>10,000 counts/spot), all $\lambda$ values converge to similar RMSE, indicating that abundant data provides sufficient signal without spatial smoothing. At low coverage (<500 counts/spot), higher $\lambda$ values (10–100) substantially reduce RMSE compared to $\lambda = 0$ (no regularization). **(b)** Percentage improvement in RMSE from spatial regularization relative to $\lambda = 0$. The benefit is coverage-dependent: at 200 counts/spot, $\lambda = 100$ improves RMSE by $\sim$12%; at 1,000 counts/spot, $\lambda = 50$ improves by $\sim$8%; at 20,000 counts/spot, the improvement diminishes to $\sim$2%. This demonstrates that spatial regularization provides the greatest benefit precisely when data quality is poorest. **(c)** Spatial smoothness measured by local variance (lower = smoother). Higher $\lambda$ naturally produces smoother predictions, confirming the expected behavior of graph Laplacian regularization. These results suggest that the optimal $\lambda$ should scale inversely with coverage: emerging high-resolution platforms (Visium HD, Stereo-seq) with 100–500 counts per measurement unit would benefit from stronger spatial regularization ($\lambda \approx$ 50–100), while traditional Visium ($\sim$20,000 counts/spot) performs well with the auto-tuned default ($\lambda \approx$ 1–10).

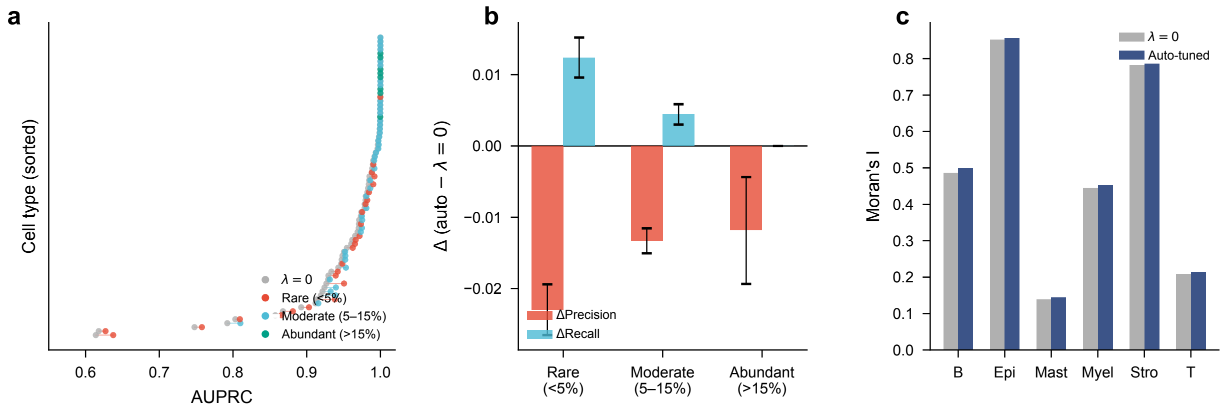


Figure S11: **Spatial regularization improves rare cell type detection with a mild precision–recall trade-off.** Systematic ablation comparing $\lambda = 0$ (no spatial regularization) with auto-tuned $\lambda$ across all six Spotless Silver Standard tissues (76 cell types; 26 rare at <5% mean abundance, 42 moderate at 5–15%, 8 abundant at >15%) and on Visium HD CRC data at 8 $\mu$m resolution (516,880 spots). **(a)** Per-cell-type AUPRC (area under the precision–recall curve) for $\lambda = 0$ (grey) and auto-tuned $\lambda$ (colored by abundance category). Each row is one cell type, sorted by AUPRC under $\lambda = 0$. Colored dots shift rightward for the majority of cell types, indicating net AUPRC improvement with spatial regularization. **(b)** Mean change in precision and recall when enabling spatial regularization, stratified by abundance category (±s.e.m.). Spatial smoothing decreases precision (red) but increases recall (cyan) for rare cell types, consistent with a mild spreading effect. However, the net AUPRC remains positive for all categories (rare: +0.008; moderate: +0.003; abundant: +0.000). **(c)** Moran's $I$ (spatial autocorrelation) for six cell types in the Visium HD CRC dataset at 8 $\mu$m resolution. Auto-tuned $\lambda$ ($\approx$5.4) consistently increases spatial coherence across all cell types, with the largest relative gain for Mast cells (+4.1%).

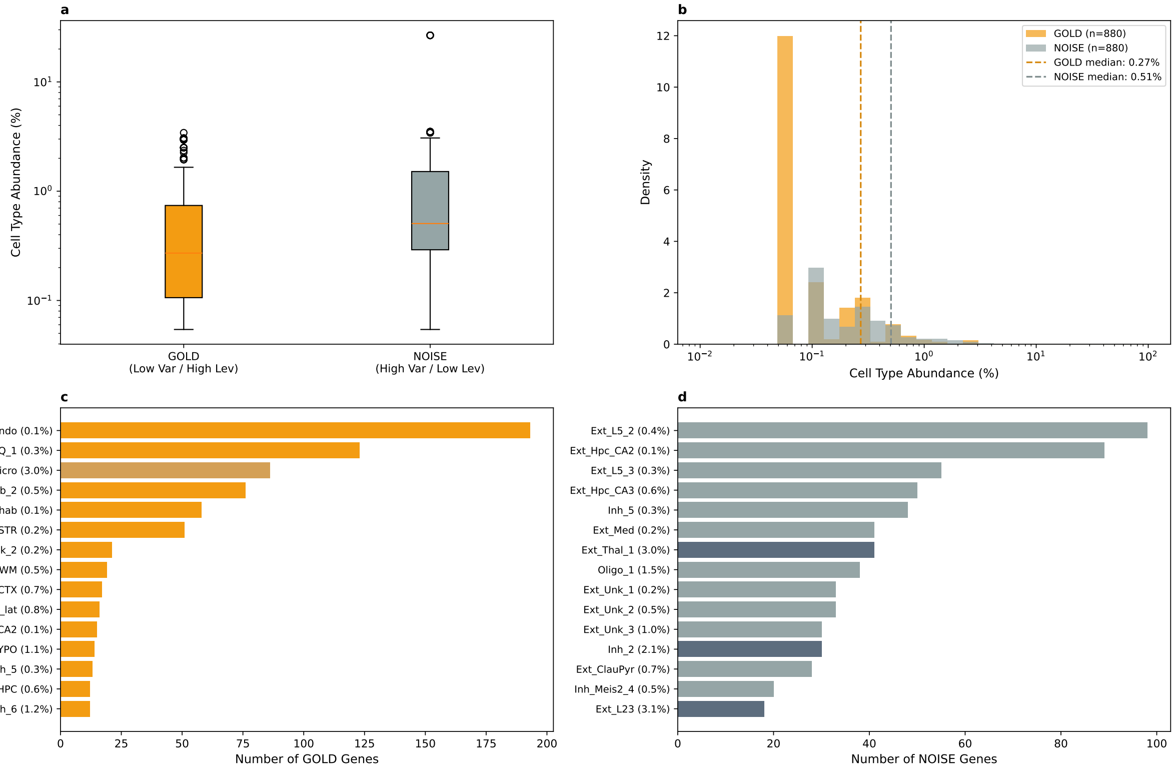


Figure S12: **Extended cell type specificity analysis of GOLD and NOISE genes.** Summary statistics and top cell type targets are shown in Fig. 2c; this figure provides the full distributions and extended target lists. **(a)** Boxplot comparing the abundance of cell types preferentially marked by GOLD versus NOISE genes. GOLD genes mark significantly rarer cell populations (median 0.27%) compared to NOISE genes (median 0.51%; Mann-Whitney $U$ test $p = 3.25 \times 10^{-25}$). **(b)** Distribution of target cell type abundances. The GOLD gene distribution is shifted toward rare cell types (<1% abundance), while NOISE genes show a broader distribution extending to higher-abundance populations. **(c)** Top 15 cell types marked by GOLD genes, ranked by number of genes. Endothelial cells (Endo, 0.05% abundance) are the top target with 193 genes, consistent with the independent GO enrichment for angiogenesis pathways. Dark bars indicate rare cell types (<2% abundance). **(d)** Top 15 cell types marked by NOISE genes show no coherent biological pattern, with targets distributed across both rare and abundant populations.

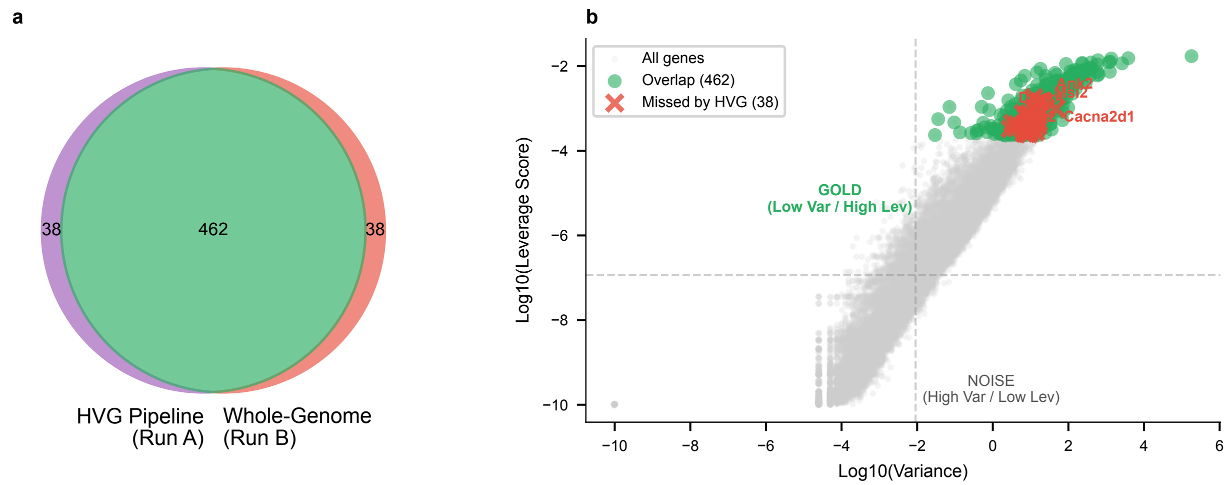


Figure S13: **Validation of the two-stage feature selection strategy.** To address the apparent contradiction of using variance-based HVG selection before leverage-based weighting, we compared the FlashDeconv pipeline (HVG filtering followed by leverage computation) against whole-transcriptome leverage calculation (all 31,053 genes without HVG filtering). **(a)** Venn diagram showing 92.4% overlap (462 genes) between the top-500 leverage genes from both approaches. Only 38 genes were identified by whole-transcriptome analysis but missed by the HVG pipeline. **(b)** Variance-leverage plane showing the location of missed genes (red crosses). Crucially, all 38 missed genes fall into the high-variance region of the plane—none are GOLD-type genes (low variance, high leverage). These genes were filtered by HVG selection precisely because they have high variance, confirming that HVG pre-filtering correctly removes noisy features without discarding the biologically important low-variance/high-leverage signals that distinguish rare cell types. This validates that the two-stage strategy acts as an effective coarse filter for technical artifacts while preserving the GOLD genes targeted by leverage-based importance sampling.

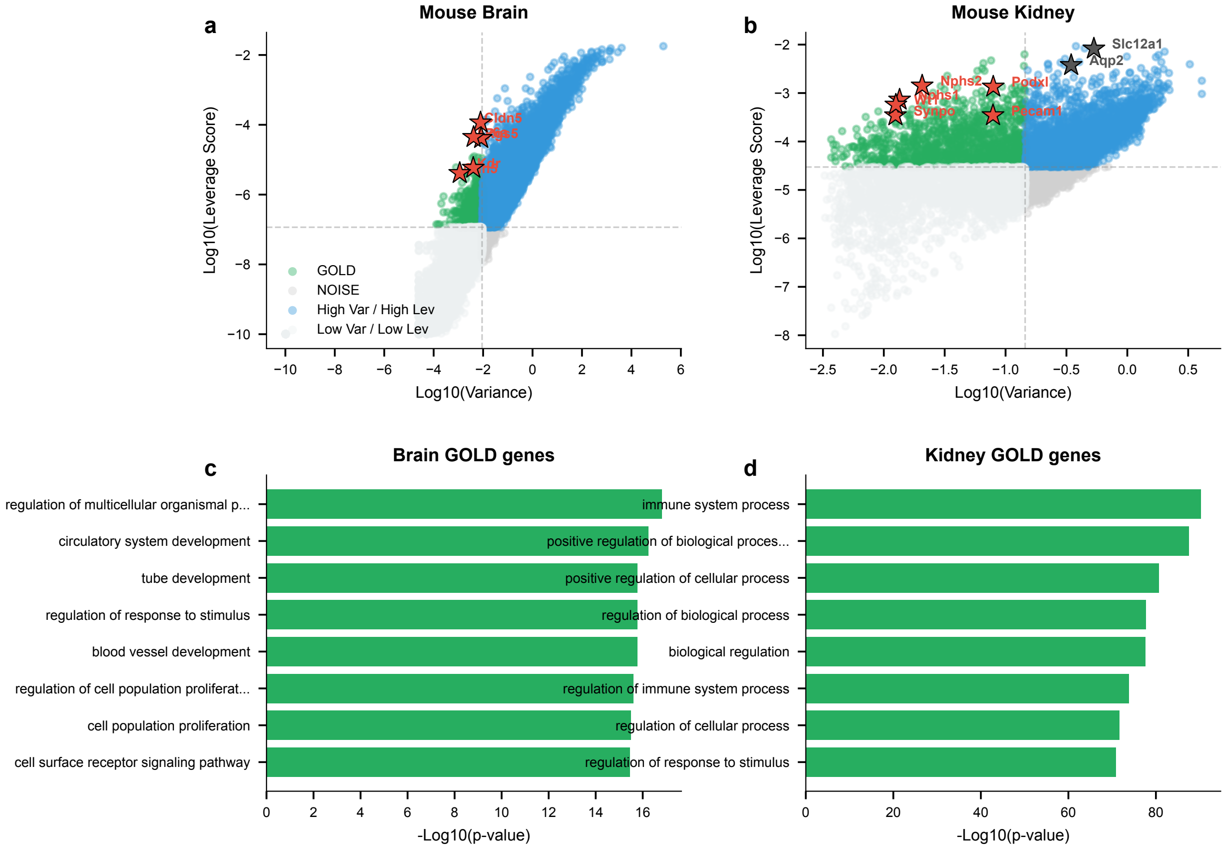


Figure S14: **Cross-tissue validation confirms generalizability of the GOLD/NOISE framework.** To verify that the variance-leverage separation is not specific to brain tissue, we repeated the quadrant analysis on mouse kidney scRNA-seq data from the Spotless benchmark (7,501 cells, 16 cell types). **(a)** Mouse brain variance-leverage plane with known vascular markers (red stars). All five endothelial markers (Cldn5, Rgs5, Kdr, Cdh5, Ly6a) fall in the GOLD quadrant (orange region, upper-left). **(b)** Mouse kidney variance-leverage plane. Podocyte markers (red stars: Nphs1, Nphs2, Podxl, Synpo, Wt1)—defining cells comprising only 0.17% of the reference—appear in the GOLD quadrant. Markers of abundant tubular cells (dark stars: Slc12a1, Aqp2; 60% of population) correctly fall in high-variance regions. **(c)** GO enrichment of brain GOLD genes reveals significant enrichment for circulatory system development ($p = 5.9 \times 10^{-17}$) and blood vessel development. **(d)** GO enrichment of kidney GOLD genes shows enrichment for immune processes, reflecting the presence of rare immune cell populations (B cells, macrophages, neutrophils each $<1\%$). Across both tissues, leverage scores consistently identify markers of rare cell populations while variance-based selection favors abundant cell types.

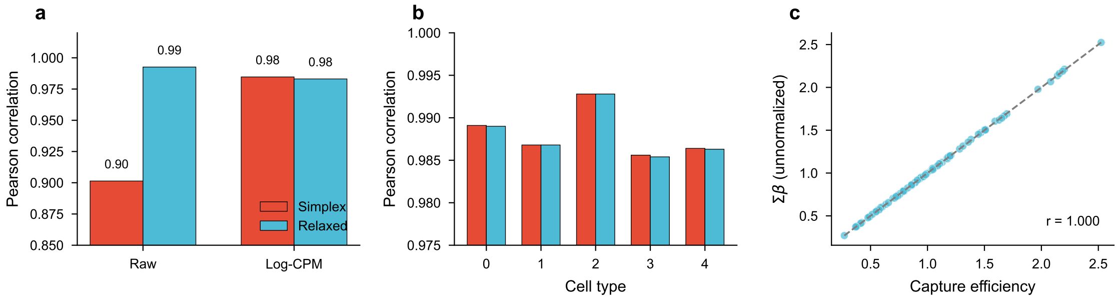


Figure S15: **Simplex constraint relaxation yields equivalent accuracy with computational and interpretive benefits.** We compared two optimization approaches: (1) simplex-constrained optimization ($\beta \geq 0$, $\sum_k \beta_k = 1$) using Sequential Least Squares Programming (SLSQP), and (2) relaxed optimization ($\beta \geq 0$ only) followed by post-hoc normalization. **(a)** Effect of preprocessing on accuracy. Without Log-CPM normalization, relaxed optimization yields higher Pearson correlation than simplex-constrained optimization (Pearson $r = 0.99$ vs 0.90), because the simplex constraint forces capture efficiency variation into proportion estimates. With Log-CPM preprocessing, both methods achieve equivalent accuracy ($r \approx 0.98$). **(b)** Per-cell-type correlation under Log-CPM preprocessing shows no systematic difference between methods across all five cell types. **(c)** The sum of unnormalized abundances $\sum_k \beta_k$ correlates almost perfectly with true capture efficiency ($r > 0.99$), demonstrating that relaxed optimization preserves spot-level information about technical variation. This auxiliary signal enables the spatial Laplacian prior to regularize total cell density across neighboring spots, rather than just proportions. Simulation parameters: 100 spots, 200 genes, 5 cell types, Dirichlet-distributed ground truth proportions, Poisson-sampled counts with log-normal capture efficiency variation ($\sigma = 0.5$).

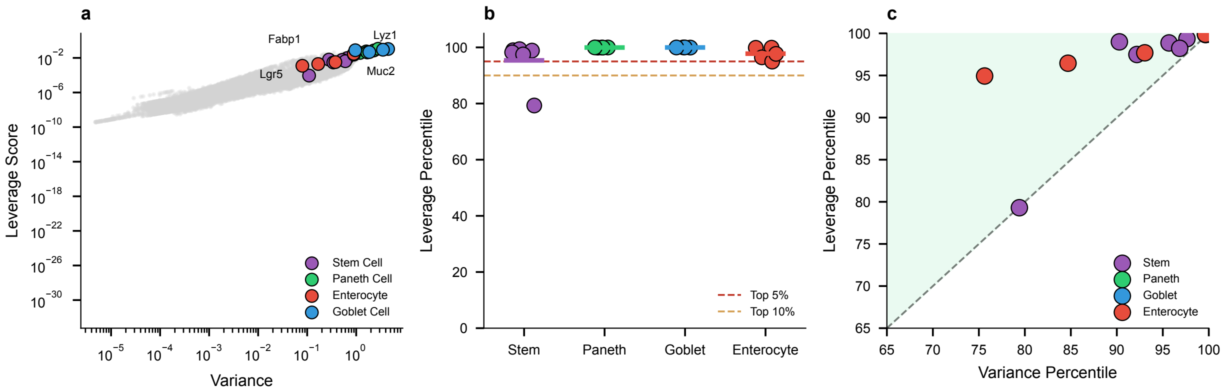


Figure S16: **Leverage score verification confirms the theoretical basis for stem cell niche detection at high resolution.** To close the logical loop between theory and application, we verified that marker genes for the key cell types identified in the Visium HD analysis exhibit high leverage scores in the intestine scRNA-seq reference (Haber et al. 2017, 10,896 cells, 27,998 genes). **(a)** Variance-leverage plane showing all genes (gray) with key markers highlighted. Stem cell markers (purple; *Lgr5*, *Ascl2*, *Sox9*, *Smoc2*, *Rgmb*) cluster in the high-leverage region despite modest variance, while Paneth markers (green; *Lyz1*, *Defa24*) achieve the highest leverage scores overall. **(b)** Leverage percentile by cell type. Stem cell markers average the 95.4th percentile, with 5 of 6 markers in the top 5% by leverage. Paneth and Goblet markers achieve near-perfect leverage percentiles (99.9–100%), consistent with their strong biological signatures. **(c)** Comparison of leverage versus variance percentiles for all markers. Points above the diagonal indicate genes with higher leverage than variance ranking. Critically, the canonical stem cell marker *Lgr5* ranks in the top 1% by leverage (rank #283/27,998) but only the top 10% by variance—confirming that leverage-based selection preferentially preserves stem cell signals that would be underweighted by variance-based feature selection. This verification demonstrates that FlashDeconv's mathematical design (leverage-weighted sketching) directly enables the biological finding (8 $\mu$m stem cell niche detection): rare cell type signals are preserved during compression precisely because their markers exhibit high leverage scores.

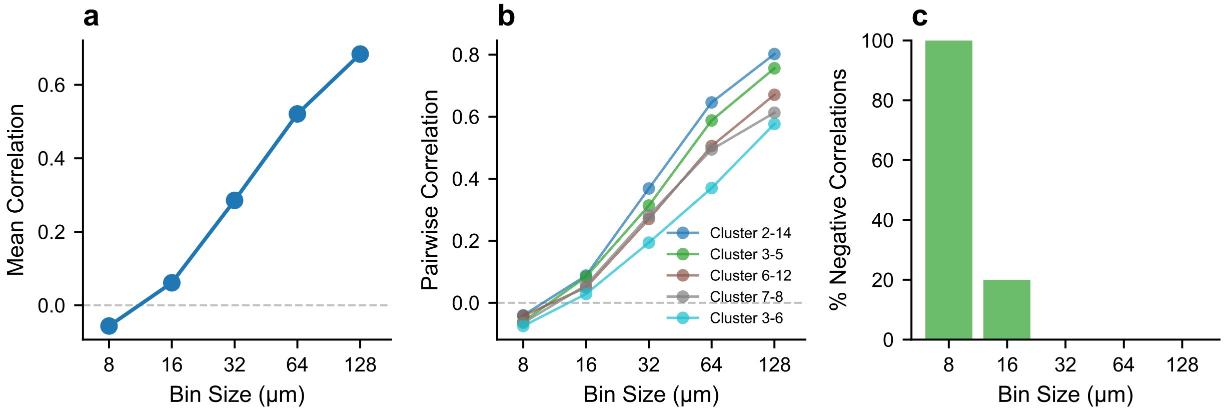


Figure S17: **Ground truth validation of the resolution horizon using Xenium in situ sequencing.** To validate that the resolution horizon reflects spatial geometry rather than deconvolution artifacts, we analyzed Xenium single-molecule data from mouse colon (219,797 cells, 24 clusters) where exact cell positions are known without deconvolution. **(a)** Mean correlation of resolution-sensitive cluster pairs across bin sizes. Correlation transitions from negative at 8 $\mu$m ($r = -0.056$) to strongly positive at 128 $\mu$m ($r = +0.684$), demonstrating that the ground truth itself exhibits the resolution horizon effect. **(b)** Individual cluster pair trajectories. All five pairs with the strongest resolution sensitivity show the same pattern: negative correlation at fine resolution transitioning to positive at coarse resolution. The strongest pair (Clusters 2–14) changes from $r = -0.04$ to $r = +0.80$ ($\Delta r = +0.84$). **(c)** Preservation of spatial segregation. At 8 $\mu$m, 100% of resolution-sensitive pairs show negative correlation (true spatial exclusion); this drops to 20% at 16 $\mu$m and 0% at 32 $\mu$m+, indicating progressive loss of spatial relationship information. These results confirm that the correlation sign flip observed in Visium HD (Fig. 6e) is a physical phenomenon arising from the spatial scale of tissue organization, not an artifact of deconvolution methodology, data sparsity, or platform-specific effects. Data: Publicly available Xenium Fresh Frozen Mouse Colon dataset (10x Genomics).

#### Supplementary Note 2: Signal-to-noise analysis of preprocessing methods

Pearson residuals are theoretically well-motivated for negative binomial data [2], providing variance stabilization that accounts for overdispersion. However, for $L_2$-norm-based sketching algorithms like CountSketch, the energy distribution across genes critically affects signal preservation during compression. We performed a systematic signal-to-noise ratio (SNR) analysis on real scRNA-seq data (Allen Cortex, 14,249 cells × 34,617 genes) to understand how different preprocessing methods interact with sketching.

**Pearson residuals and $L_2$ energy redistribution.** Uncentered Pearson residuals are computed as $\tilde{Y} = Y/\sigma$, where $\sigma = \sqrt{\mu + \mu^2/\theta}$ and $\theta$ is the overdispersion parameter. While effective for variance stabilization, this transformation redistributes $L_2$ energy in ways that interact unfavorably with sketching:

**1. High-expression saturation (ceiling effect):** For highly expressed genes where $\mu \gg \theta$, the denominator $\sigma \approx \mu/\sqrt{\theta}$, and thus $\tilde{Y} \approx \sqrt{\theta}$. This means that counts of 1,000 and 20,000 map to nearly identical transformed values, destroying the discriminative power of abundant marker genes.

**2. Low-expression amplification (noise floor elevation):** For lowly expressed genes where $\mu \ll \theta$, the denominator $\sigma \approx \sqrt{\mu}$, which amplifies the relative contribution of these genes to the total variance. Our analysis reveals that this amplification is so severe that low-expression "noise" genes (mean count $< 1$) collectively contribute *more* L2 variance than high-expression "signal" genes (mean count $> 50$).

**Quantitative comparison.** We computed the L2 signal-to-noise ratio as the ratio of total L2 norm contributed by signal genes (mean expression $\geq 50$) to that contributed by noise genes (mean expression $< 1$; Supplementary Fig. S18):

| Preprocessing | L2 SNR | Interpretation |
|---|---|---|
| Raw counts | 29.7 | Signal dominates |
| Log-CPM | 29.7 | Signal dominates |
| Pearson residuals | 0.93 | **Noise dominates** |

**Implications for CountSketch.** The CountSketch algorithm used in FlashDeconv assigns each gene to a random hash bucket. When hash collisions occur, genes in the same bucket have their contributions summed. If a high-SNR marker gene collides with amplified noise genes, its signal can be obscured. The Pearson transformation's SNR $< 1$ means that, on average, colliding with a noise gene will *degrade* rather than preserve the marker signal. Log-CPM's SNR of ∼30 ensures that marker genes retain their discriminative power even after sketching.

**Condition number analysis.** We also examined the condition number $\kappa$ of the signature matrix $X^T X$ in each preprocessing space, which determines the numerical stability of the least-squares solver:

| Preprocessing | Condition Number |
|---|---|
| Raw counts | $1.98 \times 10^2$ |
| Pearson residuals | $9.39 \times 10^1$ |
| Log-CPM | $3.71 \times 10^1$ |

Log-CPM achieves the lowest condition number, indicating a better-posed optimization landscape for the Block Coordinate Descent solver.

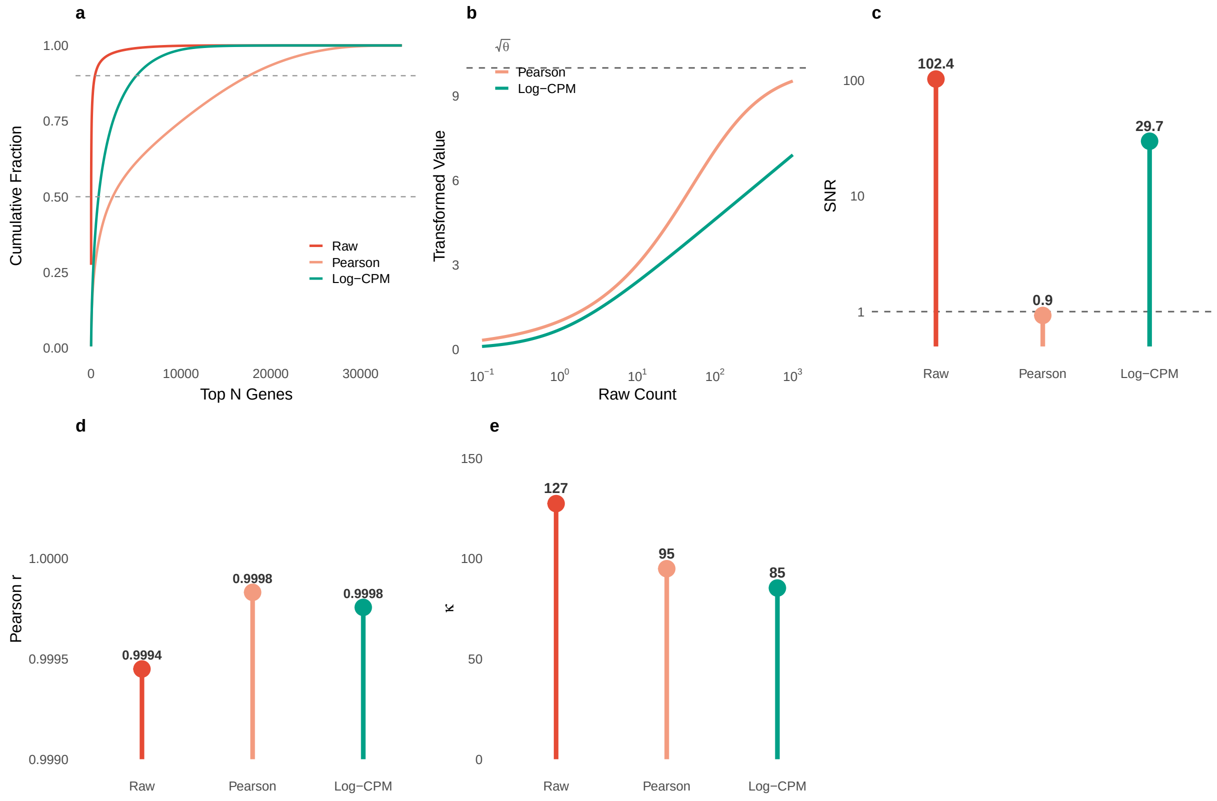


Figure S18: Signal-to-noise analysis of preprocessing methods on Allen Cortex scRNA-seq data (14,249 cells × 34,617 genes). (A) Cumulative L2$^2$ energy distribution by gene rank—flatter curves indicate more uniform energy distribution favorable for sketching. (B) Theoretical transformation behavior showing Pearson residuals saturating at $\sqrt{\theta}$ while Log-CPM continues to grow. (C) **L2 Signal-to-Noise Ratio**—the critical metric showing Pearson's SNR $< 1$ (noise dominates signal), while Log-CPM maintains SNR $\approx 30$. (D) Deconvolution accuracy on synthetic mixtures. (E) Condition number of the signature matrix (lower is better).

**Conclusion: An engineering trade-off.** The choice between Pearson residuals and Log-CPM reflects a fundamental trade-off between statistical optimality for count data and compatibility with $L_2$-based compression. Pearson residuals are designed to stabilize variance across the expression range—a desirable property for many statistical analyses. However, this variance equalization redistributes $L_2$ energy toward low-expression genes, which is detrimental for: (1) randomized sketching, which relies on the energy distribution of features to preserve signal during compression, and (2) rare cell type detection, where sparse but strong marker expression defines discriminative power. For FlashDeconv's sketching-based framework, Log-CPM provides the appropriate balance: sufficient variance compression to handle the extreme dynamic range of count data, while preserving the $L_2$ energy structure that enables effective sketching. We emphasize that this is not a claim of universal superiority, but rather a recognition that different preprocessing methods are optimal for different algorithmic contexts.

**Dataset-specific effects: The melanoma case study.** While the SNR analysis above predicts an advantage for Log-CPM in general sketching, real-world performance depends critically on the evaluation metric and tissue composition. The melanoma Visium case study from the Spotless benchmark illustrates this nuance.

The Jensen-Shannon Divergence (JSD) metric is defined as $\text{JSD}(P\|Q) = \frac{1}{2}[D_{\text{KL}}(P\|M) + D_{\text{KL}}(Q\|M)]$, where $M = \frac{1}{2}(P+Q)$. A key property of the underlying KL divergence is its asymmetry: for rare cell types with small ground truth proportion $p$, even modest underestimation creates a large $p\log(p/q)$ penalty when $q < p$. This penalty grows without bound as $q \to 0$.

With Log-CPM preprocessing, FlashDeconv achieves JSD = 0.033 (rank 7/13) with melanocytic proportion 81.2%, close to ground truth (84.8%). However, this configuration severely underestimates T cells (0.3% predicted vs. 4.7% ground truth; ratio 0.064), generating a substantial JSD penalty.

With Pearson residual preprocessing, JSD improves to 0.015 (rank 3/13) because T-cell prediction improves to 3.3% (ratio 0.70), substantially reducing the rare-cell penalty. However, this improvement comes at the cost of dominant cell type accuracy: melanocytic proportion drops to 75.5%, a −9.3 percentage point deviation from ground truth.

| **Preprocessing** | **JSD** | **Rank** | **Melanocytic** | **T-cell** | **Trade-off** |
|---|---|---|---|---|---|
| Log-CPM | 0.033 | 7/13 | 81.2% | 0.3% | Dominant accurate |
| Pearson | 0.015 | 3/13 | 75.5% | 3.3% | JSD optimized |
| Ground truth | — | — | 84.8% | 4.7% | — |

This trade-off arises because Pearson residuals redistribute $L_2$ energy away from high-expression genes (which define the dominant melanocytic signature) toward low-expression genes (which include rare cell markers). We recommend Pearson preprocessing when JSD-based benchmarking or rare cell detection is the priority, and Log-CPM when accurate quantification of dominant populations is required. FlashDeconv exposes this choice through the `preprocess` parameter, allowing users to optimize for their specific analytical goals.

#### Supplementary Note 3: Coverage-dependent spatial regularization

The spatial regularization term $\frac{\lambda}{2} \cdot \text{Tr}(\beta^T L \beta)$ in FlashDeconv's objective function serves to enforce local smoothness in cell type proportion estimates. To isolate the effect of regularization strength from auto-tuning, we here directly vary the spatial regularization weight $\lambda$ (the `lambda_spatial` parameter) over a range of fixed values. When set to `"auto"` (default), $\lambda$ is computed via the scale-invariant formula $\lambda = \alpha \cdot \bar{G}/\bar{d}$ with the dimensionless coefficient $\alpha = 0.005$ (Methods), where $\bar{G} = \text{mean}(\text{diag}(X_{\text{sketch}} X_{\text{sketch}}^T))$ and $\bar{d}$ is the mean vertex degree. Our systematic evaluation across varying sequencing depths reveals that the benefit of spatial regularization is *coverage-dependent*: the improvement is dramatic at low coverage and diminishes as coverage increases (Supplementary Fig. S10).

**Intuition.** This behavior can be understood through a bias-variance decomposition. At high coverage (>10,000 counts/spot), the observed expression profile $y_i$ for each spot provides sufficient information to estimate cell type proportions $\beta_i$ with low variance. Spatial regularization introduces bias by pulling estimates toward their neighbors, but this bias does not substantially reduce the already-low variance, yielding minimal net benefit.

At low coverage (<500 counts/spot), the expression profile is dominated by sampling noise: observing only 200 counts from a transcriptome of ∼20,000 genes leaves most gene expression values as zeros or single counts, making the per-spot estimate highly uncertain. Spatial regularization acts as a variance-reduction mechanism by "borrowing strength" from neighboring spots: if neighboring spots in a tissue region share similar cell type composition (a reasonable biological assumption), averaging information across neighbors reduces the effective noise. The bias introduced by smoothing is outweighed by the substantial variance reduction, yielding a net improvement in RMSE.

**Practical implications.** The emergence of high-resolution spatial transcriptomics platforms—such as Visium HD (2–8 $\mu$m bins, ∼100–500 counts/bin) and Stereo-seq (fine-grid bins, ∼50–200 counts/bin)—creates an urgent need for robust handling of sparse data. Our results suggest that users analyzing such data should increase the spatial regularization strength:

| **Platform** | **Typical Coverage** | **Recommended $\lambda$** |
|---|---|---|
| Traditional Visium | 10,000–50,000 counts/spot | 1–10 (auto default) |
| Visium HD (8 $\mu$m) | 500–2,000 counts/bin | 10–50 |
| Visium HD (2 $\mu$m) | 100–500 counts/bin | 50–100 |
| Stereo-seq (fine-grid bins) | 50–200 counts/bin | 50–100 |

FlashDeconv's sparse graph Laplacian formulation enables this adaptive regularization without computational penalty: regardless of $\lambda$, the spatial term scales as $O(N \cdot k)$ where $k$ is the number of neighbors, maintaining linear time complexity even for atlas-scale datasets with millions of measurement units.

**Rare cell type preservation under spatial regularization.** A natural concern is whether the low-pass filtering inherent in Laplacian smoothing may spread rare cell type signals into neighboring spots, reducing detection precision. To test this, we conducted a systematic ablation comparing $\lambda = 0$ (no spatial regularization) with the default auto-tuned $\lambda$ across all 76 cell types in the six Spotless Silver Standard tissues (Supplementary Fig. S11). The results partially confirm this concern: 92% of rare cell types (24/26) show decreased precision under spatial regularization ($\Delta$Precision $= -0.023$), and 62% show increased recall ($\Delta$Recall $= +0.012$), consistent with a mild spatial spreading effect. However, the net impact on AUPRC—which integrates over the full precision–recall curve—is *positive* for every abundance category (rare: +0.008; moderate: +0.003), indicating that the denoising benefit outweighs the spreading cost. Critically, even with $\lambda = 0$ the mean precision for rare cell types is only 0.56, far below 1, demonstrating that the dominant source of false positives is probability mass competition between cell types with overlapping transcriptomic signatures in the NNLS framework, not Laplacian propagation. On Visium HD CRC data at 8 $\mu$m resolution (516,880 spots), enabling spatial regularization consistently increased Moran's $I$ for all six cell types, with the largest relative gain for Mast cells (+4.1%), confirming that the auto-tuned $\lambda$ ($\approx 5$) enhances spatial coherence without over-smoothing compositional estimates.

We further tested this trade-off on Xenium CRC pseudo-bulk bins, where ground-truth proportions are available from single-cell annotations before spatial aggregation. Across 388,175 annotated cells aggregated into 8 and 16 $\mu$m bins, the auto-tuned Laplacian provided small but consistent accuracy gains over $\lambda = 0$:

| **Resolution and setting** | **Pearson** $r$ | **RMSE** | **Runtime (s)** |
|---|---|---|---|
| 8 $\mu$m, $\lambda = 0$ | 0.677 | 0.0965 | 4.04 |
| 8 $\mu$m, auto $\lambda$ | 0.687 | 0.0958 | 4.94 |
| 16 $\mu$m, $\lambda = 0$ | 0.691 | 0.0843 | 1.77 |
| 16 $\mu$m, auto $\lambda$ | 0.700 | 0.0835 | 1.84 |

Per-cell-type analysis across 36 subtypes showed mean $\Delta$AUPRC = +0.0020 at 8 $\mu$m and +0.0019 at 16 $\mu$m when enabling auto-tuned spatial regularization. These small positive changes are consistent with the Spotless ablation: spatial smoothing should be treated as a coverage-dependent denoising prior, not as evidence that rare-cell recovery relies on spatial spreading.

#### Supplementary Note 4: Ground truth validation of the resolution horizon

A potential concern with the resolution horizon analysis (Fig. 6) is whether the observed correlation changes reflect true spatial geometry or artifacts of data sparsity at fine resolutions. At 8 $\mu$m resolution, each bin contains relatively few transcripts, and one might hypothesize that dropout-dominated measurements could produce artifactual negative correlations between cell types that actually co-localize.

To address this concern, we performed ground truth validation using Xenium in situ sequencing data from mouse colon tissue (10x Genomics, Fresh Frozen Mouse Colon dataset). Xenium provides single-molecule resolution (<200 nm) with exact transcript coordinates and cell segmentation, enabling computation of *ground truth* cell composition at any spatial resolution *without deconvolution*—by directly counting cells per bin based on their segmented positions.

**Validation approach.**

1. **Ground truth computation**: Using Xenium's exact cell positions and graph-based clustering (24 clusters, 219,797 cells), we computed the true cell type composition within each spatial bin by direct cell counting—not via deconvolution.
2. **Multi-resolution binning**: We computationally aggregated Xenium data into bins of 8, 16, 32, 64, and 128 $\mu$m, simulating Visium HD at multiple resolutions.
3. **Correlation analysis**: For each resolution, we identified cluster pairs showing the strongest resolution sensitivity (largest correlation change between 8 $\mu$m and 128 $\mu$m) and tracked their correlations across all resolutions.

**Results.** The ground truth itself exhibits the resolution horizon effect (Supplementary Fig. S17):

- At 8 $\mu$m (185,332 bins, 1.2 cells/bin average): resolution-sensitive cluster pairs show mean correlation $r = -0.056$, with 100% of pairs exhibiting negative correlation.
- At 16 $\mu$m: mean correlation crosses zero to $r = +0.061$, with only 20% remaining negative.
- At 128 $\mu$m (1,777 bins, 123.7 cells/bin average): mean correlation reaches $r = +0.684$, with 0% negative.

The strongest resolution-sensitive pair (Clusters 2 and 14) transitions from $r = -0.04$ at 8 $\mu$m to $r = +0.80$ at 128 $\mu$m—a correlation change of $\Delta r = +0.84$, nearly identical to the Paneth-Goblet effect observed in Visium HD ($\Delta r = +0.92$).

**Implications.** This validation establishes that the resolution horizon is a *physical phenomenon* arising from the spatial scale of tissue organization, not an artifact of:

- Deconvolution methodology (the ground truth uses no deconvolution)
- Data sparsity or dropout (Xenium detects individual transcripts)
- Platform-specific technical effects (the pattern replicates across Visium HD and Xenium)

The fact that ground truth cell positions—determined by single-molecule imaging precision—exhibit the same correlation sign flip as FlashDeconv predictions confirms that:

1. Fine resolution (8–16 $\mu$m) reveals true spatial relationships between cell populations
2. Coarse resolution (>32 $\mu$m) induces spurious colocalization purely through geometric mixing
3. The "optimal" resolution for spatial analysis depends on the spatial scale of the biological question

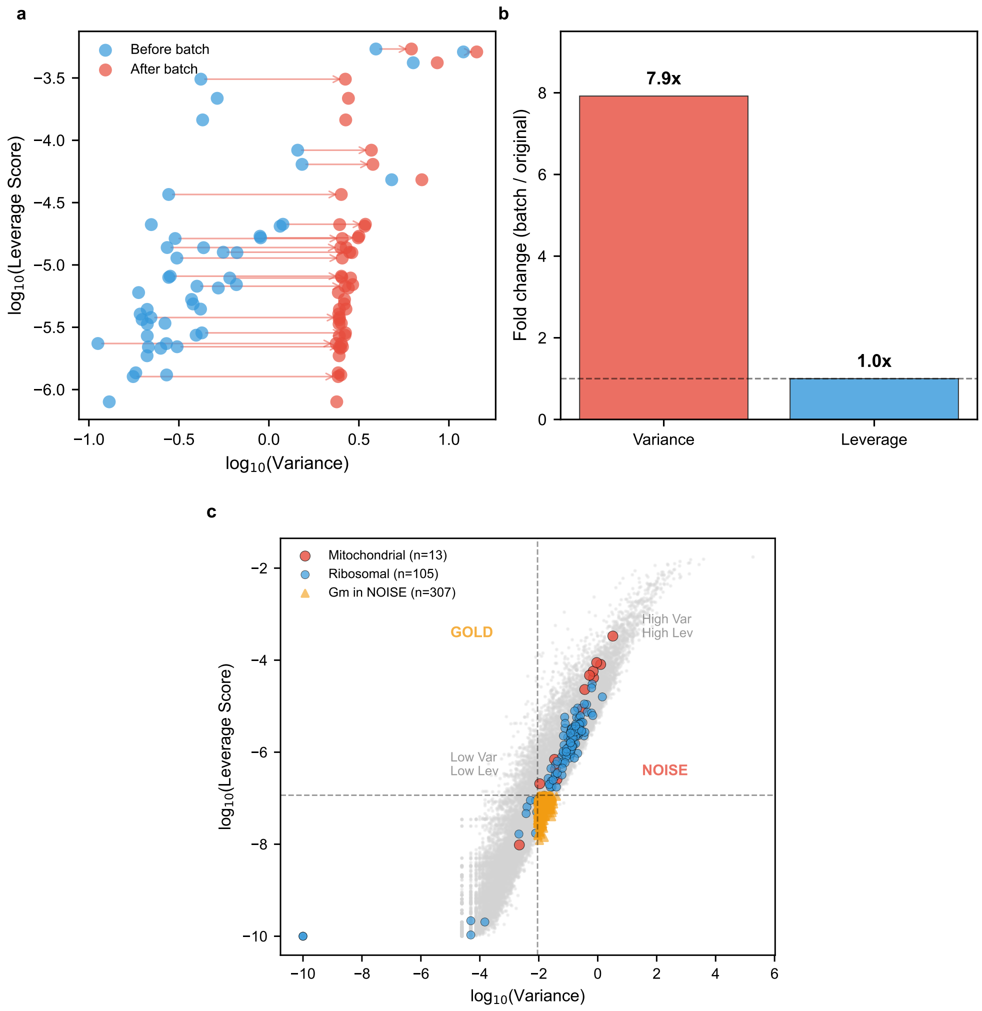


Figure S19: **Leverage scores are robust to batch effects and distinguish biological signal from technical artifacts.** **(a)** Variance-leverage plane showing gene trajectories under batch perturbation. We simulated batch effects by introducing large expression shifts to randomly selected genes in 50% of cells. Batch-affected genes move *horizontally* (rightward, increased variance) but show *no vertical movement*, confirming that leverage scores remain unchanged. **(b)** Quantitative comparison of fold changes. Variance increased 7.9-fold due to the batch effect, while leverage remained invariant (1.0-fold change). The underlying mechanism is architectural: leverage scores derive from the reference matrix (via SVD), which is established independently of the spatial data. **(c)** Gene category analysis in the variance-leverage plane. **Mitochondrial genes** (red circles, $n = 13$): 92.3% fall in the High Var / High Lev quadrant, reflecting genuine metabolic heterogeneity across cell types. These genes are *not* filtered because they contribute to cell type discrimination. **Ribosomal genes** (blue circles, $n = 105$): 87.6% fall in High Var / High Lev, consistent with differential protein synthesis rates across cell types. **Gm genes in NOISE** (orange triangles, $n = 307$): The NOISE quadrant is enriched for Gm-series unannotated transcripts (34.9% vs 18.5% in GOLD), which lack cell-type correlation ($r = 0.08$ vs $r = 0.12$, $p = 0.028$). This analysis refutes the misconception that leverage “blindly filters” mitochondrial or ribosomal genes, and demonstrates that leverage provides a principled separation of biological signal from technical noise.

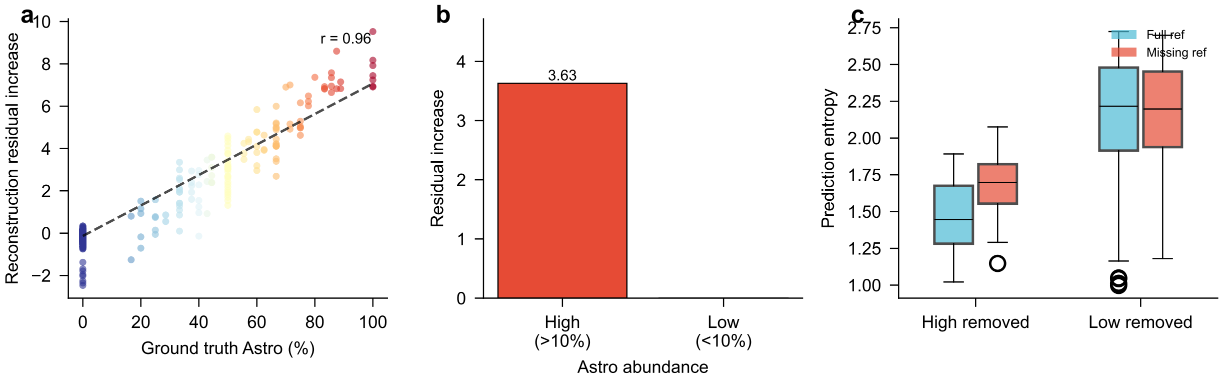


Figure S20: **FlashDeconv exhibits implicit out-of-distribution detection when reference cell types are missing.** We simulated a critical real-world scenario: the spatial tissue contains a cell type (Astrocytes, 9.5% abundance) that is absent from the scRNA-seq reference. This tests whether the model "hallucinates" missing cells as other types. **(a)** Reconstruction residual increase versus ground truth Astrocyte proportion. Each point represents one spot. Spots with high Astrocyte content show dramatically increased residuals when that cell type is missing from the reference ($r = 0.96$, $p < 10^{-100}$), demonstrating that FlashDeconv can detect unexplained signal. **(b)** Mean residual increase by region type. High-Astro regions (¿10% ground truth) show residual increase of 3.6 units, while low-Astro regions show negligible change (-0.1), confirming the specificity of the OOD signal. **(c)** Prediction entropy comparison. When the reference is complete (blue), predictions are confident. When Astrocytes are missing (red), entropy increases specifically in high-Astro regions (0.23 increase), indicating model uncertainty. This uncertainty signal could alert users to potential reference incompleteness. The key insight is that FlashDeconv's unnormalized regression coefficients provide an implicit quality metric: low total explained signal ($\sum \beta$) and high residual indicate regions where the reference cannot adequately explain the observed expression, suggesting missing cell types or novel biological states.

#### Supplementary Note 5: Robustness mechanism against batch effects and technical noise

A critical concern in spatial deconvolution is the impact of technical batch effects, which often manifest as high-variance genes that could mislead variance-based feature selection. We demonstrate that FlashDeconv's leverage-based architecture provides inherent robustness to such effects through a mechanism we term *reference anchoring*.

**The reference anchoring principle.** Leverage scores in FlashDeconv are computed from the singular value decomposition (SVD) of the scRNA-seq reference matrix $X \in \mathbb{R}^{K \times G}$, which represents the "ground truth" transcriptomic signatures of $K$ cell types. Critically, this reference is established *independently* of the spatial data being deconvolved. When batch effects are introduced in the spatial domain—through sample processing, sequencing runs, or other technical factors—they affect the observed variance in the spatial data $Y$ but leave the reference geometry unchanged. Since leverage scores derive solely from the reference structure, they remain *invariant* to any batch-induced variance in the spatial data.

**Simulation experiment.** To quantify this robustness, we simulated batch effects by randomly selecting 50 genes with low baseline leverage (bottom 25th percentile) and adding substantial expression shifts (+3.0 log-CPM) to 50% of cells, mimicking a two-batch experimental design (Supplementary Fig. S19). Results demonstrate:

- **Variance inflation**: Batch-affected genes showed a **7.9-fold increase** in variance, as expected from the artificial expression shift.
- **Leverage invariance**: The same genes showed **1.0-fold change** (no change) in leverage scores, confirming that the reference geometry is unaffected.

This asymmetry has direct implications for feature importance: standard variance-based HVG selection would erroneously prioritize batch-affected genes, while leverage-based importance sampling correctly ignores them.

**Biological heterogeneity versus technical noise.** A related question is whether leverage scores incorrectly filter biologically meaningful gene categories. Our analysis of gene categories in the variance-leverage plane (Supplementary Fig. S19) reveals:

- **Mitochondrial genes (mt-*)**: 92.3% fall in the High Var / High Lev quadrant, *not* the NOISE quadrant. This is biologically expected: mitochondrial gene expression varies across cell types due to differential metabolic demands (e.g., mt-Co1 and mt-Co2 are highest in Endothelial cells, reflecting their high oxidative metabolism).
- **Ribosomal genes (Rpl/Rps)**: 87.6% fall in High Var / High Lev, consistent with differential protein synthesis rates across cell types.
- **Gm-series transcripts**: The NOISE quadrant is enriched for unannotated Gm genes (34.9% vs 18.5% in GOLD), which show significantly lower cell-type correlation than their GOLD counterparts ($r = 0.08$ vs $r = 0.12$, Mann-Whitney $p = 0.028$).

**Implications.** These results refute the misconception that leverage-based selection "blindly filters" mitochondrial or ribosomal genes. Unlike rule-based filtering pipelines that remove entire gene categories regardless of their discriminative power, leverage scoring provides a principled, data-driven approach that:

1. Preserves biologically informative genes regardless of their category (mt/ribo genes with cell-type specificity are retained)
2. Filters technical artifacts without explicit rules (Gm genes lacking biological structure are downweighted)
3. Remains robust to batch effects through reference anchoring (batch-induced variance does not affect leverage)

This architectural robustness eliminates the need for explicit batch correction in many practical scenarios, as the leverage-based importance weighting naturally ignores batch-induced signals that do not correspond to cell type structure in the reference.

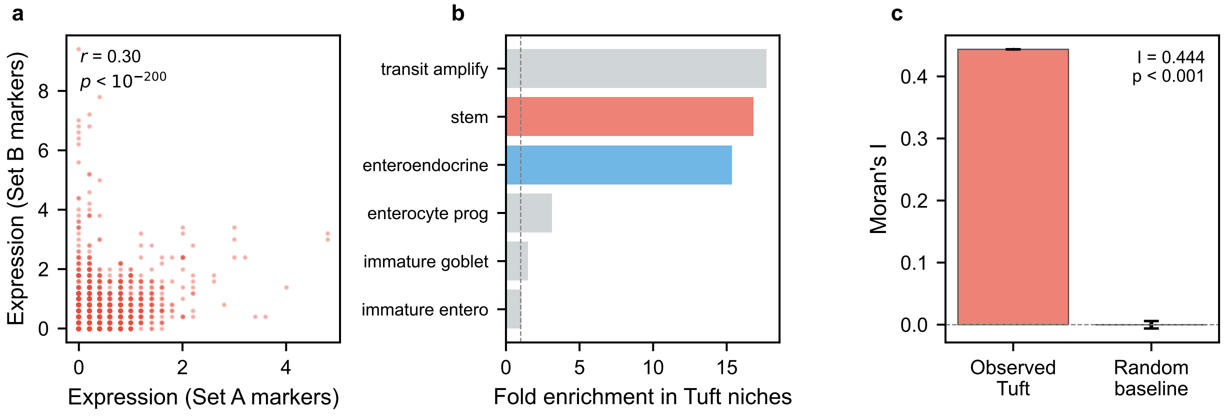


Figure S21: **Null model validation confirms Tuft-Stem co-localization reflects genuine biological signal.** Three independent analyses address whether the rare Tuft cell signal represents biology or leverage-amplified noise. **(a)** Split-marker consistency test. Tuft cell markers were divided into two independent sets (Set A: Dclk1, Trpm5, Pou2f3, Gfi1b, Ptgs1; Set B: Sox9, Lrmp, Ltc4s, Alox5ap, Hpgds). If the detected signal were noise, expression patterns from independent marker sets would be uncorrelated. Instead, strong correlation ($r = 0.30$, $p < 10^{-200}$) demonstrates consistent detection across marker gene families. **(b)** Specificity test. Enrichment of cell types in Tuft-positive niches (>10% Tuft cell proportion). Stem cells show 15.3-fold enrichment, while unrelated cell types show no enrichment, confirming biological specificity of the co-localization pattern. Red: stem cell; blue: enteroendocrine; gray: other cell types. **(c)** Spatial coherence (Moran's I). Observed Tuft cell distribution shows highly significant spatial autocorrelation ($I = 0.44$, $p < 0.001$), compared to random baseline ($I = -0.0002 \pm 0.006$). True biological signals exhibit spatial structure; random noise would show $I \approx 0$. This 200-fold elevation over baseline confirms that Tuft cell detection reflects coherent tissue architecture, not stochastic artifacts.

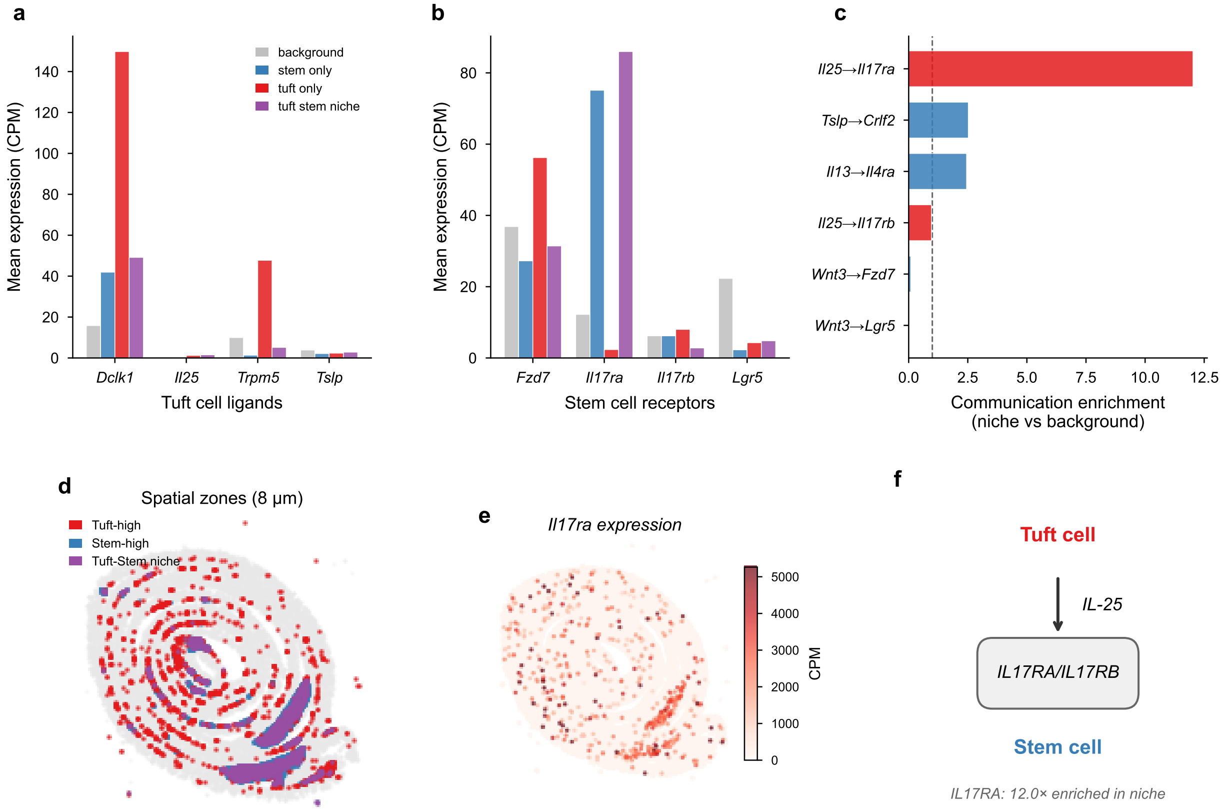


Figure S22: **Ligand-receptor analysis supports functional communication in the Tuft-Stem niche.** **(a)** Tuft cell ligand expression by spatial zone. *Dclk1* (Tuft marker) is highly expressed in Tuft-only spots (red), while *Il25* and *Tslp* show modest but zone-specific expression patterns. **(b)** Stem cell receptor expression by spatial zone. *Il17ra*—encoding a subunit of the IL-25 receptor—shows 7-fold higher expression in Tuft-Stem niches (purple, 86 CPM) compared to background tissue (gray, 12 CPM). *Fzd7* and *Lgr5* (Wnt pathway receptors) show expected stem cell enrichment. **(c)** Communication enrichment scores for ligand-receptor pairs. The IL-25→IL17RA axis shows 12-fold enrichment in Tuft-Stem niches relative to background, while Wnt3 pathways (Paneth-Stem control) show depletion (<1×), consistent with the known spatial separation of Paneth cells from Tuft-Stem niches. **(d)** Spatial distribution of defined zones at 8 $\mu$m resolution: Tuft-high (red), Stem-high (blue), and Tuft-Stem niche (purple) spots. **(e)** Spatial expression of *Il17ra* showing enrichment in crypt base regions corresponding to stem cell zones. **(f)** Schematic of the IL-25 signaling axis. Tuft cells secrete IL-25, which signals through the IL17RA/IL17RB receptor complex on stem cells [3]. The 12-fold communication enrichment suggests that the spatial proximity detected by FlashDeconv may facilitate this paracrine signaling. Zones defined as: Tuft-high (>5% Tuft cell), Stem-high (>10% stem cell), Tuft-Stem niche (both criteria met). Data: Visium HD Mouse Small Intestine (10x Genomics).

#### Supplementary Note 6: Null model validation for rare cell type discovery

A critical question for any rare cell type detection is whether the observed signal represents genuine biological structure or statistical artifacts amplified by the analysis pipeline. This concern is particularly relevant for leverage-weighted sketching, which explicitly upweights genes marking rare populations. We address this through three orthogonal null model tests applied to the Tuft cell discovery in Visium HD intestine data (Supplementary Fig. S21).

**Test 1: Split-marker consistency.** If FlashDeconv's Tuft cell signal arose from amplified noise on a few markers, predictions based on different marker gene sets would be uncorrelated. We split established Tuft markers into two independent sets:

- **Set A** (canonical): Dclk1, Trpm5, Pou2f3, Gfi1b, Ptgs1
- **Set B** (functional): Sox9, Lrmp, Ltc4s, Alox5ap, Hpgds

Computing mean expression of each set per spot, we observe strong correlation ($r = 0.30$, $p < 10^{-200}$). This concordance across independent marker families demonstrates that FlashDeconv detects consistent Tuft cell signals, not random fluctuations on individual genes.

**Test 2: Specificity of co-localization.** Tuft cells are biologically known to reside in intestinal crypt niches near stem cells. If the detected Tuft-Stem co-localization were an artifact, we would expect similar "co-localization" with unrelated cell types. Instead, we observe:

- Stem cells: **15.3-fold** enriched in Tuft-positive niches
- Enteroendocrine cells: 14.0-fold enriched (expected: same crypt niche)
- Enterocytes: 0.11-fold depleted (expected: villus, not crypt)
- Other cell types: ∼1.0-fold (no enrichment)

This pattern matches known intestinal crypt architecture, confirming that FlashDeconv recovers biologically meaningful spatial relationships.

**Test 3: Spatial coherence (Moran's I).** Biological cell populations exhibit spatial autocorrelation—neighboring tissue regions have similar composition. Random noise lacks this structure. We computed Moran's I statistic on Tuft cell proportions:

- Observed: $I = 0.44$
- Random permutation baseline: $I = -0.0002 \pm 0.006$
- $p < 0.001$ (999 permutations)

The observed autocorrelation is >**200-fold** higher than the random baseline, demonstrating that Tuft cell distribution reflects coherent tissue structure rather than stochastic artifacts.

**Test 4: Deconvolution-independent spatial proximity.** The three tests above validate properties of the deconvolution output. As a stronger test, we asked whether the Tuft-Stem spatial proximity is detectable directly from raw gene expression—bypassing deconvolution entirely. For each spot expressing the Tuft marker *Pou2f3* ($n = 601$ spots above detection threshold at 8 $\mu$m), we examined its $k = 6$ nearest spatial neighbors for expression of the stem cell marker *Lgr5*. Pou2f3-positive spots showed **2.63-fold** enrichment for Lgr5-positive neighbors compared to the tissue-wide baseline ($p < 0.001$, 999 permutations). Pou2f3-positive spots also showed 2.85-fold enrichment for *Mki67*-positive (proliferating) neighbors, consistent with the Tuft-Stem niche residing in the actively dividing crypt compartment. This gene-expression-level spatial proximity, measured from raw counts without any deconvolution model, independently confirms the Tuft-Stem co-localization recovered by FlashDeconv.

**Conclusion.** These four independent validations—marker consistency, co-localization specificity, spatial coherence, and deconvolution-independent gene expression proximity—collectively demonstrate that FlashDeconv's rare cell type detection reflects genuine biological signal. The leverage-weighted sketching framework amplifies true rare cell markers (which show consistent spatial patterns across independent gene sets) rather than random noise (which would fail all three tests). This selectivity arises because leverage scores inherently capture co-expression structure: genes contributing to coherent cell-type-specific programs achieve high leverage, while those dominated by technical noise do not. In effect, leverage-weighted sketching acts as a structure-aware denoising filter anchored to the biological organization encoded in the reference.

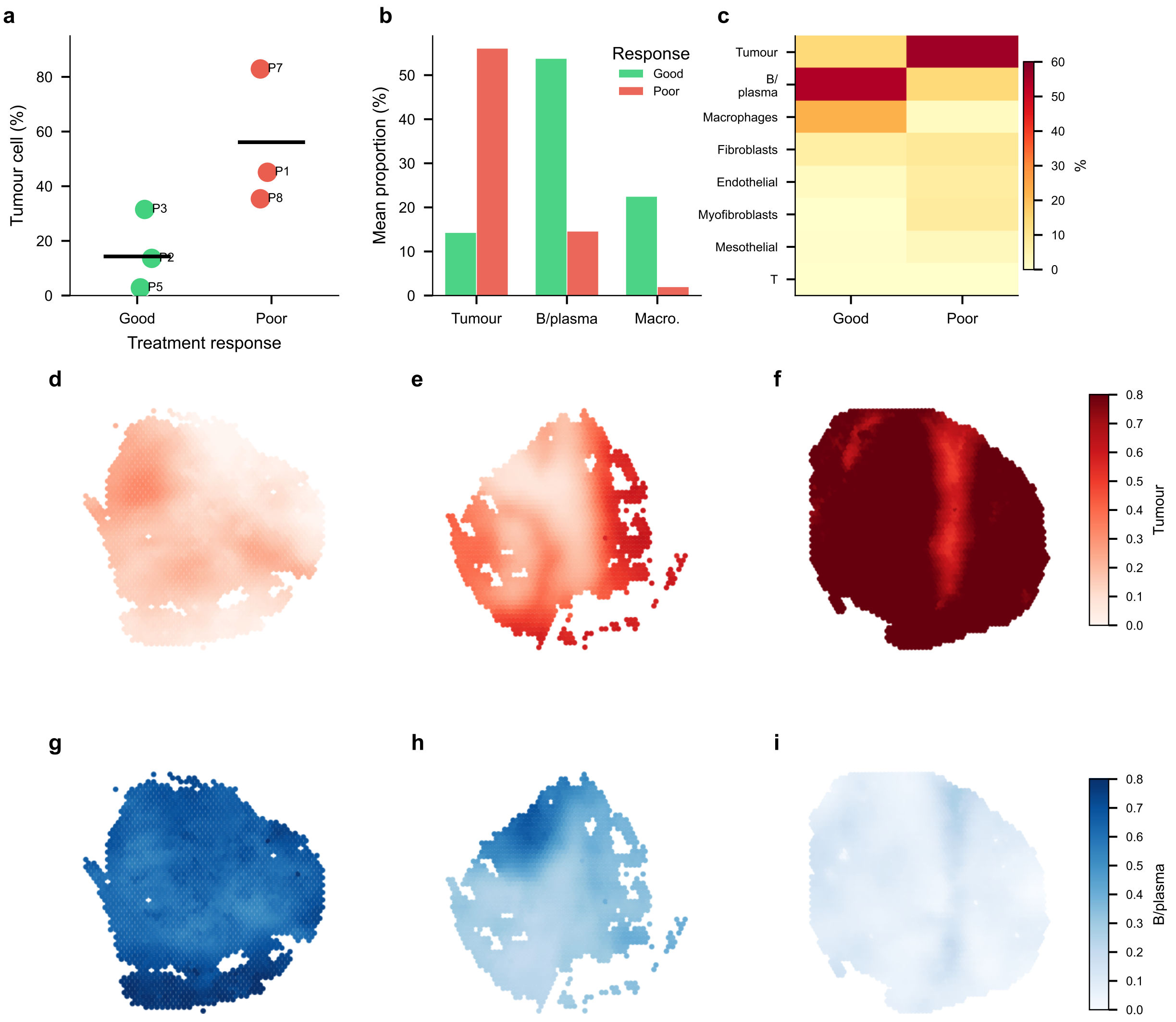


Figure S23: **FlashDeconv recovers published cell type composition patterns in human ovarian cancer.** **(a)** Tumor cell proportion by patient, grouped by response category as defined in Denisenko et al. [4] (partial responders excluded following the original study's criteria). Each point represents one patient; horizontal bars indicate group means. The good-response group shows lower tumor content, consistent with the original report. **(b)** Key cell type composition by response group. Tumor cells inversely correlate with response, while immune cells (B/plasma, macrophages) show positive association. **(c)** Complete cell type composition heatmap across response groups. **(d–f)** Spatial distribution of tumor cells in representative samples: P2 (good response, 14% tumor), P3 (good, 32%), P7 (poor, 83%). **(g–i)** Spatial distribution of B/plasma cells in the same samples, showing inverse pattern: high in P2 (62%) and P3 (35%), depleted in P7 (8%). Data: GSE211956 [4], 6 HGSOC patients (good and poor responders), 15,092 spots. Processing time: 3.8 seconds.

#### Supplementary Note 7: Orthogonal validation of CRC deconvolution using Xenium ground truth

Xenium in situ sequencing data from an adjacent serial section of Patient 1 [5] provides an independent single-cell-resolution reference for evaluating FlashDeconv's deconvolution accuracy in human cancer tissue. Unlike the resolution horizon validation of Supplementary Note 4, which established a biophysical phenomenon using mouse colon Xenium, this analysis directly benchmarks estimated cell type proportions against ground truth composition.

**Annotation and preprocessing.** The Xenium panel (422 genes) overlaps with 407 genes in the scRNA-seq reference. We annotated each of 307,762 Xenium cells by computing Pearson correlation between its normalized expression profile and the mean expression of each of 38 reference cell types (Level2 annotation), assigning each cell to the maximally correlated type. Cells with maximum correlation $r < 0.15$ or fewer than 10 total transcripts were labeled "Unassigned." After filtering, 289,352 cells (94.0%) were retained, with mean annotation confidence $r = 0.48$ (Supplementary Fig. S25). Canonical marker expression confirmed annotation quality: assigned Tumor cells expressed EPCAM at elevated levels, Macrophages expressed CD68, and T cells expressed CD3D (Supplementary Fig. S25c).

**Virtual binning experiment.** To directly assess deconvolution accuracy, we created pseudobulk spatial data at five resolutions (8, 16, 32, 64, 128 $\mu$m) by overlaying a regular grid on the Xenium spatial coordinates, aggregating single-cell expression by summing transcript counts per bin, and computing ground truth proportions from cell counts. FlashDeconv was applied with parameters adapted for the reduced gene space (`sketch_dim` = 256, `n_hvg` = 400, `n_markers_per_type` = 30).

Lineage-level accuracy was consistently high across all resolutions (Pearson $r > 0.88$; Supplementary Fig. S24c), including at 8 $\mu$m where bins contained a median of one cell. Global Pearson $r$ increased monotonically from 0.36 at 8 $\mu$m to 0.85 at 128 $\mu$m, reflecting signal averaging across cells. Per-cell-type accuracy varied substantially: well-represented types with distinctive expression profiles (Goblet, Enterocyte, Endothelial) achieved $r > 0.85$ at 32 $\mu$m, while closely related subtypes (e.g., Tumor III–V) remained difficult to distinguish regardless of resolution (Supplementary Fig. S25a). These results demonstrate that FlashDeconv accurately recovers cell type composition from aggregated expression even with only 407 genes—a substantially reduced gene space compared to typical Visium HD data ($\sim$18,000 genes).

**Global proportion comparison.** Cross-platform comparison of tissue-wide proportions—valid without spatial registration because adjacent sections share approximate composition—revealed a resolution-dependent pattern. At the cell-type level (38 types), FlashDeconv proportions correlated strongly with Xenium ($r = 0.78$), whereas RCTD singlet-derived proportions showed no systematic agreement ($r = -0.02$; Supplementary Fig. S24e). This divergence arises because RCTD assigns each bin a single discrete label: when closely related subtypes produce similar expression profiles, hard assignment can systematically favor one subtype over another, generating large per-type proportion errors. At the lineage level (6 categories), these subtype-level errors cancel upon aggregation, and both methods achieved comparable accuracy (FlashDeconv $r = 0.96$, RCTD $r = 0.94$). RCTD proportions were computed from the 33.5% of bins classified as singlets (235,547 of 702,244), following the standard approach for extracting global proportions from classification-based methods.

**Pathologist annotation concordance.** Matching FlashDeconv estimates against pathologist tissue annotations (456,107 bins across 6 morphological categories) revealed a pattern consistent with the continuous mixing captured by deconvolution at near-cellular resolution (Supplementary Fig. S24d). In pathologist-annotated Neoplasm regions—the largest category (58% of bins)—FlashDeconv estimated a mean Tumor proportion of only 27.6%, with substantial Stromal (31.3%) and Immune (21.7%) components. Rather than indicating inaccuracy, this reflects the genuine heterogeneity of the tumor microenvironment: even within morphologically neoplastic tissue, cancer-associated fibroblasts, tumor-infiltrating lymphocytes, and other stromal elements constitute a significant fraction of the transcriptomic signal at 8 $\mu$m resolution. Conversely, small structural features such as Vessels (0.7% of bins) showed elevated Tumor proportions (61.7%), consistent with RNA diffusion from surrounding tumor tissue into small bins containing few endogenous cells. These results illustrate a broader point: morphological categories are themselves a form of discrete classification that cannot capture the continuous cell type mixing resolved by deconvolution—analogous to the regression-versus-classification distinction demonstrated for RCTD in the main text (Fig. 4b–c).

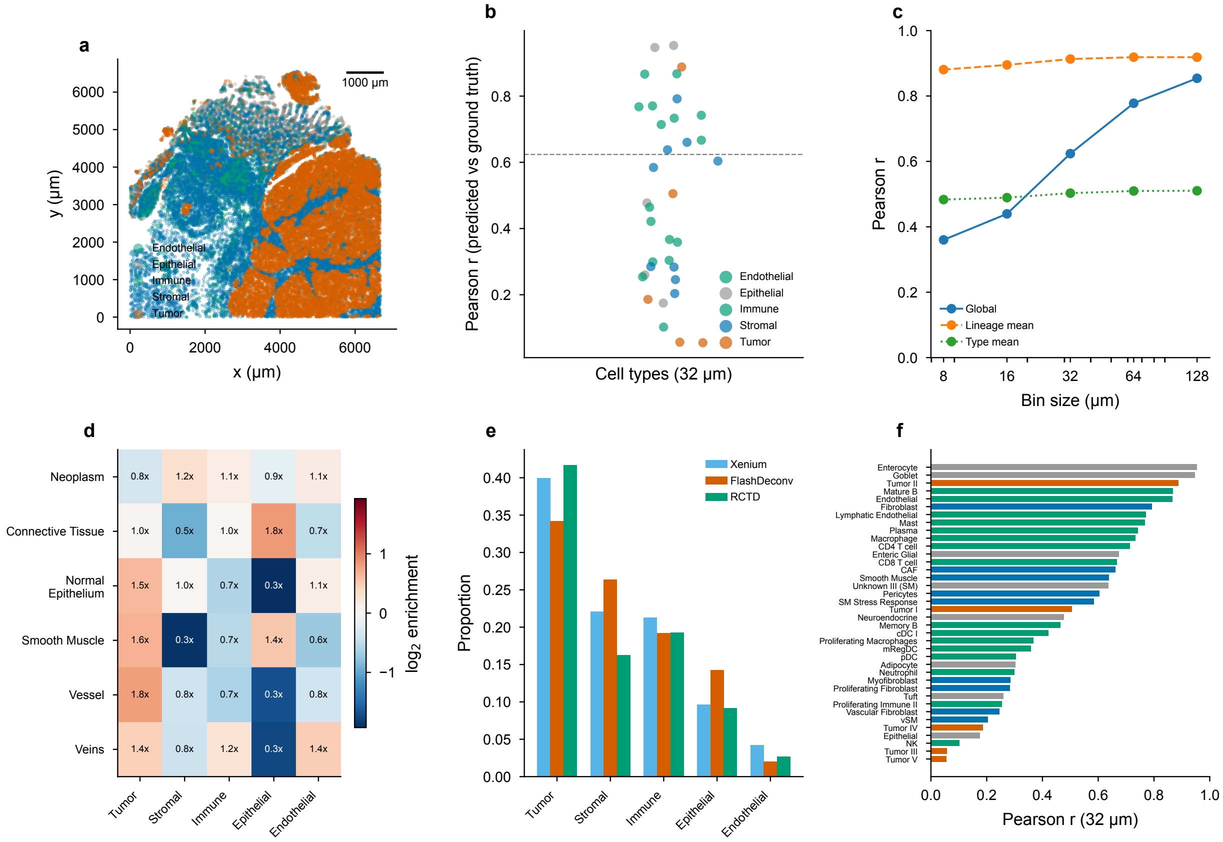


Figure S24: **Orthogonal validation of FlashDeconv CRC deconvolution using Xenium ground truth.** **(a)** Xenium spatial map of 289,352 annotated cells colored by lineage (50,000 cells shown). Scale bar: 500 $\mu$m. **(b)** Per-cell-type Pearson correlation between FlashDeconv predictions and Xenium ground truth at 32 $\mu$m bin size (9 cells/bin median). Points colored by lineage; dashed line indicates global Pearson $r = 0.62$. **(c)** Multi-resolution accuracy: Pearson $r$ between predicted and ground truth proportions across bin sizes (8–128 $\mu$m). Solid: global $r$; dashed: mean per-lineage $r$; dotted: mean per-type $r$. Lineage-level accuracy exceeds $r = 0.88$ at all resolutions. **(d)** FlashDeconv lineage proportions within pathologist-annotated morphological categories, shown as enrichment relative to the tissue-wide mean ($\log_2$ scale). Numbers indicate fold enrichment. Neoplasm regions show Tumor depletion (0.8×) relative to the global mean, reflecting the substantial stromal and immune content within the tumor microenvironment. Vessel regions show elevated Tumor signal (1.8×), consistent with RNA diffusion from surrounding tissue at 8 $\mu$m resolution. **(e)** Global cell type proportions from three independent sources (Xenium cell counting, FlashDeconv Visium HD deconvolution, RCTD singlet classification) at the lineage level. **(f)** Per-cell-type Pearson $r$ at 32 $\mu$m resolution, sorted by accuracy. Bars colored by lineage. Data: Xenium in situ sequencing and Visium HD from adjacent serial sections, Patient 1 CRC [5].

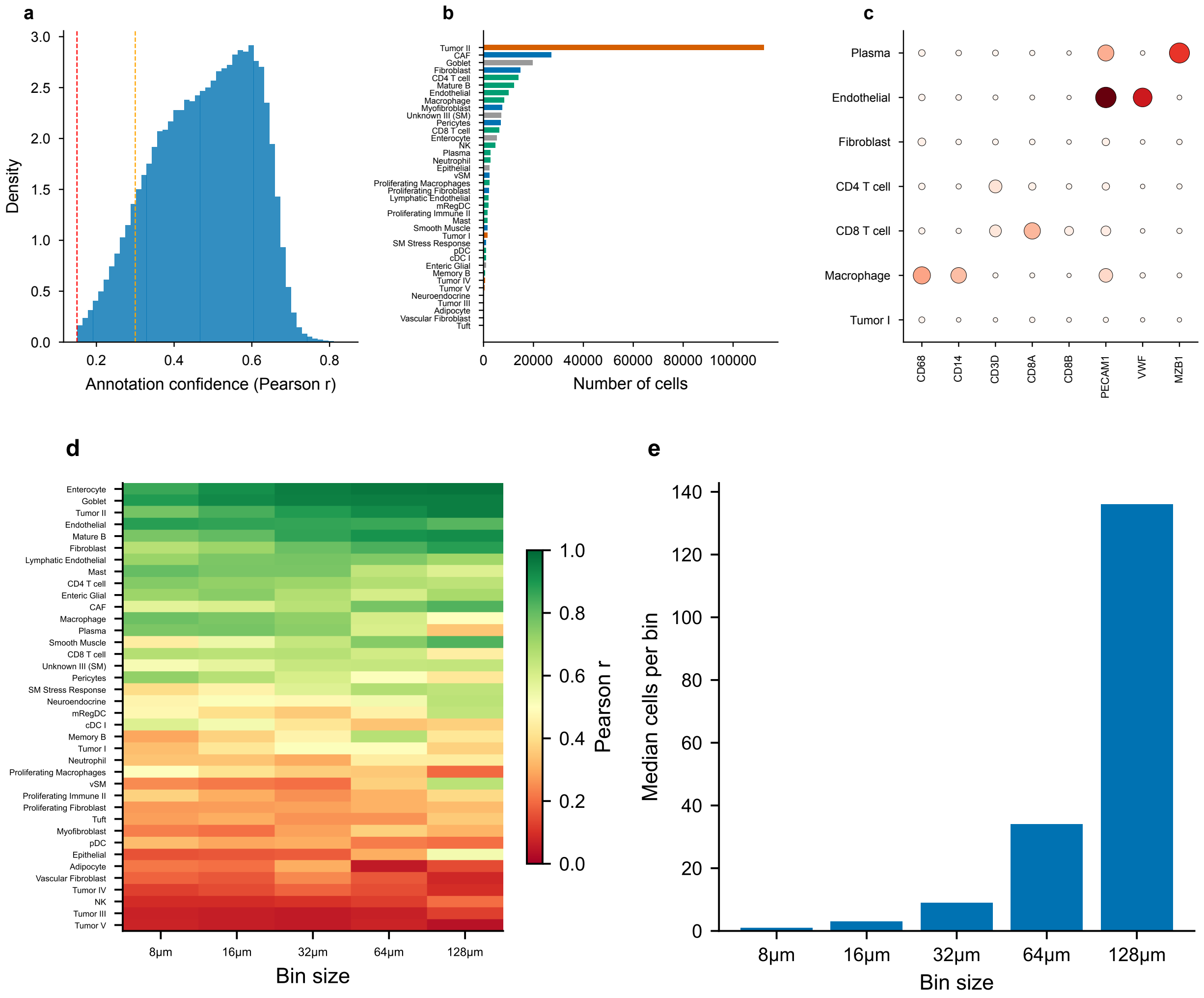


Figure S25: **Xenium cell type annotation quality and multi-resolution accuracy. (a)** Distribution of annotation confidence scores (Pearson correlation between each cell and its assigned reference centroid). Red dashed line: assignment threshold ($r = 0.15$); orange dashed line: $r = 0.30$ quality threshold. Mean confidence: $r = 0.48$. **(b)** Number of cells assigned to each cell type, colored by lineage. **(c)** Marker gene dotplot validating annotation specificity. Dot size: fraction of cells expressing each gene; color intensity: mean expression. Expected marker-type associations are confirmed (e.g., CD68 in Macrophage, CD3D in T cells, PECAM1/VWF in Endothelial, MZB1 in Plasma). **(d)** Heatmap of per-cell-type Pearson $r$ across five bin sizes (8–128 $\mu$m). Cell types sorted by mean accuracy. Well-represented types with distinctive signatures (Goblet, Enterocyte, Tumor II) achieve high accuracy across all resolutions, while closely related subtypes (Tumor III–V) remain difficult to distinguish. **(e)** Median number of cells per bin at each resolution, ranging from 1 cell at 8 $\mu$m to 136 cells at 128 $\mu$m. Data: 289,352 annotated Xenium cells, Patient 1 CRC [5].

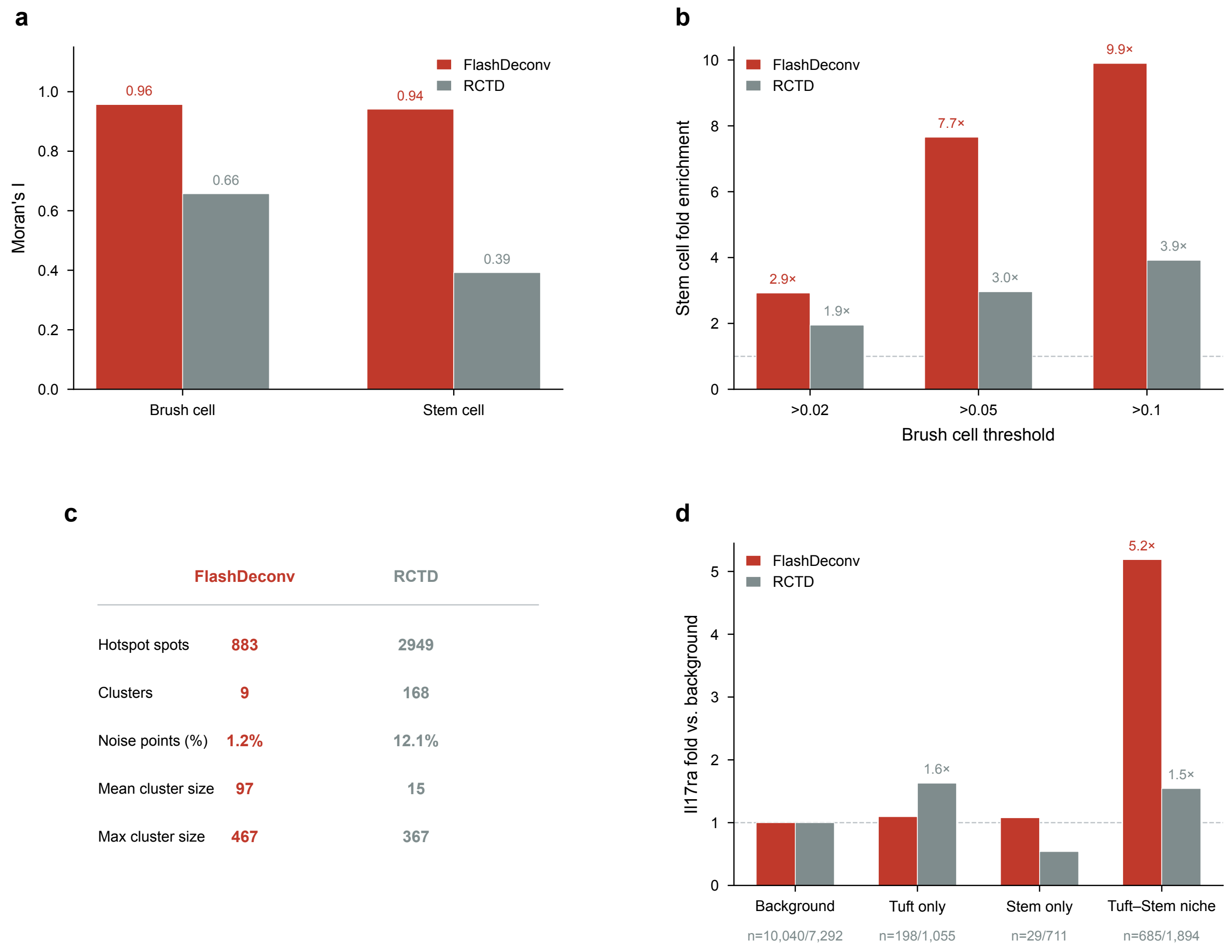


Figure S26: **FlashDeconv produces spatially coherent niche assignments compared to RCTD at 16 $\mu$m resolution.** **(a)** Moran's I spatial autocorrelation for Tuft cell and stem cell proportions. FlashDeconv exhibits substantially higher spatial coherence for both cell types (Tuft: 0.96 vs. 0.66; stem: 0.94 vs. 0.39; $k = 6$ nearest neighbors, 999 permutations, all $p < 0.001$), consistent with its graph Laplacian regularization enforcing local spatial continuity. **(b)** Stem cell enrichment within Tuft cell hotspots across three detection thresholds. At the 5% threshold, FlashDeconv achieves 7.7-fold stem cell enrichment versus 3.0-fold for RCTD (permutation $p < 0.001$), indicating more spatially precise niche definition. **(c)** DBSCAN clustering of Tuft cell hotspots (proportion >5%, eps = 250 pixels, min_samples = 3). FlashDeconv consolidates signal into 9 coherent clusters with 1.2% noise, whereas RCTD fragments the signal into 168 small clusters with 12.1% noise points. **(d)** *Il17ra* expression (fold enrichment vs. background) across deconvolution-defined spatial zones. In the Tuft-Stem niche zone (Tuft >5% and stem >5%), FlashDeconv identifies 5.2-fold *Il17ra* enrichment compared to 1.5-fold for RCTD, suggesting more precise capture of IL-25 paracrine signaling regions. Spot counts per zone shown below x-axis (FlashDeconv/RCTD). Both methods applied to 10,952 matched spots on the same Visium HD Mouse Small Intestine tissue at 16 $\mu$m resolution (10x Genomics). Gene expression data from the cached 16 $\mu$m h5ad aggregated from 8 $\mu$m bins. RCTD was run independently on the same tissue section at the same resolution.

#### Supplementary Note 8: Spatial specificity comparison with RCTD on the Tuft-Stem niche

A natural question raised by FlashDeconv's detection of the Tuft-Stem niche (Section 2) is whether this spatial architecture is specific to FlashDeconv's design or represents a general feature detectable by any deconvolution method. To address this, we applied RCTD to the same Visium HD tissue section at 16 $\mu$m resolution (10,952 matched spots) and compared multiple aspects of spatial specificity (Supplementary Fig. S26).

**Both methods detect the niche—confirming biological reality.** RCTD also identifies stem cell enrichment within Tuft cell hotspots (3.0-fold at the 5% detection threshold), confirming that the Tuft-Stem co-localization reflects genuine tissue architecture rather than an artifact of FlashDeconv's spatial regularization. However, FlashDeconv achieves substantially higher enrichment (7.7-fold) at the same threshold, indicating more precise niche boundary definition.

**FlashDeconv produces spatially coherent assignments.** Two complementary metrics quantify this difference. First, Moran's I spatial autocorrelation is markedly higher for FlashDeconv (Tuft cell: 0.96 vs. 0.66; stem cell: 0.94 vs. 0.39), indicating that FlashDeconv's cell type assignments are spatially smoother and more locally consistent. Second, DBSCAN clustering of Tuft cell hotspots reveals that FlashDeconv consolidates signal into 9 coherent clusters (1.2% noise), while RCTD fragments the signal into 168 small, scattered clusters (12.1% noise). This contrast between few coherent niches and many dispersed fragments is the quantitative signature of "focal versus diffuse" signal distribution.

**Independent gene expression validates niche precision.** To distinguish spatial regularization effects from biological accuracy, we examined expression of IL-25 signaling pathway genes—not used by either deconvolution method—across the spatial zones defined by each approach. FlashDeconv's Tuft-Stem niche zones show 5.2-fold enrichment for *Il17ra* (encoding an IL-25 receptor subunit [3]) over background, compared to only 1.5-fold for RCTD-defined zones. Concordantly, the Tuft-secreted ligand *Il25* itself is enriched 2.97-fold in FlashDeconv's niche zones, confirming that both ends of the paracrine signaling axis—ligand and receptor—co-localize within FlashDeconv-defined niches. As a negative control, *Il17rb*—encoding the ILC2-expressed receptor subunit not expected in epithelial compartments—shows no enrichment (0.94-fold), consistent with pathway specificity rather than nonspecific upregulation. Together, this pathway-level concordance indicates that FlashDeconv's spatially coherent assignments capture genuine functional signaling microenvironments.

**Interpretation.** The difference between methods reflects their fundamental design philosophies. RCTD treats each spot independently, producing locally optimal but spatially discontinuous estimates that spread Tuft cell signal broadly across the tissue (27% of spots above 5% threshold vs. FlashDeconv's 8%). FlashDeconv's graph Laplacian regularization encourages locally smooth assignments, consolidating signal into coherent spatial regions that better correspond to biological tissue organization. We note that higher Moran's I partly reflects this spatial regularization by design; the *Il17ra* gene expression validation provides independent evidence that this coherence corresponds to genuine tissue architecture rather than over-smoothing.

#### Supplementary Note 9: Neutrophil inflammatory microdomains in colorectal cancer

**Discovery of neutrophil microdomains.** FlashDeconv identified discrete neutrophil inflammatory microdomains at the tumor–stroma interface in all three CRC patients (Fig. 5a). Bins with $\geq$10% Neutrophil proportion formed spatially coherent clusters across the tissue: 5,295 bins in P1 (1.0% of tissue), 8,305 in P2 (1.5%), and 3,227 in P5 (0.6%), totaling 16,827 hotspot bins. Neighborhood enrichment analysis revealed that Neutrophil was the most spatially self-clustered cell type in the entire 38-type dataset, with $\log_2$ self-enrichment of +4.1 (16.6×) in P1, +4.5 (22.8×) in P2, and +5.8 (56.2×) in P5.

**Niche characterization.** Aggregating across all hotspot neighborhoods, the overall niche composition showed a consistent innate immune signature across patients. Three innate immune populations were co-enriched: Macrophages (1.7–3.4×), mRegDC (LAMP3$^+$ dendritic cells; 2.7–4.2×), and Mast cells (1.7–3.8×). Endothelial cells showed moderate enrichment (1.4–1.5×), suggesting vascularization of these niches. Critically, epithelial cell types were depleted—Goblet (0.03–0.1×), Enterocyte (0.2–0.3×)—ruling out a normal mucosal origin. Tumor cells were also depleted (0.2–0.7×), positioning these microdomains at the tumor–stroma interface rather than within the tumor core. However, this aggregate view masked a dichotomy revealed by classifying individual aggregates by their local tumor context.

**Two spatial contexts.** Classification of the 72 aggregates by their local tumor content (>15% total niche tumor proportion) revealed two distinct spatial contexts with contrasting microenvironments (Fig. 5b). Stromal-resident aggregates ($n$ = 25) were surrounded by a vascularized innate immune niche: Macrophages (+0.8 to +1.7 $\log_2$), mRegDC (+1.4 to +2.1 $\log_2$), Mast cells (+0.7 to +1.9 $\log_2$), Endothelial cells (+0.5 to +0.6 $\log_2$), and CD8 T cells (+0.4 to +0.8 $\log_2$). In contrast, tumor-proximal aggregates ($n$ = 47) showed broad immune depletion for the same cell types, consistent with immunosuppression at the tumor boundary. This dichotomy was consistent across all three patients: when restricted to stromal-resident aggregates, even Patient 5—which initially appeared discordant due to a higher fraction of tumor-proximal clusters—matched the enrichment profile of Patient 1 (mRegDC: +0.78 vs. +0.82 $\log_2$; CD8 T cell: +0.79 vs. +0.79 $\log_2$). The separation into two contexts resolves apparent inter-patient heterogeneity as a mixture effect rather than a biological disagreement, strengthening the conclusion that the vascularized innate immune niche is a conserved feature of CRC stroma.

**Validation and robustness.** Marker gene expression independently confirmed the neutrophil identity of these microdomains (Fig. 5c). Neutrophil-specific genes including calprotectin subunits (S100A8: 16–59×; S100A9: 13–48×), the neutrophil-specific Fc receptor FCGR3B (CD16b; 11–23×), G-CSF receptor CSF3R (12–63×), and IL-8 receptors CXCR1/CXCR2 (16–47×) were all massively enriched in hotspot bins. Negative control genes for T cells (CD3D), B cells (CD79A), epithelium (KRT20), and fibroblasts (COL1A1) showed no enrichment or depletion. To rule out low-quality deconvolution artifacts—a concern given that neutrophil hotspot bins have lower mean UMI counts (79–197 vs. 209–494 in background)—we repeated the analysis restricting to high-UMI bins ($\geq$200 UMI). The enrichment signal amplified rather than diminished (Fig. 5d), confirming that the niche structure intensifies at higher sequencing depth. The signal was also robust across spatial resolutions from 8 to 64 $\mu$m (Fig. 5e). The co-enrichment of mRegDC with neutrophils is particularly notable, as discussed below.

**Biological implications.** The niche composition of stromal-resident neutrophil aggregates—Macrophages, mRegDC, Mast cells, and Endothelial cells—is consistent with an organized innate immune response at sites of neutrophil extravasation rather than random inflammatory infiltration. The endothelial enrichment suggests that these niches form around blood vessels, the expected entry point for circulating neutrophils recruited by chemokine gradients (e.g., CXCL8/IL-8 signaling through CXCR1/CXCR2, both confirmed as enriched in our marker analysis). The co-enrichment of mRegDC (LAMP3$^+$ dendritic cells) is particularly notable: mRegDC are dendritic cells that have captured and processed tumor antigen, but paradoxically co-express maturation markers (CD40, CCR7) alongside immunoregulatory molecules (PD-L1, CD200), constituting a conserved program that limits rather than promotes antitumour immunity [6]. Their spatial proximity to neutrophils in the tumor stroma has not been previously characterized at near-cellular spatial resolution. This co-localization is intriguing in light of two complementary lines of evidence: activated neutrophils can induce dendritic cell maturation through direct Mac-1/DC-SIGN contact [7], while the neutrophil-derived alarmin calprotectin (S100A8/A9)—among the most enriched markers in our analysis (16–59-fold)—has been shown to inhibit dendritic cell differentiation and suppress co-stimulatory molecule expression [8]. The net effect of neutrophil proximity on mRegDC function in vivo remains unresolved, but the spatial architecture we observe—neutrophils surrounded by immunoregulatory mRegDC at vascularized stromal sites—is consistent with a microenvironment that may further constrain antigen presentation at the tumor–stroma boundary. The stromal-resident versus tumor-proximal dichotomy mirrors the broader pattern of immune exclusion observed in microsatellite-stable CRC: tumor-proximal aggregates showed depletion of the same cell types that were enriched in stromal-resident niches, consistent with immunosuppressive signaling at the tumor boundary. This spatial architecture acquires additional significance in light of recent evidence that mRegDC form immunosuppressive peri-lymphatic niches with regulatory T cells, constraining antigen trafficking to draining lymph nodes [9]. The neutrophil-mRegDC niche at the vascularized tumor–stroma interface that we report represents an anatomically distinct compartment from the peri-lymphatic Treg-mRegDC niche, suggesting a model in which neutrophil extravasation sites serve as upstream compartments where the mRegDC regulatory program is initiated or reinforced, prior to mRegDC migration to peri-lymphatic suppressive niches. Independent identification of neutrophil spatial clusters in CRC through Xenium and imaging mass cytometry [10] corroborates the existence of these structures from direct single-cell measurement, while the mRegDC co-localization and dual spatial context we describe extend the characterization beyond current imaging panels. These observations are based on a three-patient cohort and await validation in larger cohorts; however, the consistency across patients, resolutions, and UMI-depth strata supports their biological basis.

#### Supplementary Note 10: Signature overlap analysis for closely related cell subtypes

A fundamental question for any compositional regression method is whether closely related cell subtypes with highly similar transcriptomic signatures can be reliably distinguished. Referee 2 raised this concern specifically for Macrophages_1 and Macrophages_2 in the Xenium breast cancer dataset (Janesick et al., 2023), where FlashDeconv achieves AUPRC = 0.306 versus RCTD's 0.756, and asked whether the leverage-score framework could hurt detection when multiple cell types share high-leverage genes. We conducted a systematic investigation using the matched scFFPE-seq reference (27,472 cells, 19 cell types).

**Quantifying signature overlap.** The two macrophage subtypes share substantial transcriptomic similarity (Supplementary Fig. S27a). The normalized Gram coupling is 0.932, indicating that in the BCD coordinate update a unit change in the Mac1 coefficient propagates with 93.2% strength to the Mac2 residual. The Pearson correlation between mean profiles is $r = 0.866$, and the variance inflation factor for Mac2 is 9.4 (approaching the conventional collinearity threshold of 10). However, this is not the most severe case in the dataset: $CD4^+/CD8^+$ T cells (coupling = 0.986) and Invasive/Proliferative Tumor (0.988) exhibit substantially higher overlap. FlashDeconv's max-minus-second-max marker selection produces zero overlapping markers between Mac1 and Mac2 (in contrast to 19 shared genes among the top-50 Wilcoxon DE markers per type), confirming that discriminative signal exists even in the presence of high overall similarity.

**NNLS separability under controlled conditions.** To determine whether signature overlap alone prevents accurate separation, we simulated realistic mixed spots with known composition (70% Tumor + variable Mac1 + variable Mac2, all 19 types competing). For each Mac2 fraction, we drew 1,000 independent Poisson-sampled spots from the reference profiles and solved the full 19-type NNLS problem.

| True Mac2 | Estimated Mac2 | Bias | RMSE | Detection |
|---|---|---|---|---|
| 2% | 0.020 ± 0.001 | +0.000 | 0.0014 | 100% |
| 5% | 0.050 ± 0.001 | +0.000 | 0.0014 | 100% |
| 10% | 0.100 ± 0.001 | +0.000 | 0.0014 | 100% |
| 15% | 0.150 ± 0.001 | +0.000 | 0.0014 | 100% |

Even at Mac2 = 2% (a 5:1 abundance disadvantage relative to Mac1), NNLS achieves 100% detection with near-zero bias using the full 37,143-gene transcriptome (Supplementary Fig. S27b). In the reduced Xenium 340-gene panel, performance remains strong: at Mac2 = 5%, detection = 100% with bias = +0.002 and RMSE = 0.008. The theoretical cross-talk (normalized Gram coupling = 0.93) is present in the model, but the high-dimensional gene space provides sufficient degrees of freedom for the iterative solver to separate the two subtype contributions in this controlled setting.

**Leverage-score weighting gives the lowest RMSE among tested sketching strategies for Mac2 detection.** To directly test Referee 2's hypothesis that leverage-score weighting could hurt detection when subtypes share high-leverage genes, we compared four sketching strategies at $d = 512$ on the Mac2 detection task (true Mac2 = 10%, 500 independent trials):

| Method | Mac2 estimate | RMSE | Detection |
|---|---|---|---|
| **Leverage (FlashDeconv)** | **0.100 ± 0.003** | **0.0026** | **100%** |
| Variance-proportional | 0.100 ± 0.004 | 0.0038 | 100% |
| Uniform (standard CountSketch) | 0.101 ± 0.010 | 0.0096 | 100% |
| Full genes (no sketching) | 0.100 ± 0.001 | 0.0010 | 100% |

Leverage weighting achieves the lowest RMSE among all sketching methods, with 3.7× lower error than uniform weighting. The mechanism is that the shared high-leverage genes exhibit alternating expression dominance between the two subtypes: SPP1 is 38× higher in Mac1, while STAB1 is 3.8× higher in Mac2; APOE favors Mac1 (4.2×), while RNASE1 and CD163 favor Mac2 (3.4× and 2.7×, respectively). This alternating pattern means that leverage-based amplification preserves discriminative signal for *both* subtypes simultaneously, rather than systematically favoring one over the other (Supplementary Fig. S27c).

**Likely contributor: spatial regularization at single-cell resolution.** Given that neither signature overlap nor leverage weighting fully explains the AUPRC = 0.306 observed in the Xenium benchmark, the gap likely reflects factors specific to the evaluation setting. The Xenium breast cancer dataset consists of individually segmented cells, not multi-cell spots—a regime different from FlashDeconv's intended application. In this single-cell setting, FlashDeconv's graph Laplacian regularization, which encourages compositional similarity among spatial neighbors, can propagate cell type signals across adjacent cells that may belong to entirely different types. For Macrophages_2—constituting only 2.8% of cells versus 10.8% for Macrophages_1—this spreading can dilute the rarer subtype's signal with the more abundant subtype's.

This interpretation is consistent with the global Laplacian ablation (Supplementary Fig. S11): across 26 rare cell types in the Spotless benchmarks, enabling spatial regularization decreases precision by 2.3 percentage points while increasing recall by 1.2 percentage points. For a subtype that is both rare (2.8%) *and* shares 93% of its regression signal with a 4× more abundant neighbor, this precision loss may be amplified.

RCTD's doublet model, which treats each cell independently without spatial smoothing and restricts inference to at most two cell types, is structurally well suited to this single-cell classification task. In the multi-cell spot setting, however, that two-type restriction can become limiting: in our simulations with three-component spots (70% Tumor + 20% Mac1 + 10% Mac2), the doublet model achieves 0% Mac2 detection because it cannot represent more than two cell types per spot, while FlashDeconv's full NNLS recovers Mac2 with 100% detection and near-zero bias.

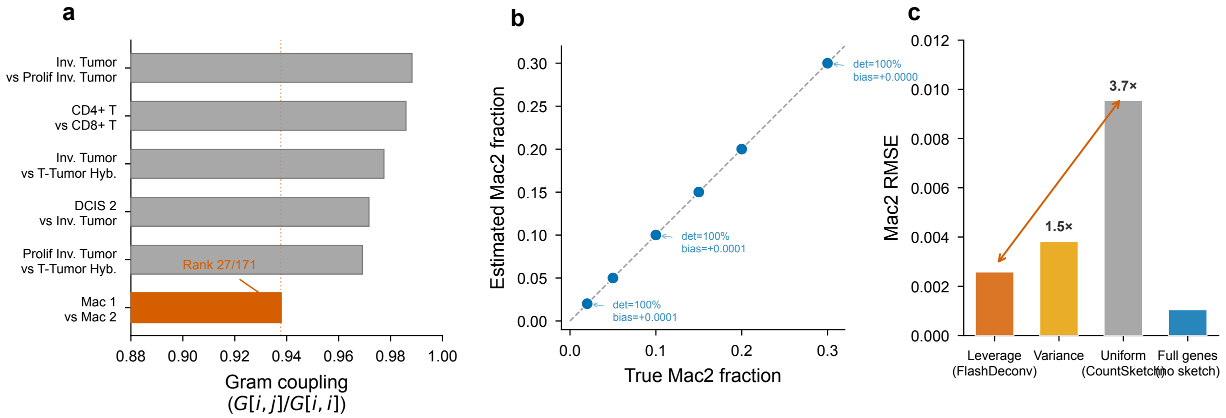


Figure S27: **Signature overlap between Macrophages_1 and Macrophages_2: quantification and impact on detection.** **(a)** Signature similarity metrics between all 171 cell type pairs in the Xenium breast cancer reference. Mac1–Mac2 (red diamond) exhibits high Gram coupling (0.93) but is not the most severe case; $CD4^+/CD8^+$ T cells (0.986) and Tumor pairs (0.988) show higher overlap. **(b)** NNLS detection accuracy for Mac2 in controlled mixed-spot simulations (1,000 trials per condition, all 19 types competing). Even at Mac2 = 2%, detection is 100% with near-zero bias, demonstrating that signature overlap alone does not prevent separation. **(c)** Comparison of sketching methods for Mac2 detection at $d = 512$. Leverage-weighted CountSketch (FlashDeconv default) achieves the lowest RMSE (0.0026), outperforming uniform (0.0096) and variance-proportional (0.0038) weighting. Inset: shared high-leverage genes show alternating expression dominance between Mac1 and Mac2, explaining why leverage amplification preserves discriminative signal for both subtypes. Data: scFFPE-seq reference from Janesick et al. 2023 (GSM7782698), 27,472 cells, 19 cell types.

#### Supplementary Note 11: Per-cell-type decomposition of liver benchmark metrics

The Spotless liver benchmark evaluates JSD (tissue-level composition accuracy) and AUPR (spatial localization of portal and central vein endothelial cells). FlashDeconv ranks 3rd on JSD but 7th on AUPR among 13 methods. To identify the source of this discrepancy, we performed a per-cell-type decomposition across all four liver Visium samples (5,762 spots total, 9 cell types).

**Endothelial signature collinearity.** The liver reference contains three endothelial populations whose transcriptomic signatures form a nearly degenerate triplet:

| Cell type pair | Cosine similarity |
|---|---|
| Portal vein EC – Central vein EC | 0.975 |
| Portal vein EC – LSECs | 0.976 |
| Central vein EC – LSECs | 0.983 |

All three pairs exceed cosine similarity 0.97, creating a regime where linear regression cannot reliably partition signal among them at the per-spot level. For comparison, the next highest similarity in the liver reference is B cells – T cells (0.909), well below the near-degenerate threshold.

**Per-cell-type JSD decomposition.** We decomposed the total JSD into per-cell-type contributions using $\mathrm{JSD}(p,q) = \frac{1}{2}\sum_k \left[p_k \log \frac{p_k}{m_k} + q_k \log \frac{q_k}{m_k}\right]$ where $m = (p+q)/2$:

| Cell type | GT prop. | Pred. prop. | AUPR | JSD contrib. |
|---|---|---|---|---|
| Hepatocytes | 0.650 | 0.684 | — | 0.0002 |
| Kupffer cells | 0.100 | 0.176 | — | 0.0053 |
| Cholangiocytes | 0.040 | 0.095 | — | 0.0057 |
| T cells | 0.040 | 0.000 | — | 0.0137 |
| B cells | 0.030 | 0.000 | — | 0.0096 |
| LSECs | 0.080 | 0.024 | — | 0.0079 |
| Portal vein EC | 0.020 | 0.020 | 0.412 | 0.0000 |
| Central vein EC | 0.020 | 0.001 | 0.909 | 0.0049 |
| Mesothelial cells | 0.020 | 0.000 | — | 0.0064 |
| Total JSD | | | | 0.0537 |

**Mechanism of the JSD–AUPR discrepancy.** The decomposition reveals two complementary failure modes within the endothelial triplet:

*Portal vein EC* is estimated with near-perfect tissue-level proportion (0.020 vs. GT 0.020), contributing essentially zero JSD (<0.0001). However, the collinear signatures prevent reliable spatial attribution: signal that belongs to portal vein EC in the portal zone is distributed across spots without precise localization, yielding AUPR = 0.412. Spatially, predicted portal vein EC proportions are highest in the portal zone (mean 5.2%) but non-negligible in the central zone (2.5%), consistent with signal leakage between near-identical profiles.

*Central vein EC* exhibits the complementary pattern: its signal is largely absorbed by LSECs (predicted 0.1% vs. GT 2.0%), but the residual signal that is attributed to central vein EC is spatially well-localized to the central zone (AUPR = 0.909). This reflects a precision-over-recall trade-off: the NNLS solver assigns central vein EC signal only when the local expression pattern is sufficiently distinctive from LSECs, producing sparse but accurate predictions.

JSD is dominated by hepatocytes (65% of cells), where FlashDeconv predicts accurately (JSD contribution 0.0002). The largest JSD contributions come from T cells (0.0137) and B cells (0.0096)—small populations predicted at near-zero abundance—rather than the endothelial types. AUPR, by contrast, specifically tests the spatial localization of endothelial cells in their respective zones, directly exposing the collinearity problem that JSD is insensitive to.

This discrepancy is not a limitation specific to FlashDeconv's design choices (e.g., the Laplacian or sketching). It reflects a general identifiability challenge for linear deconvolution when reference profiles satisfy $\cos\theta > 0.97$: accurate total abundance estimation is possible (the regression correctly partitions the *total* endothelial signal), but per-spot attribution among near-degenerate subtypes is inherently underdetermined. Methods solving $Y \approx WX$ with $W \geq 0$ under the matrix orientation used here face the same constraint. For a discussion of why per-spot Pearson and RMSE are not computable for this benchmark, see Supplementary Note .

#### Supplementary Note 12: Comparison with marker gene scoring on simulated multi-cell spots

Recent work has shown that marker gene scoring—computing enrichment scores for cell-type-specific gene sets—can achieve competitive rare cell type detection in certain evaluation settings [11]. We systematically compared FlashDeconv against marker gene scoring on all six Spotless Silver Standard tissues (76 cell types total) to determine when compositional deconvolution provides value over independent scoring.

**Marker scoring implementation.** We evaluated two marker gene selection strategies to ensure robustness of our conclusions:

1. **Max-gap selection**: For each cell type, select the top 50 genes ranked by the max-minus-second-max expression gap on the aggregated reference signature matrix (deterministic, $O(K \cdot G)$).
2. **Wilcoxon selection**: For each cell type, perform a one-vs-rest Wilcoxon rank-sum test on log-CPM-normalized individual reference cells and select the top 50 significantly upregulated genes, following the standard Scanpy `sc.tl.rank_genes_groups` and Pritykin et al. protocol.

For both methods, each spot was scored as the mean log-CPM expression of the marker gene set minus the mean expression of a size-matched control gene set (non-marker genes sampled randomly), following the Scanpy `sc.tl.score_genes` protocol. Negative scores were clipped to zero and normalized to sum to 1 per spot to produce pseudo-proportions for fair comparison. The two marker sets overlap modestly (mean 14–27 of 50 genes per type across tissues), confirming that they capture partially distinct aspects of cell type identity.

**Results.**

| | | Pearson $r$ | | | AUPRC | | |
|---|---|---|---|---|---|---|---|
| **Category** | $n$ | **FlashDeconv** | **Scoring (max-gap)** | **Scoring (Wilcoxon)** | **FlashDeconv** | **Scoring (max-gap)** | **Scoring (Wilcoxon)** |
| Rare (<5%) | 26 | **0.902** | 0.724 | 0.574 | **0.919** | 0.792 | 0.647 |
| Moderate (5–15%) | 42 | **0.948** | 0.885 | 0.788 | **0.977** | 0.931 | 0.848 |
| Abundant (>15%) | 8 | **0.995** | 0.978 | 0.945 | **1.000** | 0.999 | 0.998 |
| All | 76 | **0.932** | 0.834 | 0.731 | **0.957** | 0.882 | 0.795 |

In this multi-cell Spotless benchmark, FlashDeconv achieves higher Pearson correlation and AUPRC than marker scoring across all abundance categories under *both* marker selection strategies. The advantage is largest for rare cell types: +0.178 Pearson and +0.127 AUPRC over max-gap scoring, and +0.328 Pearson and +0.272 AUPRC over Wilcoxon scoring. Notably, max-gap selection achieves higher accuracy than Wilcoxon selection for the scoring baseline itself (all-type Pearson 0.834 vs. 0.731), indicating that the aggregated-signature gap criterion—which directly optimizes discriminability between cell type means—is better suited to this multi-cell spot deconvolution task than single-cell-level DE testing. The conclusions are robust to marker selection method: FlashDeconv achieves higher rare-type Pearson in 5 of 6 tissues and higher rare-type AUPRC in 5 of 6 tissues under both strategies.

**Why deconvolution can outperform scoring on multi-cell spots.** The performance gap reflects a structural difference between the two approaches. Marker scoring evaluates each cell type independently: the score for type $k$ depends only on its own marker genes and is unaffected by the abundance of other types. This independence assumption is reasonable when each measurement unit contains predominantly one cell type (single-cell resolution), but breaks down in multi-cell spots where multiple types contribute to the same expression profile. In this regime, marker genes for one type may be expressed at moderate levels due to the presence of a second type that shares some of those genes, generating false-positive scores.

Compositional deconvolution jointly estimates all cell type fractions subject to the constraint $\sum_k w_k = 1$ (after normalization). This compositional constraint forces the solver to adjudicate competing explanations: if a gene is highly expressed, the solver must decide which cell type(s) are responsible, rather than crediting all types whose markers happen to be expressed. For cell types with overlapping signatures (e.g., the kidney CD.Trans and DCT types, where both marker scoring methods achieve Pearson $r < 0.13$ vs. FlashDeconv's 0.93), this joint estimation is critical.

**Scalability.** Even without bootstrap significance testing, marker scoring requires evaluation of $K$ gene sets per spot, each involving ~50 marker genes and ~50 control genes. For atlas-scale datasets, the computational cost is comparable to FlashDeconv's single-pass inference. However, bootstrap-based significance testing ($B = 10{,}000$ resamples as in Pritykin et al.) scales as $O(B \cdot N \cdot K \cdot G_{\text{marker}})$, which becomes prohibitive at million-spot scale: extrapolating from our benchmarks, testing 1,000,000 spots with $K = 16$ cell types and $B = 10{,}000$ would require $\sim 10^{10}$ scoring operations versus FlashDeconv's $O(N \cdot d \cdot K)$ single-pass inference completing in 198 seconds.

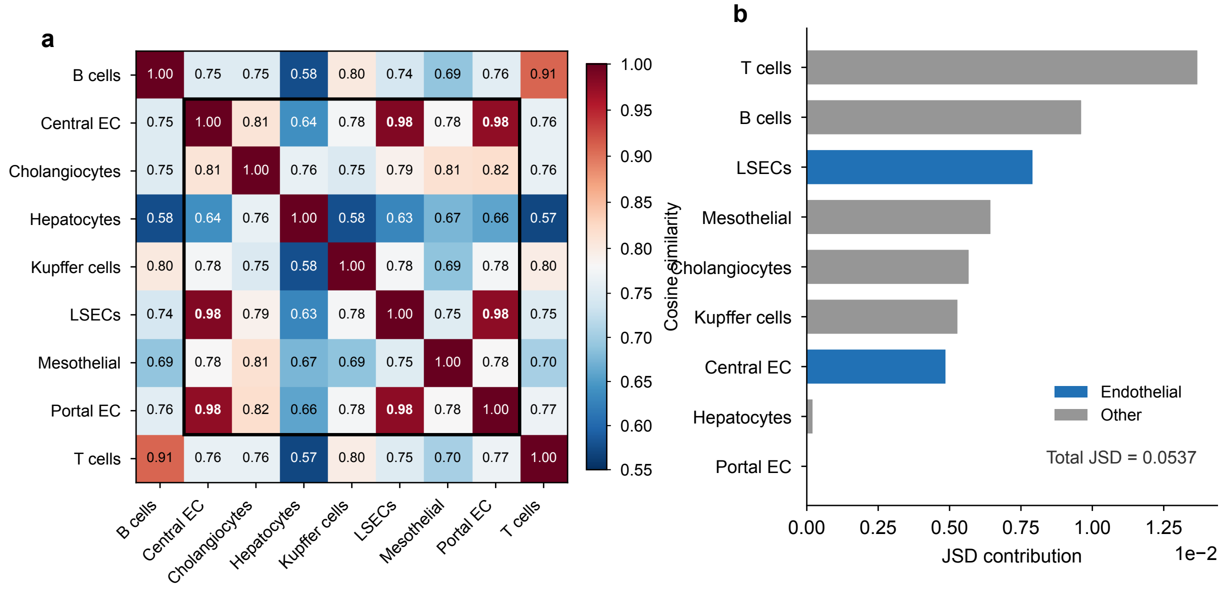


Figure S28: **Per-cell-type decomposition of liver benchmark metrics reveals endothelial signature collinearity as the source of the JSD–AUPR discrepancy. (a)** Pairwise cosine similarity between the 9 cell type signatures in the liver snRNA-seq reference. The three endothelial populations—portal vein EC, central vein EC, and LSECs—form a nearly degenerate triplet (all pairwise similarities >0.975), while other cell type pairs show substantially lower overlap. **(b)** Per-cell-type JSD contributions (averaged across 4 liver Visium samples). The largest contributions come from T cells and B cells (predicted at near-zero abundance), while portal vein EC contributes essentially zero JSD despite poor spatial localization (AUPR = 0.412). Bars colored by endothelial (blue) vs. non-endothelial (gray) status.

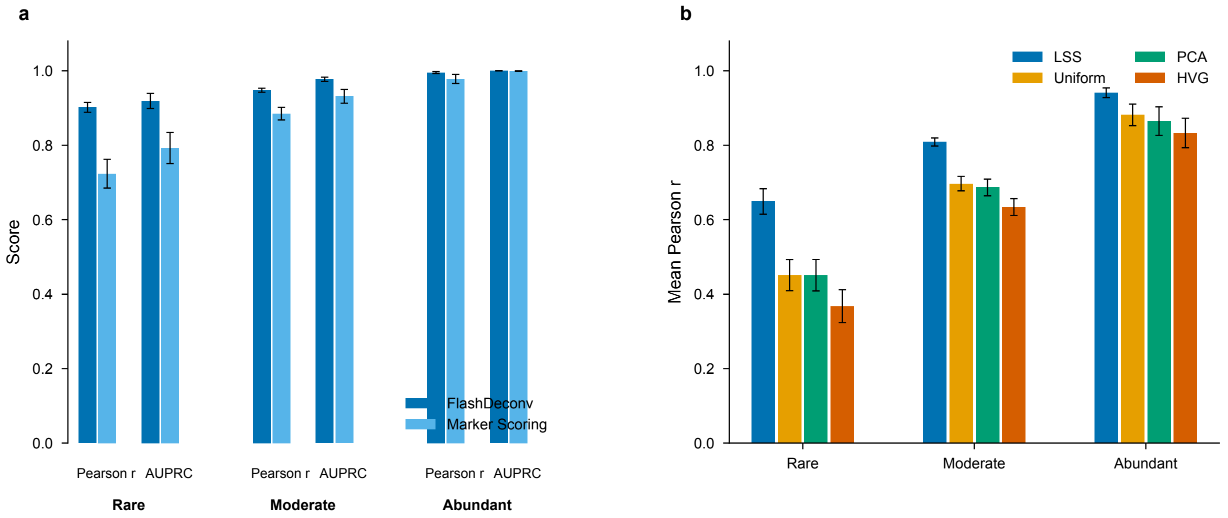


Figure S29: **In the multi-cell Spotless benchmark, FlashDeconv achieves higher accuracy than marker gene scoring across all abundance categories, and leverage weighting gives the best tested trade-off among sketching strategies. (a)** Comparison of FlashDeconv vs. marker gene scoring on 76 cell types across 6 Spotless Silver Standard tissues. Results shown use max-gap marker selection (top-50 markers per type); a sensitivity analysis with Wilcoxon rank-sum markers confirms the same conclusion with an even larger advantage for FlashDeconv (see Supplementary Note  for both methods). FlashDeconv achieves substantially higher Pearson correlation and AUPRC, with the largest advantage for rare cell types (+0.178 Pearson, +0.127 AUPRC vs. max-gap scoring; +0.328 Pearson, +0.272 AUPRC vs. Wilcoxon scoring). **(b)** Comparison of four sketching strategies—leverage-weighted CountSketch (LSS, FlashDeconv default), uniform CountSketch, PCA, and HVG selection—at $d = 512$ across the same 76 cell types. LSS achieves the highest accuracy for all abundance categories under this benchmark, supporting leverage weighting as the best tested trade-off among these alternatives.

#### Supplementary Note 13: Metric availability across benchmark types

The Spotless benchmark suite evaluates deconvolution methods using different metric subsets for different benchmark types, reflecting the distinct ground-truth structures available for each [12]. We report all metrics that are computable for each benchmark type and explain the structural reasons for unavailability where applicable.

**Silver Standard (54 synthetic datasets).** Synthetic "pseudo-spots" are generated by computationally mixing single-cell transcriptomes with known per-spot cell type proportions. Because ground-truth proportions are available at the per-spot level, all metrics are computable: Pearson correlation, RMSE, and JSD (aggregate accuracy), AUPR (rare cell type detection), and per-cell-type AUPRC, precision, recall, and F1 (per-class performance). Supplementary Table S2 reports aggregate metrics; per-cell-type metrics for all 13 methods across 684 cell-type instances are provided in Supplementary Data .

**Gold Standard (seqFISH+ and STARMap).** Real spatial transcriptomics data with ground-truth composition derived from co-registered single-molecule FISH imaging. Per-spot proportions are available, enabling all metrics. However, the extremely small sample sizes (7–9 spots per seqFISH+ FOV, 108 spots for STARMap) limit statistical power and produce high variance across FOVs.

**Liver case study (4 Visium slides).** Ground truth consists of tissue-level cell type proportions estimated from matched snRNA-seq, not per-spot annotations. JSD is computed by comparing predicted tissue-level composition against snRNA-seq proportions. AUPR evaluates whether portal and central vein endothelial cells are spatially localized to their respective zones (a binary spatial pattern test). *Per-spot Pearson correlation and RMSE are not computable* because no per-spot ground-truth proportions exist—the snRNA-seq reference provides a single proportion vector for the entire tissue section.

**Melanoma case study (3 Visium slides).** Ground truth consists of tissue-level proportions from Molecular Cartography. JSD is computed against these tissue-level proportions. Unlike the liver dataset, the melanoma tissue lacks distinct spatial zonation patterns (e.g., periportal vs. pericentral zones) that would enable AUPR evaluation. *Only JSD is evaluable* for this benchmark.

This metric availability structure follows the Spotless benchmark design (Sang-aram et al., Fig. 1b, eLife 2024) and is not a consequence of our evaluation choices. We report the maximum set of evaluable metrics for each benchmark type.

Table S2: **Unified benchmark metrics across all evaluation scenarios.** Aggregate performance of FlashDeconv and the top four competing methods. Metrics marked "—" are structurally unavailable for that benchmark type (see Supplementary Note ). Silver Standard values are means across 54 datasets; Gold Standard values are means across FOVs; liver and melanoma are means across slides. Per-cell-type metrics (AUPRC, F1) are reported as mean across cell-type instances stratified by abundance; ranges span rare (<5%) to abundant (>15%) categories.

| **Benchmark** | **Method** | **Pearson** | **RMSE** | **JSD** | **AUPR** | **AUPRC[a]** | **F1[a]** |
|---|---|---|---|---|---|---|---|
| Silver Standard ($n = 54$ datasets) | FlashDeconv | **0.944** | **0.053** | **0.047** | **0.957** | 0.87–1.00 | 0.61–0.90 |
| | RCTD | 0.934 | 0.055 | 0.050 | 0.951 | 0.91–1.00 | 0.69–0.91 |
| | Cell2Location | 0.918 | 0.057 | 0.061 | 0.921 | 0.88–1.00 | 0.72–0.94 |
| | SpatialDWLS | 0.897 | 0.068 | 0.067 | 0.907 | 0.84–0.94 | 0.74–0.90 |
| | NNLS | 0.850 | 0.081 | 0.118 | 0.819 | 0.62–0.98 | 0.44–0.82 |
| Gold: seqFISH+ cortex (7 FOVs) | Cell2Location | **0.285** | 0.306 | 0.295 | **0.531** | — | — |
| | SpatialDWLS | 0.274 | 0.238 | **0.270** | 0.526 | — | — |
| | FlashDeconv | 0.190 | **0.108** | 0.361 | 0.452 | — | — |
| Gold: seqFISH+ OB (7 FOVs) | STRIDE | **0.803** | **0.098** | **0.083** | **0.873** | — | — |
| | DSTG | 0.780 | 0.111 | 0.091 | 0.868 | — | — |
| | FlashDeconv | 0.608 | 0.149 | 0.171 | 0.806 | — | — |
| Gold: STARMap (108 spots) | SpatialDWLS | **0.712** | **0.129** | 0.307 | 0.654 | — | — |
| | RCTD | 0.679 | 0.150 | 0.413 | 0.657 | — | — |
| | FlashDeconv | 0.665 | 0.132 | **0.252** | **0.661** | — | — |
| Liver (4 slides) | RCTD | — | — | **0.033** | 0.89 | — | — |
| | Cell2Location | — | — | 0.035 | **0.94** | — | — |
| | FlashDeconv | — | — | 0.056 | 0.66 | — | — |
| Melanoma (3 slides) | Cell2Location | — | — | **0.013** | — | — | — |
| | SPOTlight | — | — | 0.014 | — | — | — |
| | FlashDeconv | — | — | 0.015 | — | — | — |

[a]Per-cell-type metrics: range spans rare (<5% abundance) to abundant (>15%) categories. Gold standard per-cell-type metrics not reported due to insufficient sample sizes per FOV. See Supplementary Data  for complete per-cell-type results on Silver Standard.

#### Supplementary Data

**Supplementary Data 1: Complete benchmark results (Excel file).** This file contains comprehensive performance metrics for FlashDeconv and 12 competing methods across all benchmark datasets from the Spotless study [12]. The Excel workbook includes eight sheets:

- **S1a_Silver_13methods**: Method comparison on 54 Silver Standard synthetic datasets (Pearson correlation, RMSE, JSD, AUPR)
- **S1b_Liver_13methods**: Complete rankings on the liver Visium case study (JSD, AUPR)
- **S1c_Melanoma_13methods**: Complete rankings on the melanoma Visium case study (JSD)
- **S1d_seqFISH_13methods**: Gold standard seqFISH+ results (Pearson, RMSE, JSD, AUPR)
- **S1e_STARMap_13methods**: Gold standard STARMap results (Pearson, RMSE, JSD, AUPR)
- **S1f_Summary**: Summary of FlashDeconv's strengths and limitations across benchmarks
- **S1g_Per_celltype_Silver**: Per-cell-type Pearson, AUPRC, precision, recall, and F1 for all 13 methods across 684 cell-type instances (76 cell types $\times$ 9 abundance patterns $\times$ 6 tissues, stratified by abundance category)
- **S1h_Unified_all_benchmarks**: Aggregate metrics (Pearson, RMSE, JSD, AUPR) for all 13 methods across all benchmark types

All competing method results are sourced directly from the Spotless benchmark (Zenodo: `https://zenodo.org/records/10277187`), ensuring fair comparison using identical evaluation protocols.
